# Multifaceted roles of cohesin in regulating transcriptional loops

**DOI:** 10.1101/2024.03.25.586715

**Authors:** Minji Kim, Ping Wang, Patricia A. Clow, I (Eli) Chien, Xiaotao Wang, Jianhao Peng, Haoxi Chai, Xiyuan Liu, Byoungkoo Lee, Chew Yee Ngan, Feng Yue, Olgica Milenkovic, Jeffrey H. Chuang, Chia-Lin Wei, Rafael Casellas, Albert W. Cheng, Yijun Ruan

## Abstract

Cohesin is required for chromatin loop formation. However, its precise role in regulating gene transcription remains largely unknown. We investigated the relationship between cohesin and RNA Polymerase II (RNAPII) using single-molecule mapping and live-cell imaging methods in human cells. Cohesin-mediated transcriptional loops were highly correlated with those of RNAPII and followed the direction of gene transcription. Depleting RAD21, a subunit of cohesin, resulted in the loss of long-range (>100 kb) loops between distal (super-)enhancers and promoters of cell-type-specific genes. By contrast, the short-range (<50 kb) loops were insensitive to RAD21 depletion and connected genes that are mostly housekeeping. This result explains why only a small fraction of genes are affected by the loss of long-range chromatin interactions due to cohesin depletion. Remarkably, RAD21 depletion appeared to up-regulate genes located in early initiation zones (EIZ) of DNA replication, and the EIZ signals were amplified drastically without RAD21. Our results revealed new mechanistic insights of cohesin’s multifaceted roles in establishing transcriptional loops, preserving long-range chromatin interactions for cell-specific genes, and maintaining timely order of DNA replication.

## INTRODUCTION

The human genome is organized within the three-dimensional (3D) nuclear space as large-scale chromosomal territories, which encompass chromatin folding compartments or domains (Lieberman-Aiden et al., 2009; Dixon et al., 2012) and individual chromatin loops mediated by protein factors (Fullwood et al., 2009; Li et al., 2012; Tang et al., 2015; Kim et al., 2016; Weintraub et al., 2017; Mumbach et al., 2017; Grubert et al., 2020; Dejosez et al., 2023). Each of these components has potential functional implications in nuclear processes. Notably, chromatin loops—the basic unit of chromosome folding—largely define the chromosome folding architectures and their underlying functions (Phillips-Cremins et al., 2013; Braccioli and de Wit, 2019; Davidson et al., 2023). It is known that CTCF, a zinc finger protein, binds specific DNA motifs to block the ring-like extruder cohesin and define the boundaries of a complete chromatin loop with a pair of convergent CTCF binding motifs (Rao et al., 2014; Tang et al., 2015). Genetic perturbation of CTCF sites alters genome organization and has implications in cancer and genetic disorders through direct or indirect effects on gene transcription (Ushiki et al., 2021; Kubo et al., 2021), further highlighting the key role of genome architecture in cellular functions. In addition to CTCF/cohesin-based loops, there is a class of chromatin loops that involve transcription regulatory elements (i.e., transcriptional loop). For example, enhancer-promoter (E-P) and promoter-promoter (P-P) loops associated with RNA Polymerase II (RNAPII) and transcription factors such as oestrogen-receptor-α (ERα) and YY1 appear to modulate transcriptional activity (Fullwood et al., 2009; Li et al., 2012; Weintraub et al., 2017). Therefore, it is important to comprehensively characterize the molecular mechanisms of chromatin loop formation, and to test how such features impact genome function.

A pressing unresolved question in 3D genomics is the precise causal relationship between genome structure and function, particularly gene transcription. Acute depletion of cohesin, CTCF, YY1, or a cohesin unloading factor WAPL resulted in a range of alterations in genome topology, yet these changes appear to have minimal impact on gene expression (Rao et al., 2017; Nora et al., 2017; Haarhuis et al., 2017; Liu et al., 2021; Hsieh et al., 2022). This discrepancy raises the interesting question of whether cohesin-mediated chromatin topology is relevant to gene transcription and if any cohesin-independent mechanisms exist that regulate transcription, perhaps through interactions that are not detectable by Hi-C experiments due to resolution or specificity (Goel et al., 2023). The converse experimental scenario is also unsettled: a rapid depletion of RNA Polymerases eliminated transcription but resulted in only subtle or no changes to the overall landscape of chromatin folding domains as demonstrated in mouse embryonic stem cells (Jiang et al., 2020) and in human cells (El Khattabi et al., 2019). Although cohesin has been proposed to activate gene expression by tethering cognate promoters and enhancers (Grubert et al., 2020) and has been tested at the *Shh* locus (Kane et al., 2022), the genome-wide causal impact of cohesin-mediated loop extrusion on the formation of these enhancer-promoter contacts and subsequent gene expression remains largely unresolved. Also debatable is whether RNAPII pushes cohesin, or vice versa, or both (Busslinger et al., 2017; Thiecke et al., 2020; Banigan et al., 2023). Moreover, the extent to which multiple enhancers regulate a single gene through “transcription hubs” (Iborra et al., 1996; Li et al., 2012; Sabari et al., 2018) is not yet fully characterized in the mammalian genome.

These gaps in knowledge are partially due to technical limitations. Despite recent breakthroughs in imaging technologies (Bintu et al., 2018; Wang et al., 2019; Gabriele et al., 2022), it remains challenging to visualize genome folding changes over time *in vivo* with high resolution, throughput, and specificity. High-throughput sequencing based techniques Hi-C (Lieberman-Aiden et al., 2009) and related methods including Micro-C (Hsieh et al., 2015) provide genome-wide unbiased chromatin folding maps, yet often lacks the specificity and has high background noise. ChIA-PET (Fullwood et al., 2009; Tang et al., 2017), and its variants HiChIP (Mumbach et al., 2016) and PLAC-seq (Fang et al., 2016), employed immunoprecipitation (ChIP) to enrich pairwise chromatin interactions mediated by a specific protein, thereby enhancing a functional specificity. Furthermore, ligation-free methods were also developed to capture multiplex interactions based on sample slicing (GAM; Beagrie et al., 2017; Beagrie et al., 2023), split-and-pool barcoding (SPRITE; Quinodoz et al., 2018; Vangala et al., 2020; Arrastia et al., 2022), microfluidic encapsulation (ChIA-Drop; Zheng et al., 2019), and long-read sequencing (Allahyar et al., 2018; Deshpande et al., 2022; Dotson et al., 2022). However, most of these methods were performed without immunoprecipitation and thus could not delineate which transcription factors are responsible for mediating or maintaining certain types of chromatin structures, precluding researchers from quantifying the extent to which—if any—cohesin impacts transcriptional processes genome-wide.

Here, we leverage ChIP-enriched protein-specific ChIA-Drop (Zheng et al., 2019) to investigate cohesin’s involvement in establishing transcriptional multiplex chromatin interactions in conjunction with RNAPII in the human genome. Our data revealed high correlation between cohesin and RNAPII in modulating transcriptional activities. We observed that cohesin at its loading sites largely follows RNAPII in the orientation of transcription, and that cohesin highly correlates with RNAPII at facilitating transcriptional contacts among promoters and distal (super-)enhancers. By depleting RAD21 (a cohesin subunit) and enriching for specific protein factor via in situ ChIA-PET (Wang et al., 2021), we directly tested if there is a causal relationship between cohesin and RNAPII: cohesin is responsible for long-range (>100 kb) interactions involving convergent CTCF sites and enhancer-promoter pairs, but short (<50 kb) promoter-promoter loops are maintained independent of cohesin. Our study suggests that while cohesin participates in all transcriptional loops between E-P and P-P interactions, only the long-range transcriptional loops require cohesin to maintain such topological structure for cell-type-specificity of genes. Finally, up-regulated genes, which were often overlooked and could not be explained by differential chromatin loops (Rao et al., 2017; Hsieh et al., 2022), had distinct DNA replication patterns from publicly available 16-stages Repli-seq data (Emerson et al., 2022): upon depleting RAD21, its replication signal is drastically amplified and disrupted particularly for early stages. These results represent a comprehensive dissection of the relationship between cohesin and RNAPII and the impact of cohesin subunit RAD21 depletion on down-regulated, unchanged, and up-regulated genes.

## RESULTS

### ChIA-Drop data reveal detailed multiplex chromatin interactions mediated by CTCF, cohesin, and RNA Polymerase II

It has been established that RNAPII-associated loops often connect distal cis-regulatory elements to gene promoters for transcription regulation (Li et al., 2012). Although cohesin loading has been related to transcriptionally active regions via NIPBL ChIP-seq (Kagey et al., 2010; Kieffer-Kwon et al., 2013; Zuin et al., 2014; Busslinger et al., 2017; Zhu et al., 2021) and is thought to be involved in transcription (Peric-Hupkes and van Steensel, 2008), it is unclear whether cohesin is actively involved in establishing chromatin interactions among promoters and enhancers, and if so, to what extent.

To unravel the intricate details of the chromatin looping process *in vivo*, we analyzed the human genome in the GM12878 B-lymphoblastoid cell line with ChIA-Drop (Zheng et al., 2019), a single-molecule technique for mapping multiplex chromatin interactions (**Figure 1A**). We added a chromatin immunoprecipitation (ChIP) step to the ChIA-Drop protocol and probed CTCF, cohesin (by targeting its subunits SMC1A and RAD21), and RNA Polymerase II (RNAPII) (**Table S1**) to explore chromatin interactions mediated by proteins directly involved in topology or transcription. DNA sequencing reads were processed using ChIA-DropBox (Tian et al., 2019) and MIA-Sig (Kim et al., 2019) algorithms to identify putative chromatin looping complexes containing multiple chromatin fragments indexed with the same barcode (see **Methods**). Interestingly, with the chromatin immunoprecipitation (ChIP) added in the ChIA-Drop protocol, the proportion of 2-fragment chromatin complexes comprised more than 80% (**Figure S1A**), indicating that two-point contacts in mammalian genomes are largely the basic units of chromatin interactions mediated by specific factors. Although a relatively small proportion (less than 20%), we have captured millions of chromatin folding complexes containing three or more fragments (**Figure S1A**, **Table S1**), reflecting multiplex chromatin interactions in each ChIA-Drop experiments. For example, a 1.5 Mbps region on chromosome 10 encompass three main domains in 2D contact maps of Hi-C, CTCF, cohesin, and RNAPII ChIA-PET (**Figure 1B**, left panel). When zoomed into one domain of 430 kbps size, CTCF ChIA-Drop complexes are enriched at CTCF binding sites shown in light blue. Cohesin ChIA-Drop data additionally have fragments in the non-CTCF site in the middle, which is the enhancer site that RNAPII ChIA-Drop data bind exclusively and connect to other sites (**Figure 1B**, right panels). We verified that the ChIA-Drop data are highly reproducible and comparable to those of the ligation-based ChIA-PET (**Figure S1B**), and next focused on cohesin and RNAPII.

**Figure 1:**
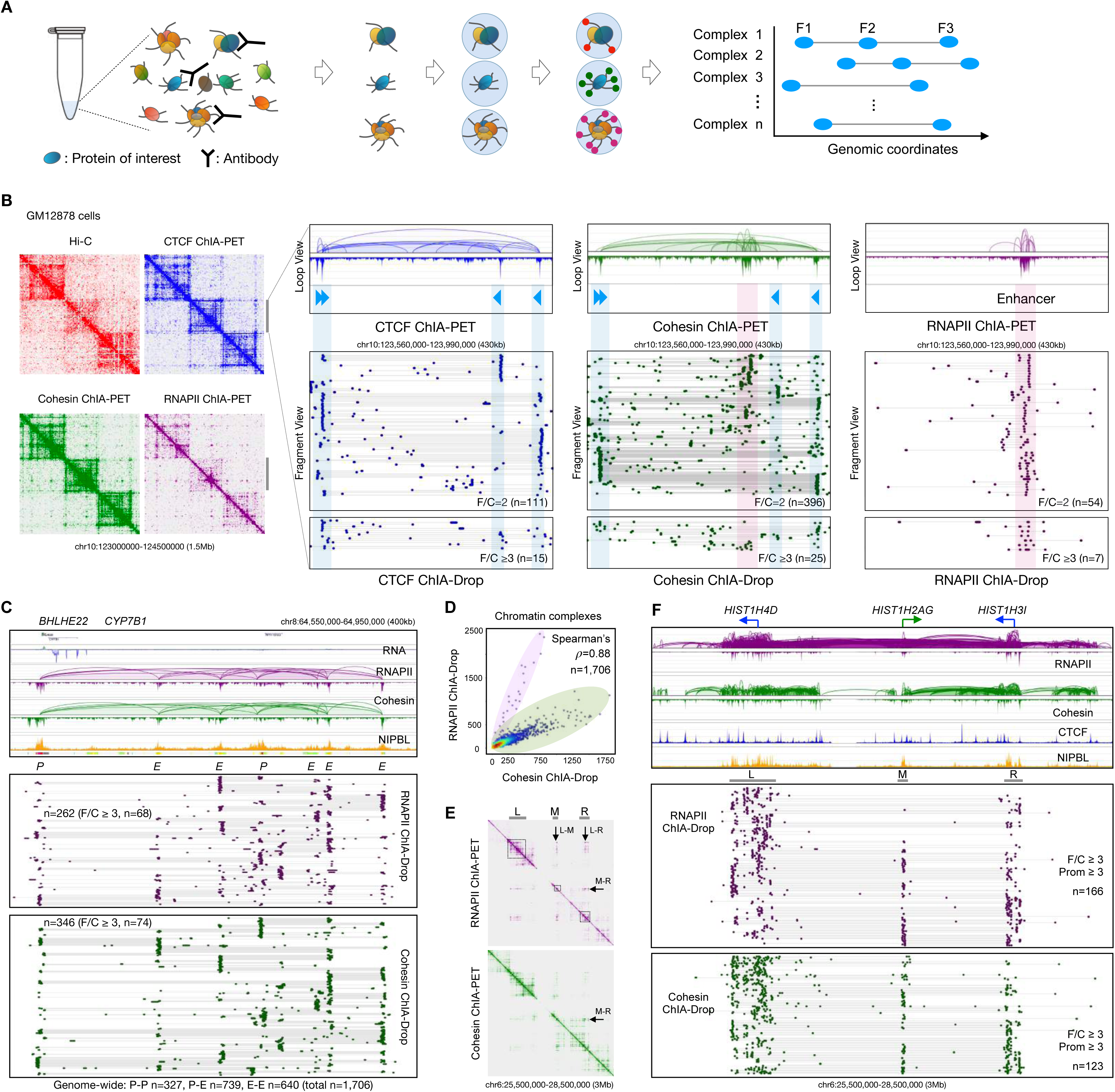
ChIA-Drop data for mapping chromatin interactions mediated by CTCF, cohesin, and RNAPII. **(A)** A brief schematic of ChIA-Drop, which encapsulates ChIP-enriched samples of chromatin complexes into individual droplets with unique barcodes for obtaining single-molecule multiplex chromatin interactions via DNA sequencing and mapping analysis. Each ChIA-Drop complex contains multiple fragments (in blue ovals F1, F2, F3) connected by a straight line. **(B)** 2D contact matrices of Hi-C, CTCF, cohesin, and RNA Polymerase II (RNAPII) ChIA-PET data at a 1.5 Mb region. Corresponding ChIA-PET loops and peaks at a further zoomed-in 430kb region are included as references (top panels). CTCF binding motifs in CTCF and cohesin data tracks are marked with light blue arrows indicating the binding motif orientation. Below in the bottom panels, fragment views show detailed chromatin interactions by ChIA-Drop data, where each row of dots and a connecting line represents a putative chromatin complex with ≥ 2 interacting fragments; pairwise (fragments per complex (F/C) = 2) and multi-way (F/C ≥ 3) interactions are presented separately with the number of complexes in each category denoted as *n*. CTCF-enriched and RNAPII-enriched regions are highlighted in blue and purple, respectively. **(C)** An example of a chromatin domain. Top tracks are RNA-seq, RNAPII and cohesin ChIA-PET loops/peaks, NIPBL ChIP-seq, and ChromHMM states promoters (*P*) and enhancers (*E*). Lower tracks are chromatin fragment views of RNAPII and cohesin ChIA-Drop complexes with two or more (≥ 2) enhancers (*E*) and promoters (*P*) simultaneously connected by RNAPII and cohesin ChIA-Drop complexes, with their numbers recorded as n. **(D)** A scatter plot of RNAPII and cohesin ChIA-Drop chromatin complexes co-localized at 1,706 loci, a subset of which exhibit significantly higher RNAPII counts than cohesin (highlighted in purple) and deviate from the main trajectory of other chromatin loci (highlighted in green). **(E)** The histone gene cluster (*HIST1*) on chromosome 6 is organized into left (L), middle (M), and right (R) regions. The 2D pairwise contact map of RNAPII ChIA-PET data (purple) have inter-region interactions as indicated by arrows for clusters L-M, L-R, and M-R, while the cohesin ChIA-PET data (green) show relatively weak inter-region signals. **(F)** Detailed browser views of the 3 Mb chromatin domain harboring histone gene clusters. Top tracks: RNAPII and cohesin ChIA-PET loops/peaks and CTCF and NIPBL ChIP-seq peaks. Bottom tracks: chromatin fragment views of RNAPII and cohesin ChIA-Drop data with more than 2 fragments per complex (F/C≥3). The number of multiplex chromatin interactions among individual histone genes are provided as n. See also **Figure S1**.

### Cohesin and RNAPII are correlated in connecting multiple enhancers and promoters

We observed many chromatin loops between enhancer and promoter loci (Tang et al., 2015) that are also cohesin loading (NIPBL binding) sites (**Figure 1C**, top panel) in RNAPII and cohesin ChIA-PET data. In total, we identified 1,706 high quality transcriptional loops (see **Methods**), including 327 promoter-promoter, 739 promoter-enhancer, and 640 enhancer-enhancer loops from ChIA-PET datasets; these loops serve as a reference set for the downstream ChIA-Drop analyses. Most of these loops were typically inter-connected in “daisy-chain” patterns (Li et al., 2012) and were also captured by RNAPII and cohesin ChIA-Drop as multiplex chromatin interaction complexes connecting 2 or more regulatory elements in a single-molecule resolution (**Figure 1C**, bottom panel). This region had higher proportion of chromatin complexes containing 3 or more fragments (26% in RNAPII, 21% in cohesin) than genome-wide distributions (13.5% in RNAPII, 13.4% in cohesin; **Figure S1A**).

Importantly, we noticed that cohesin and RNAPII ChIA-Drop complexes are highly correlated (**Figure 1D**), particularly when using CTCF ChIA-Drop data as a background reference (**Figure S1C**; see **Methods**). While most of the transcriptional loops (data points highlighted in green in the scatter plot) have approximately equal numbers of chromatin complexes represented by both RNAPII and cohesin ChIA-Drop data, a small group of 21 loops stands out with a higher number of RNAPII ChIA-Drop complexes than cohesin (data points highlighted in purple in the scatter plot) (**Figure 1D**). Interestingly, the latter group falls into a 2 Mb region on chromosome 6, which harbors histone (*HIST1*) gene clusters *HIST1H4D*, *HIST1H2AG*, *HIST1H3I*, and others. Interactions among the three large domains ‘L’ (left), ‘M’ (middle), and ‘R’ (right) are more pronounced in RNAPII than cohesin as shown by 2D contact maps (**Figure 1E**). Notably, both RNAPII and cohesin ChIA-Drop data exhibit a high degree of multiplexity in this 2 Mb region: more than a hundred chromatin complexes (n=166 for RNAPII, n=123 for cohesin) connect 3 or more gene promoters, with some connecting as many as 15 (**Figure 1F**). RNAPII generally connected higher number of promoters and enhancer than expected (**Figure S1D**, see **Methods**). Thus, our ChIA-Drop data provide concrete evidence that cohesin—along with RNAPII—connects cis-regulatory elements, and with high degree of multiplexity as a potential topological mechanism to regulate gene expressions including the histone gene clusters. A potential implication of these clusters of multiway chromatin interactions is the notion of co-transcription regulation for multiple genes, as previously demonstrated for pairs of genes (Li et al., 2012).

### Cohesin coordinates with RNAPII in the direction of gene transcription

Several studies hinted that cohesin may facilitate transcriptional activity by genome-wide correlation or by testing at selected few loci (Schaaf et al., 2013; Mannini et al., 2015; Busslinger et al., 2017; Thiecke et al., 2020; Grubert et al., 2020; Kane et al., 2022), but the direct genome-wide evidence of cohesin’s role—if any—in gene regulation is lacking. We sought to comprehensively characterize the relationship between RNAPII and cohesin in transcriptional regulation in the context of ongoing chromatin looping events.

We first examined how RNAPII behaves in association with chromatin loops at CTCF binding motifs, i.e., cohesin anchoring sites that also overlap with transcription start sites (TSS). Aggregating the RNAPII ChIA-Drop contacts at promoters of active genes (n=264; see **Methods**) whose transcription direction is in concordance with the orientation of CTCF motifs revealed that RNAPII-associated chromatin complexes follow the CTCF motif orientation, and so do cohesin-associated chromatin complexes (**Figure 2A**). As exemplified at the *PIEZO2* locus (**Figure 2B**, top panel), the gene promoter (TSS) is upstream of a CTCF binding peak with a motif in an orientation that is concordant with the direction of transcription. Accordingly, RNAPII and cohesin ChIA-Drop complexes that are in concordant direction are 4 to 9 times more frequent than those in discordant direction (107 vs. 12 for RNAPII and 120 vs. 28 for cohesin), with a clear bias towards the left orientation in the 2D contact maps of ChIA-PET and Hi-C data including all pairwise contacts (**Figure 2B**). A similar pattern is also observed for *LRCH1* (**Figure S2A**). However, when CTCF motif and transcription of active genes are in opposite orientations (n=199; see **Methods**), the RNAPII and cohesin ChIA-Drop data showed balanced signals in both directions (**Figure 2A**). For example, at the *LPAR1* locus (**Figure S2B**), there were similar number of chromatin complexes extending to the left and to the right for both the RNAPII and cohesin ChIA-Drop data (31 vs. 27 for RNAPII and 95 vs. 84 for cohesin), implying that the direction of RNAPII-associated transcription may be interfered by CTCF: some proportion of the cohesin may move in the direction of transcription while others move along the CTCF motif orientation when TSS and CTCF/cohesin anchoring sites are co-localized but in opposite orientations.

**Figure 2:**
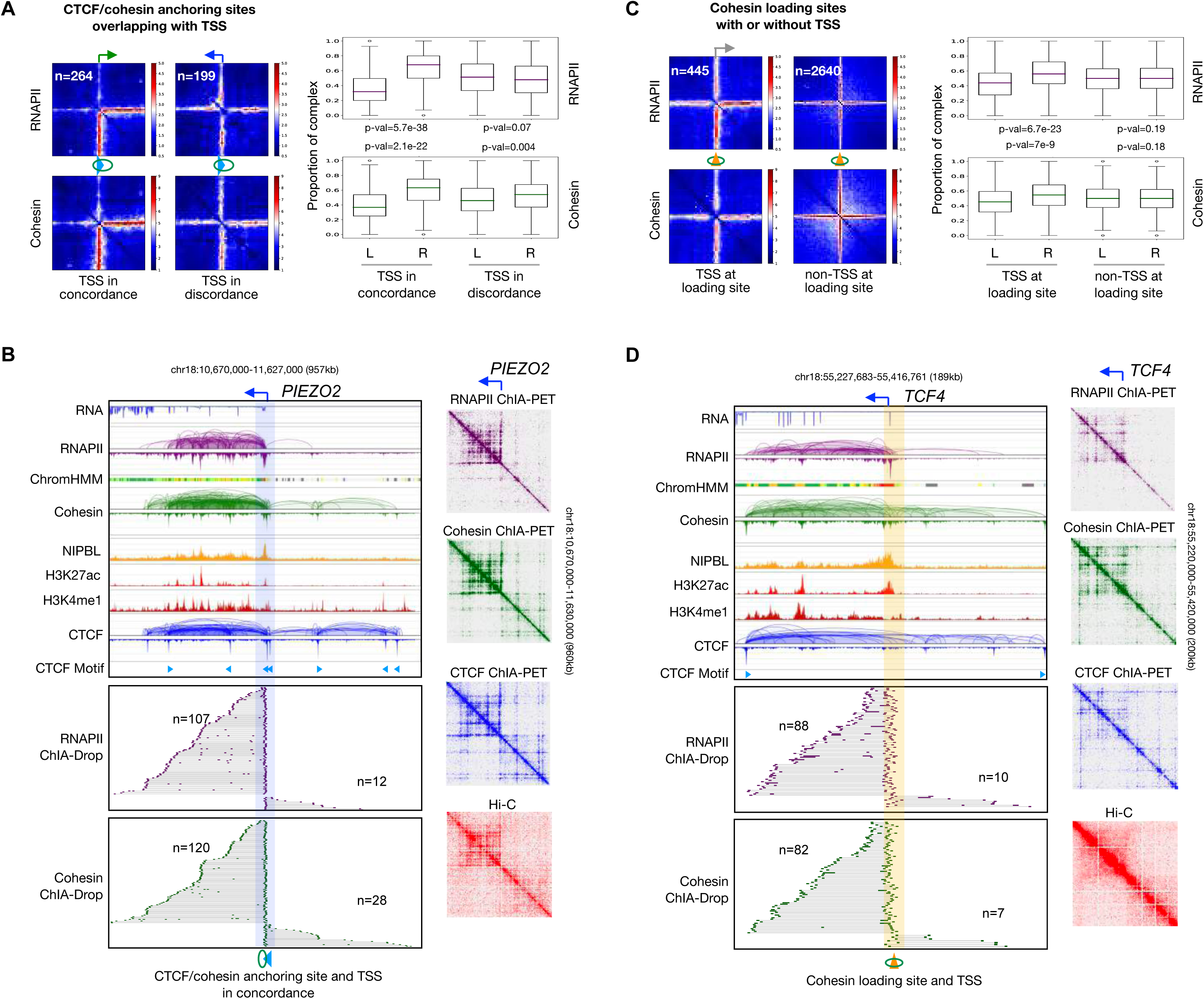
Transcriptional loops mediated by RNAPII and cohesin through concerted efforts. **(A)** Left panel: 2D pairwise contact aggregation maps of chromatin loop anchoring sites (CTCF-cohesin, blue arrow with green circle) co-localized with gene transcription start site (TSS) in RNAPII and cohesin ChIA-Drop data. TSS (green right-angled arrow, rightward) is in concordance with the direction of CTCF binding motif (blue arrow, rightward), or TSS (blue right-angled arrow, leftward) is in discordance with CTCF motif direction (blue arrow, rightward). Right panel: boxplots of RNAPII and cohesin ChIA-Drop complexes corresponding to the left panel. Proportions of chromatin contacts from the CTCF binding motif leftward (L) and rightward (R) with TSS in concordance (left) and in discordance (right) are calculated. All p-values are from the two-sided Mann-Whitney test. **(B)** An example of RNAPII-associated chromatin loops at TSS (right-angled arrow) co-localized with CTCF/cohesin anchoring site (highlighted in blue) and in concordance with CTCF binding motif at the *PIEZO2* locus. Top tracks are RNA-seq, RNAPII ChIA-PET, chromatin states (ChromHMM; enhancers in yellow, promoters in red), cohesin ChIA-PET, NIPBL ChIP-seq, H3K27ac ChIP-seq, H3K4me1 ChIP-seq, CTCF ChIA-PET, and CTCF binding motif (directional blue arrows). Middle and lower tracks are RNAPII and cohesin ChIA-Drop complexes extending from CTCF/cohesin anchoring site in each direction, with n denoting their numbers. On the right are the 2D contact maps of RNAPII ChIA-PET, cohesin ChIA- PET, CTCF ChIA-PET, and Hi-C data encompassing all possible interactions. **(C)** Same as in panel **A** but for RNAPII binding at cohesin loading sites (yellow cone with circle, non-directional), which also coincide with TSS (grey right-angled arrow). **(D)** Similar to panel **B** but at TSS co-localized with NIPBL binding/cohesin loading site (highlighted in yellow) at the *TCF4* locus. See also **Figure S2**.

Next, at the cohesin loading sites (NIPBL binding without CTCF binding; see **Methods**) that coincide with the transcription start sites (TSS) of actively transcribed genes, we noticed that RNAPII and cohesin ChIA-Drop complexes displayed significant bias towards the direction of transcription genome-wide (**Figure 2C**) as exemplified at the loci of *TCF4* (**Figure 2D**) and *HIVEP1* (**Figure S2C**). This biased pattern of RNAPII ChIA-Drop data is akin to the one-sided reeling model by RNAPII in transcription that we demonstrated in *Drosophila* (Zheng et. al., 2019) and reflects a RNAPII-mediated action of transcriptional looping in human cells. Considering that cohesin on its own has no directional preference at its loading sites (**Figure 2C, S2D**), the specific biased pattern of cohesin ChIA-Drop data here implies that RNAPII may guide cohesin to move along in the direction of transcription.

### Cohesin- and RNAPII-associated chromatin loops connect super-enhancer to target gene promoter

Super-enhancers (SEs), large clusters of gene regulatory elements, play a key role in transcriptional regulation in mammalian cells (Hnisz et al., 2013). They primarily modulate the expression of cell-type-specific genes by strengthening long-range contacts between super-enhancer and cognate promoters (SE-P) (Dowen et al., 2014). However, it is unclear whether cohesin is essential for mediating all or a subset of these interactions; also unknown is the precise additive role of constitutive enhancer elements within SE (Dukler et al., 2016).

As expected, we observed extensive 2D contact signals in RNAPII ChIA-Drop data inside SEs compared to the random control, implying abundant E-E interactions among the constitutive enhancer elements within a SE unit (**Figure S3A**). Interestingly, the profiles of RNAPII and cohesin ChIA-Drop data within SEs were similar, suggesting that cohesin may also be extensively involved in SE-associated transcription regulation. By contrast, the CTCF ChIA-Drop data did not have enriched signals within SEs, indicating that CTCF’s involvement in SE-P interactions, if any, is different from that of cohesin and RNAPII. To investigate the details of SE-P conformation, we systematically analyzed RNAPII and cohesin ChIA-PET and ChIA-Drop datasets for the involvement of these two factors. Of the 257 known SEs described in GM12878 cells (Hnisz et al., 2013), we identified the connectivity in 188 SEs and their target genes by incorporating both RNAPII ChIA-PET and ChIA-Drop data (see **Methods**). One of the target genes is *MYC*, which encodes a master regulator in cells and requires SEs for expression (Schuijers et al., 2018). Although ChIA-PET data showed extensive “daisy-chain” interactions in aggregated maps, it could not uncover chromatin complexes engaging more than 2 (≥3) regulatory elements (**Figure 3A**, top left panel). Thus, we examined ChIA-Drop data.

**Figure 3:**
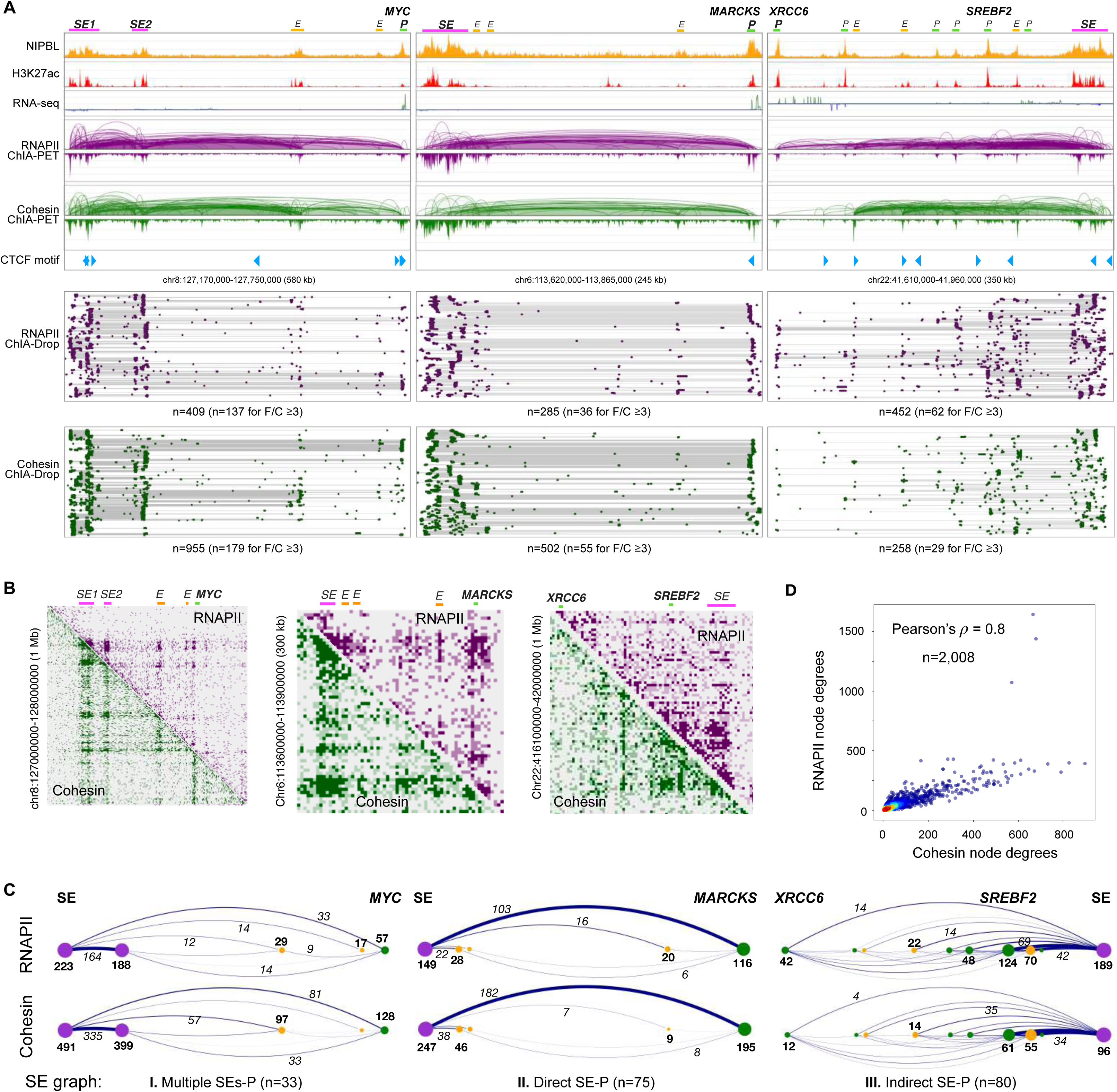
Multiplex transcriptional chromatin interactions involving super-enhancers. **(A)** Three examples of chromatin interaction paths from super-enhancers (*SE*) to target gene promoters (*P*): left panel, multiple *SE*s connecting to *MYC* promoter; middle panel, direct connections from SE to *MARCKS* promoter; right panel, indirect connections through many intermediate *Es* and *Ps* from SE to connect to *XRCC6* promoter. Top tracks: ChIP-seq of NIPBL and H3K27ac, RNA-seq, ChIA-PET loops/peaks of RNAPII (purple) and cohesin (green), and CTCF binding motifs. Bottom tracks: fragment view of multiplex chromatin complexes in RNAPII (purple) and in cohesin (green) ChIA-Drop data, where n is the number of chromatin complexes, a subset of which have more than 2 fragments per complex (F/C ³3). **(B)** The 2D contact maps of RNAPII and cohesin ChIA-Drop data at three exemplary regions, each including *MYC*, *MARCKS*, *XRCC6* genes. **(C)** Graph representations for the three categories of SE-P chromatin interaction patterns in RNAPII (top panel) and cohesin (bottom panel) ChIA-Drop data corresponding to the three examples of SE-P in panel **A**. The nodes in SE-P graph are SE in purple, enhancer E in yellow, and promoter P in green dots. The node degrees are reflected by the dot sizes and indicated by bold numbers, while the edge weights (number of complexes connecting the two nodes) are reflected in the line thickness and recorded as italic numbers. The three types of SE-P graphs in GM12878 cells are: I) Multiple SEs-P involving two or more SEs together connecting to the target gene promoter (n=33); II) Direct SE-P where SE directly connect to the target gene promoter (n=75); III) Indirect SE-P, where SE through multiple intermediate elements in a series of cascade connections indirectly connect to the target gene promoter (n=80). **(D)** A scatterplot between node degrees (see **Methods**) of RNAPII and cohesin ChIA-Drop complexes. See also **Figure S3**.

From 2D contact maps, both RNAPII and cohesin ChIA-Drop data showed strong interaction signals involving two SEs and two intermediate enhancers connecting to *MYC* as indicated (**Figure 3B**, left panel). We next sought to visualize the detailed multiplex interactions therein. When zoomed into the SE-*MYC* domain by both RNAPII and cohesin ChIA-Drop complexes in fragment views, we observed extensive chromatin interactions between the two SEs and their connections to the intermediate enhancers (Es) or directly to *MYC*. Many of them are multiplex interactions consisting of ≥3 chromatin fragments per complex (33% in RNAPII and 19% in cohesin ChIA-Drop data, at a higher level than genome-wide 13.5% and 13.4% for RNAPII and cohesin ChIA-Drop, respectively) (**Figure 3A**, bottom left panel). By converting the ChIA-Drop complexes into a graph representation with elements as nodes and the number of ChIA-Drop interactions as edges, we obtained a simplified representation of the mapping data: the two SEs interact highly with each other and connect to *MYC* directly or indirectly via the intermediate enhancers (**Figure 3C**, left panel). Another example is SE-*MARCKS*, which showed a strong direct connection between SE and *MARCKS* promoter (**Figure 3A-B** middle panels). By contrast, the SE on chr22 not only has a strong and direct connection to *SREBF2* but also trickles down along the path via direct and indirect contacts to the intermediate genes and enhancers to *XRCC6* (**Figure 3A-B**, right panels). Each of 188 SE-P (target promoter) connections was categorized into one of the following: 1) Multiple SEs-TP (n=33), 2) Direct SE-P (n=75), and 3) Indirect SE-P (n=80) (**Figure 3C**; see **Methods**).

To quantify the extent to which a cis-regulatory element is connected to other elements, we computed the node degrees, which is the sum of weighted edges stemming from each node (see **Methods**). There were 2,008 elements (e.g., nodes) within the 188 SE-P structures, and the node degrees of RNAPII and cohesin ChIA-Drop data were highly correlated with Pearson’s correlation coefficient of 0.8, implying that cohesin and RNAPII both establish these SE-associated interactions (**Figure 3D**; see **Methods**). Genome-wide statistics show that SEs— including other SEs along the path (OSE)—have the highest node degrees, followed by the target gene promoter (P), intermediary promoter (IP), and enhancer (E) (**Figure S3B**). These results suggest that SE acts as a hub to the other regulatory elements and target gene promoters, as well as having abundant interactions within SEs. The 188 SE-P structures in RNAPII and cohesin data have a wide range of genomic span ranging from 0.15 Mb to 5.2 Mb with a median of 3.6 Mb (**Figure S3C**, top panel) and involve a median of 6 intermediate regulatory elements (**Figure S3C**, bottom panel).

A remaining mystery is the role of multiple constituent enhancers (cE) residing within a SE. One idea is that densely clustered cE elements act together in mass to achieve stable super-enhancing transcriptional effect (Sabari et al., 2018). With the unique ability to probe multiplex chromatin interactions with single-molecule resolution, ChIA-Drop offers an opportunity to explore the details. For example, about 93% of the ChIA-Drop complexes (n=52 for RNAPII, n=60 for cohesin) connecting SE to *CD53* gene promoter involve only 1 cE, and only a few complexes (n=4 for each of RNAPII and cohesin) link 2 cEs within a SE to the target promoter (**Figure S3D**). These results indicate that the members of cEs in a SE individually interact with target promoter one-at-a-time, suggesting a probability-based mechanism instead of all of them working together in mass as previously speculated (Sabari et al., 2018).

### Long loops are sensitive to RAD21 depletion while short loops are insensitive

Our data thus far indicate that cohesin has a dual role in chromatin folding: 1) creating architectural loops by extruding chromatin at CTCF anchor sites, and 2) establishing enhancer to promoter (E-P) interactions by extruding chromatin in the direction of RNAPII transcription. However, the precise causal relationship between architectural proteins and transcription remains unknown (Merkenschlager and Nora, 2016). Given that cohesin is in the center of our questions (**Figure S4A**), we explore this problem by utilizing HCT116-RAD21-mAC cells for an acute degradation of RAD21 in HCT116 cells via auxin-inducible degron system (Natsume et al., 2016) (**Figure S4B**).

As previously reported (Rao et al., 2017), rapidly depleting RAD21 nearly abolished RAD21 signals throughout the genome, while it had no impact on CTCF binding as confirmed by ChIP-seq data (**Figure 4A, S4C, Table S1**). Surprisingly, the RNAPII binding intensity increased as auxin treatment time prolonged, implying that cohesin may modulate RNAPII’s activity to some extent. Others have suggested that cohesin may impact RNAPII movements at promoters and enhancers just as transcription can relocate cohesin (Busslinger et al., 2017; Banigan et al., 2023), a hypothesis to be tested in future studies (see **Discussions**).

**Figure 4:**
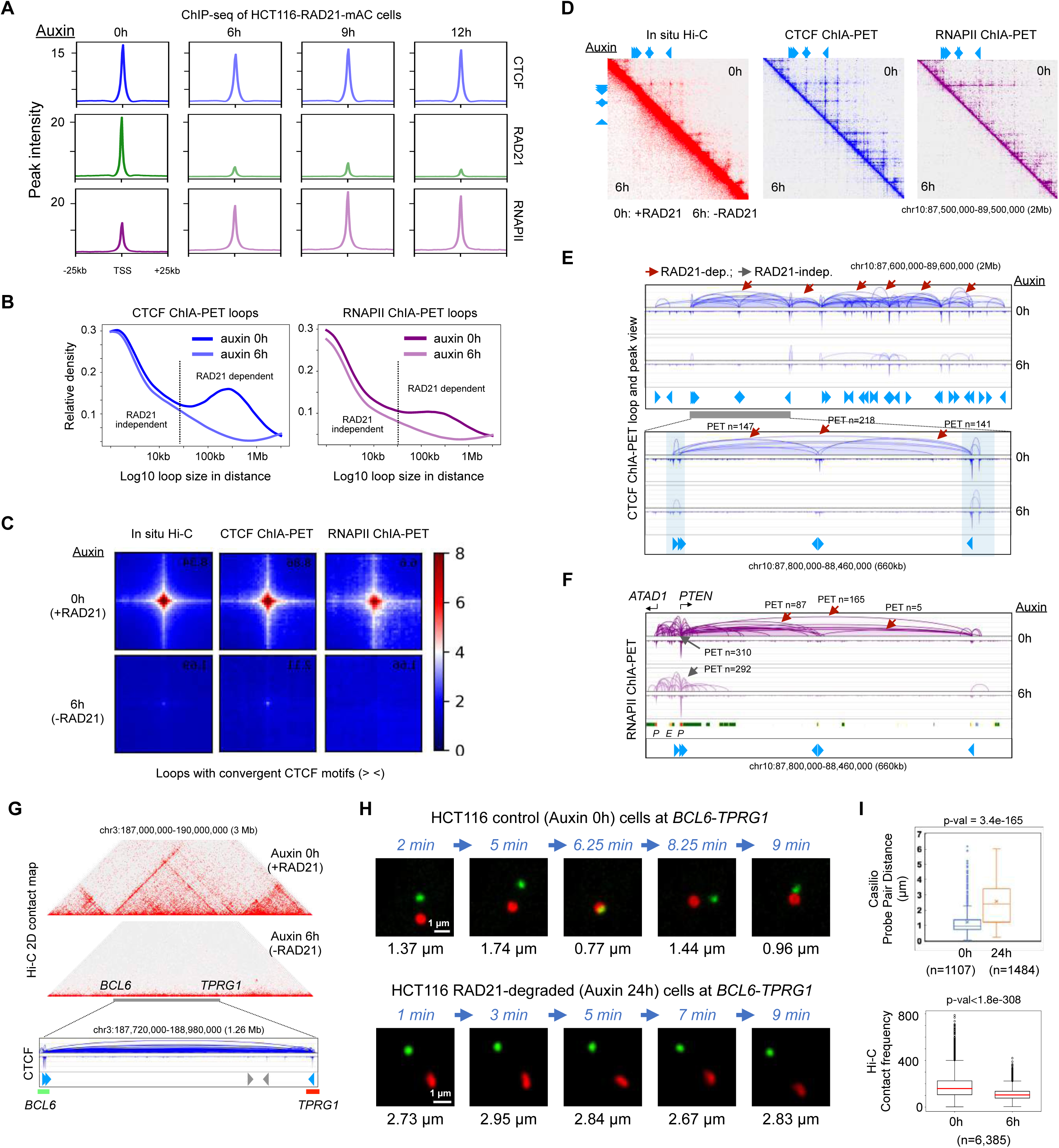
Effects of RAD21 depletion on long- and short-range chromatin interactions mediated by CTCF and RNAPII. **(A)** Aggregation plots of ChIP-seq signal at binding sites of CTCF, RAD21, and RNAPII in control (auxin 0h; h: hours) and RAD21-depleted cells treated with auxin at three time points (auxin 6, 9, and 12 hours). **(B)** A relative density function is plotted for the distance between two contact points of chromatin loops in CTCF ChIA-PET (left) and RNAPII ChIA-PET (right) data before (auxin 0h) and after (auxin 6h) RAD21 depletion. The two curves are similar for loops that are less than 30 kb in length (left side of the dotted vertical line) and deviate for loops that are greater than 30 kb, indicating that small loops (< 30 kb) are RAD21-independent and large loops (> 30 kb) are RAD21-dependent. **(C)** An aggregation of 2D contacts of chromatin loops with convergent (> <) CTCF motifs in *in situ* Hi-C, CTCF ChIA-PET, and RNAPII ChIA-PET data from HCT116 cells before (0h) and after 6 hours of auxin treatment (6h). **(D)** In a large segment of 2Mb region of chromosome 10, 2D contact maps of in situ Hi-C, CTCF ChIA-PET, and RNAPII ChIA-PET data are shown for HCT116 cell line before (0h; upper right triangle) and after (6h; lower left triangle) depleting RAD21. **(E)** Browser views of the same 2 Mb region in panel **D** and a zoomed-in region (660 kb) for CTCF ChIA-PET loops and peaks before (0h, dark blue) and after (6h, light blue) RAD21 depletion, along with CTCF binding motifs illustrated as blue arrows. RAD21-dependent (reduced) loops are indicated by red arrows and RAD21-independent (unchanged) loops are labeled with gray arrows, while PET n is the number of paired-end-tags in each loop. **(F)** The same zoomed-in region of 660 kb in panel **E** is shown for RNAPII ChIA-PET loops and peaks before (0h) and after (6h) RAD21 depletion, along with chromatin states, promoters (P) and enhancers (E) and CTCF motifs in blue arrows. **(G)** A 3 Mb chromatin domain in HCT116-RAD21-mAC used for real-time imaging analysis in live cells. Top: a 2D contact matrix of Hi-C data before (auxin 0h) and after RAD21 depletion (auxin 6h) in the 3 Mb region. Bottom: zoom-in browser view of CTCF ChIA-PET loops/peaks and binding motifs (blue) in wild-type HCT116 cells illustrating the normal chromatin looping topology. Non-repetitive regions flanking each of the two anchor sites (shown as blue arrows in convergent CTCF motifs) of this loop near *BCL6* and *TPRG1* genes are selected as the Casilio imaging targets illustrated in green and red bars, which are 1,243 kb apart. **(H)** Time-lapse images of the probe pairs with pairwise 3D distances (µm) measured at each time point in minutes (min). Scale bars, 1 µm. **(I)** Top panel: Boxplot of pairwise spot distances in untreated control (0h) (n=1,107 measurements in 21 nuclei and 27 probe pairs; mean=1.21 µm; median=0.94 µm) and 24-hour auxin-treated (24h) HCT116 cells (n=1,484 measurements in 26 nuclei and 38 probe pairs; mean=2.25 µm; median=2.00 µm). ‘x’ denotes the mean and middle line is median. Bottom panel: Boxplot of Hi-C contact frequency in genomic loci (n=6,385) with convergent CTCF motifs in untreated control (0h) and 6-hour auxin-treated (6h) cells. p-values from the two-sided Mann-Whitney U test. See also **Figure S4, Video S1, S2**.

Next, to examine the impact of cohesin perturbation on the function of CTCF and RNAPII, we generated CTCF and RNAPII ChIA-PET datasets in the presence and absence of RAD21 (**Table S1**). The CTCF loops in large distance (>50 kb) were sensitive to RAD21-depletion, implying that they are cohesin-dependent (**Figure 4B**, left panel). Globally, 18,897 CTCF loops were lost (“RAD21/cohesin-dependent”), while 4,407 loops remained (“RAD21/cohesin-independent”), with the loop span in the former category larger than the latter (median of 217 kb vs. 23 kb) (**Figure S4D**, left panel; see **Methods**). However, the roles and biophysical properties of short RAD21-independent CTCF loops remain mysterious: CTCF is capable of forming its own small loops (MacPherson and Sadowski, 2010), yet confirming such mechanism for these 4,407 loops requires additional experiments. Consistent with a previously published Hi-C data (Rao et al., 2017), convergent CTCF loops were in large distance and were cohesin-dependent as supported by CTCF ChIA-PET data genome-wide (**Figure 4C**).

Interestingly, the transcriptional loops mapped by RNAPII ChIA-PET also showed reduction upon RAD21 depletion, particularly for long-range (>50 kbps) loops (**Figure 4B**, right panel). It is known that many gene promoters are proximal to CTCF sites (Filippova et al., 1996; Cuddapah et al., 2009), as we have also confirmed (**Figure 2A**). Several studies demonstrated that RNAPII may also bind proximal to CTCF sites (Tang et al., 2015) and that some enhancer-promoter loops are CTCF-dependent (Kubo et al., 2021), but its relation to cohesin at CTCF sites has not been characterized. Our data show that RNAPII also connects convergent CTCF motif sites in a cohesin-dependent manner (**Figure 4C**), and the cohesin-dependent RNAPII loops were larger than cohesin-independent loops (median of 101 kb vs. 26 kb) (**Figure S4D**, right panel).

A representative example of a 2Mb domain on chromosome 10 exhibits a loss of most chromatin loop domains in Hi-C, CTCF ChIA-PET and RNAPII ChIA-PET data (**Figure 4D**). Specifically, CTCF loops drastically decreased in numbers and strengths even though the CTCF binding intensity remained similar before (0h) and after (6h) depleting RAD21, a feature evident in the zoomed-in 660 kb region encompassing one large loop and two small loops therein (**Figure 4E**). This large CTCF loop around an actively transcribed gene *PTEN* (same domain portrayed in **Figure 4E**) was recapitulated by RNAPII ChIA-PET profiles and mostly disappeared upon RAD21 depletion, whereas the relatively small RNAPII-associated chromatin loops connecting *PTEN* and *ATAD1* promoters resisted RAD21 ablation (**Figure 4F**).

Another example of a chromatin domain between genes *BCL6* and *TPRG1* shows a strong CTCF loop with convergent motifs in ChIA-PET data and a clear stripe pattern in our Hi-C data (see **Table S1**) of control cells (auxin 0 hours), which disappears in auxin treated (auxin 6 hours) cells (**Figure 4G**). To validate these mapping results and to measure the real-time dynamics of loop formation, we applied our live-cell imaging technique Casilio, which tags two site-specific, non-repetitive loci via a modified CRISPR-dCas9 approach (Clow et al., 2022). In particular, we tagged the regions flanking the two anchors of the *BCL6*-*TPRG1* CTCF loop (**Figure 4G**) with Clover (green) and iRFP670 (red) and tracked their movements over time with the Andor Dragonfly High Speed Confocal microscope. When the movements of paired probes were tracked over time, for example, one probe pair in a control cell became close together around 6 minutes and then separated again (**Figure 4H**, top panel**, Video S1**). We infer that a loop may have been formed in 6 minutes and dissolved thereafter. By contrast, a probe pair in a RAD21-depleted cell remained far apart throughout the measurements (**Figure 4H**, bottom panel, **Video S2**). The probe pairs in control cells had significantly closer distances (median 0.94 µm; 21 nuclei) than in the RAD21-depleted cells (median 2.41 µm; 26 nuclei) (**Figure 4I**, top panel). This trend is also in line with the Hi-C mapping data, where the loop strength—as measured by the number of chromatin complexes—is significantly higher in control than in RAD21-depleted cells (**Figure 4I** bottom panel).

Genome-wide characterization of CTCF and RNAPII loops confirm these patterns. The majority (close to 80%) of RAD21-dependent CTCF loops are convergent CTCF loops (**Figure S4E**). The RNAPII-associated loops fall into two categories (**Figure S4F**): 1) cohesin-dependent RNAPII loops (n=1,201) that are usually large (median = 101 kb), of which about 60% overlap with CTCF binding motifs and 40% with convergent CTCF loops, potentially providing a topological framework for long-range transcription regulations; 2) cohesin-independent RNAPII loops (n=4,605) that are relatively small in size (median=26 kb), mostly connecting active promoters to enhancers, and may be responsible for the transcriptional activities in the human genome as further examined below.

### Super-enhancers connect promoters of down-regulated genes through cohesin-dependent long-range interactions

To investigate the functional impacts of RAD21 depletion on gene transcription, we generated RNA-seq data in HCT116 cells before and after auxin treatments at timepoints 0 hours (no treatment), 6 hours, 9 hours, and 12 hours, and noted that auxin-treated samples deviated from 0 hour data while minimal difference is observed between 6h, 9h, and 12h (**Table S1**, **Figure S5A**). Consistent with previous findings (Rao et al., 2017), we also did not observe a drastic change in transcriptional activities but identified 356 down-regulated and 361 up-regulated genes; the majority (5,391) were unchanged upon RAD21 depletion (**Figure 5A**; see **Methods**). We further evaluated the quality of our RNA-seq data by comparing them to the published PRO-seq data (Rao et al., 2017), where both PRO-seq and RNA-seq data have statistical significance between the log2 fold-change of up-regulated and down-regulated genes (**Figure S5B**), while unchanged genes are centered around 0 (**Figure S5C**). The RNAPII occupancy signals at promoters of down-regulated genes decreased while those at promoters of up-regulated genes increased after 6 hours of auxin treatment for RAD21 depletion in RNAPII ChIA-PET data (**Figure 5B**), thereby further validating the quality of our data.

**Figure 5:**
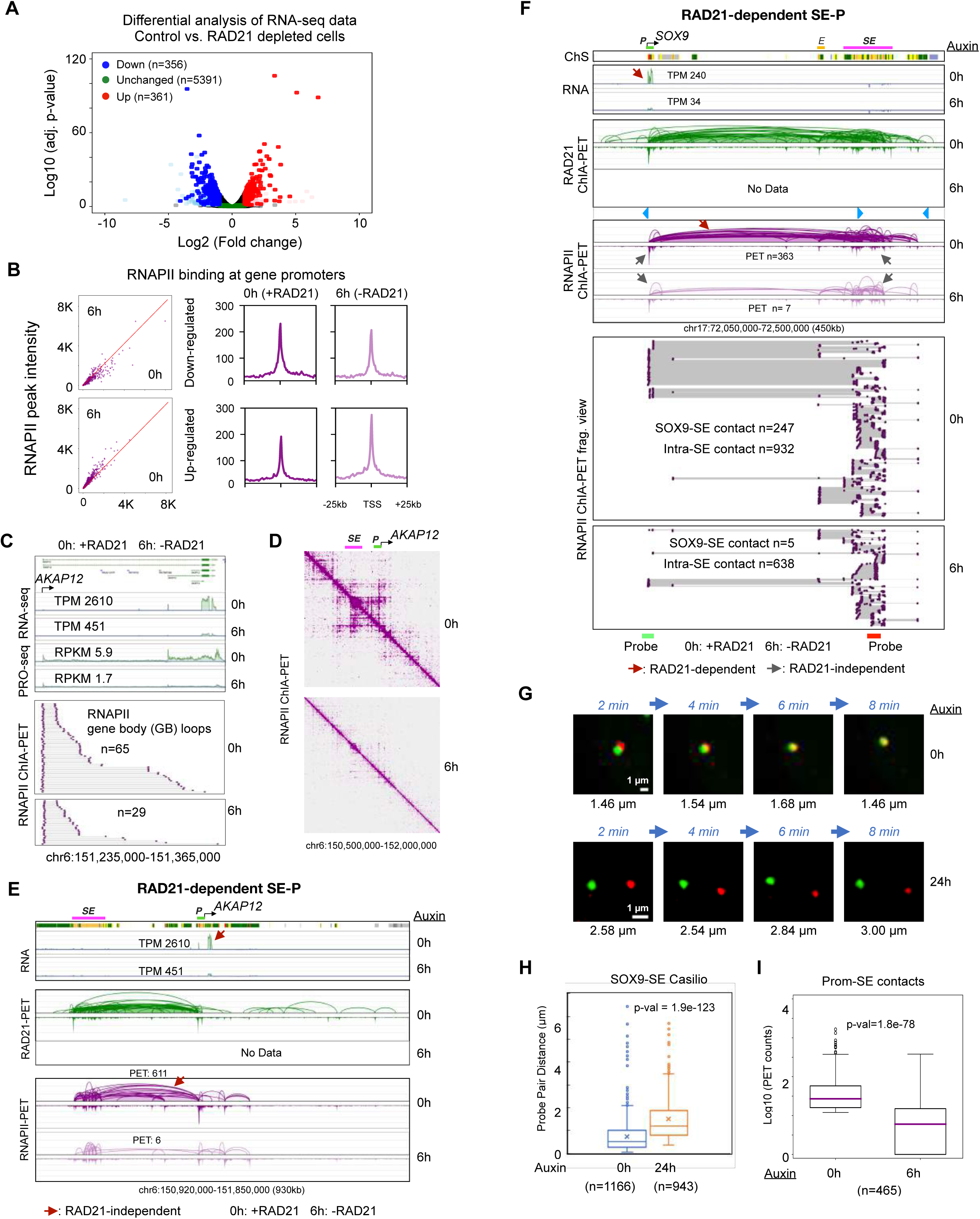
Functional roles of RAD21-dependent super-enhancer to promoter loop in transcription. **(A)** A volcano plot of differential expression analysis showing down-regulated (n=356; blue), unchanged (n=5,391; green), up-regulated (n=361; red) genes using RNA-seq data before and after auxin treatment (6 hours) in HCT116 cells. Other genes are presented in light blue, pink, and grey. See **Methods**. **(B)** RNAPII binding intensity at promoters of down-regulated and up-regulated genes with scatterplots between 0h and 6h (left panel) and aggregated peaks 25 kb upstream and downstream of TSS (right panel). **(C)** Gene body loops of RNAPII ChIA-PET data before (0h) and after (6h) depleting RAD21 are sorted from the promoter of a down-regulated *AKAP12* gene towards the transcription end site in the same forward orientation as the gene transcription; n denotes the number of chromatin complexes. TPM (transcript per kilobase million) from RNA-seq (this study) and RPKM (reads per kilobase million) from PRO-seq (Rao et al., 2017) are also recorded. **(D)** The 2D contact maps of 0h (with RAD21) and 6h (without RAD21) RNAPII ChIA-PET data are presented at a 1.5 Mb region encompassing super-enhancer (SE) and its target gene promoter (P) *AKAP12*. **(E)** A browser view of SE-*AKAP12* interactions between *AKAP12* gene promoter (P) and super-enhancer (SE). Top: tracks of ChromHMM chromatin states and RNA-seq showing that *AKAP12* is expressed with TPM (transcripts per million) of 2,610 in control cells (0h) and of 451 in RAD21-depleted cells (6h). Bottom: tracks of ChIA-PET data for RAD21 (green) and RNAPII (purple) in control (0h) and 6 hours of auxin-treated cells (6h). PET n: number of paired-end tags. **(F)** Browser views of SE-*SOX9* interactions encompassing *SOX9* promoter (*P*) and the associated enhancer (*E*) and super-enhancer (*SE*). Top: tracks of ChromHMM chromatin states (ChS) demarcating *SOX9* promoter (*P*), enhancer (*E*), and super-enhancer (*SE*), and RNA-seq data in control cells (0h) and RAD21-depleted cells (6h). Middle: tracks of ChIA- PET data for RAD21 (green) and RNAPII (purple) in control (0h) and 6 hours auxin treated cells (6h) capturing chromatin loops connecting *SOX9* to distal enhancer (*E*) and super-enhancer (*SE*). The RAD21-dependent (reduced) loops and RAD21-independent (unchanged) loops are indicated with red and grey arrows, respectively. Bottom: RNAPII ChIA-PET fragment view for connections of *SOX9-E, SOX9-SE, E-SE,* and intra*-SE*. The approximate genomic positions of probes (green and red bars) for Casilio live cell imaging are depicted. TPM: transcripts per kilobase million; PET n: number of paired-end tags. **(G)** Representative time-lapse images of the probe pairs indicated by arrows in **Figure S5G** with pairwise 3D distances (µm) recorded at each time point in minutes (min) in both control (0h) and Auxin-treated (24h) cells. Scale bars, 1 µm. **(H)** Boxplot of Casilio distances between the paired green and red probes for *SOX9*-SE (super-enhancer) loop in control (0h) cells (1,166 measurements of 31 pairs in 17 nuclei; mean=0.74 µm, median=0.53 µm) and auxin-treated (24h) RAD21-depleted cells (943 measurements of 23 pairs in 15 nuclei; mean=1.51 µm; median=1.20 µm) cells. ‘x’ denotes the mean, and middle line is median. p-value from the two-sided Mann-Whitney U test. **(I)** Boxplot of PET counts between target gene promoters and SE in RNAPII ChIA-PET data in log10 scale. p-value from the two-sided Mann-Whitney U test. See also **Figure S5, Video S3, S4**.

The gene ontology (GO) results (**Figure S5D**) indicate that the down-regulated genes in RAD21-depleted cells were enriched in functions for transcriptional regulation in response to stress, while the unchanged genes had general housekeeping functions such as RNA processing and translation. Thus, down-regulated genes may be related to regulatory and cell-type specific functions, while the unchanged genes are constitutive and for housekeeping roles. Intriguingly, our GO results also suggest that the up-regulated genes upon RAD21 depletion were almost exclusively involved in DNA replication (**Figure S5D**), which hints another dimension of cohesin’s functions in organizing the genome as we later recapitulate. We next focused on these down- regulated and unchanged genes for potential topological mechanisms.

One of the down-regulated genes is *AKAP12*, which was identified as statistically significant by both RNA-seq and PRO-seq data with fold-change 5.8 and 3.5, respectively (**Figure 5C**). We have previously proposed a loop reeling model for transcription in *Drosophila*, where the RNAPII reels in chromatin from the transcription start site (TSS) to the end site (TES) (Zheng et al., 2019). Given that each contact in ChIA-PET data represents a single-molecule interaction derived from a single cell, we converted paired-end tags to chromatin complexes and applied a similar method to sort complexes from TSS to TES; these are referred to as ‘gene body’ loops. As a result, a down-regulated *AKAP12* gene had 2.2-fold reduction in gene body loops (**Figure 5C**), while an unchanged *PTEN* gene had small 1.2-fold change (**Figure S5E**). A genome-wide statistics support that down-regulated genes are accompanied by a reduction in gene-body loops and unchanged genes retain similar numbers without a statistical difference between 0h and 6h data (**Figure S5F**).

We next investigated a potential molecular mechanism by which a down-regulated gene represses its expression level. We have established that SE-P interactions are mediated by both cohesin and RNAPII (**Figure 3, S3A**) and that the genomic span is large, with a median of ∼360 kb (**Figure S3C**). Revisiting *AKAP12*, we observed that overall, its promoter was highly connected to a distal super-enhancer and other regions, but such strong connections disappear without RAD21 (**Figure 5D**). In a further zoomed-in 930kb region, the promoter of *AKAP12* is tightly connected to a distal (∼300 kb) upstream SE as shown in both RAD21 and RNAPII ChIA-PET data (**Figure 5E**). However, the RNAPII-associated SE-*AKAP12* interactions were reduced by 100-fold (611 vs. 6 PET counts) after RAD21 degradation, potentially contributing to the 5.8-fold reduction in *AKAP12* gene expression. The intra-SE and the small loops around the gene were minimally affected, which may contribute to the maintenance of a basal level expression of *AKAP12*.

In another example, the promoter of an actively expressed gene *SOX9* was highly connected to a downstream SE via long-range chromatin loops (∼320 kb) as evidenced in both cohesin and RNAPII ChIA-PET data and not overlapping with convergent CTCF motifs (**Figure 5F**). However, after RAD21 depletion, the long-range RNAPII interactions were dramatically reduced from PET counts 363 down to merely 7 PET counts, accompanied by more than 7-fold (from 240 TPM to 34 TPM) reduction in expression level. The chromatin fragment views of the RNAPII ChIA-PET data in single-molecule resolution revealed tight connections of *SOX9*-SE before auxin treatment, and a 50-fold reduction after auxin treatment (PET counts before treatment n=247 vs. after treatment n=5). By contrast, the small loops (up to 60 kb) within the SE (intra-SE) showed only a moderate 1.5-fold change after auxin treatment (intra-SE PET counts before treatment n=932 vs. after treatment n=638 PET counts) (**Figure 5F**), further confirming that large loops are more dependent on cohesin than small loops are.

To validate the observed long-range *SOX9*-SE contacts through an orthogonal imaging method, we designed Casilio probes proximal to the loci of *SOX9* (Clover; green) and SE (iRFP670; red) (**Figure 5F**). An exemplary nuclear image showed that the two paired probes are extremely close to each other in the control (auxin 0h) and far from each other in RAD21-depleted (auxin 24h) cells (**Figure S5G**). In live cells with time-lapse imaging over 8 minutes, the probe pair in control cells stayed close together (**Figure 5G** top panel**, Video S3)**, but far apart in RAD21- depleted cells (**Figure 5G** bottom panel**, Video S4**). The overall distances between the paired probes in control cells were much closer (n=1166, median=0.53 µm) than the probe pairs in the RAD21-depleted cells (n=943, median=1.2 µm) (**Figure 5H**). This pattern is also in line with our RNAPII ChIA-PET mapping data, where SE-P connection counts (PETs) are dramatically reduced upon RAD21 depletion genome-wide (**Figure 5I**). Together, our mapping and imaging results provided quantitative evidence for us to deduce that cohesin-dependent long-range chromatin loops involving SE-P and E-P are a likely topological mechanism that provides a structural framework for regulating transcription of genes that are sensitive to RAD21 depletion (cohesin-dependent).

Finally, we re-visited the *MYC* location examined in **Figure 3** to test whether *MYC* is connected to the same super-enhancer (SE) in HCT116 cells as in GM12878 cells, and more importantly, whether the connections are RAD21-dependent. The 2D contact maps of Hi-C, CTCF, RNAPII, and RAD21 ChIA-PET data show that *MYC* is highly connected to upstream SE and downstream enhancer (E), which all vanish without RAD21 (6h) (**Figure S5H**). A closer examination of chromatin loops confirms that *MYC* promoter is connected to SE and E by both RAD21 and RNAPII, and that *MYC*-E loop and *MYC*-SE loop are attenuated in line with a 1.3-fold reduction in gene expression (**Figure S5I**). Since *MYC* is an oncogene with many functions, we sought to answer whether its transcription is activated in a similar pattern across cell lines. As we had observed, GM12878 cells have exclusive connection to two upstream super-enhancers, and 4 other cell lines have highly variable patterns: MCF7 and K562 show connections only to downstream enhancers (distinct to cell line, denoted by chromHMM states), HepG2 with both upstream and downstream contacts, and H1 has minimal loops to a nearby enhancer (**Figure S5J**). This intriguing observation led us to further investigate genome-wide lineage-specific transcriptional loops with respect to cohesin.

### Genes associated with short loops are cohesin-independent

The patterns observed in SE-P were also found in many long-range enhancer-promoter (E-P) interactions that are RAD21-dependent and are associated with genes that are down-regulated due to RAD21 depletion. Of 130 down-regulated genes with measurable high quality chromatin loops stemming from their promoters, more than half (n=71) had chromatin loops that were RAD21-dependent, and close to 75% (n=50) were characterized as long-range enhancer-promoter (E-P) loops. For example, *UPP1* is expressed in HCT116 cells and is down-regulated upon RAD21 depletion with a 4-fold reduction in transcripts from TPM counts 192 to 53 (**Figure 6A**). The promoter of *UPP1* is connected to a promoter of another nearby gene (*HUS1*) and 3 enhancers that are approximately 100 kb and up to 1 Mb upstream. However, these RNAPII chromatin loops were diminished in RAD21-depleted cells, resulting in a complete loss of chromatin loops stemming from the promoter of *UPP1*, as evident in both 2D contact maps and loops. Another gene *CDKN2B* is down-regulated with its expression level reduced by a 14-fold (TPM 96 to 7), and its promoter had significant reduction in RNAPII signals and chromatin loops that connect to distal enhancers (**Figure S6A**).

**Figure 6:**
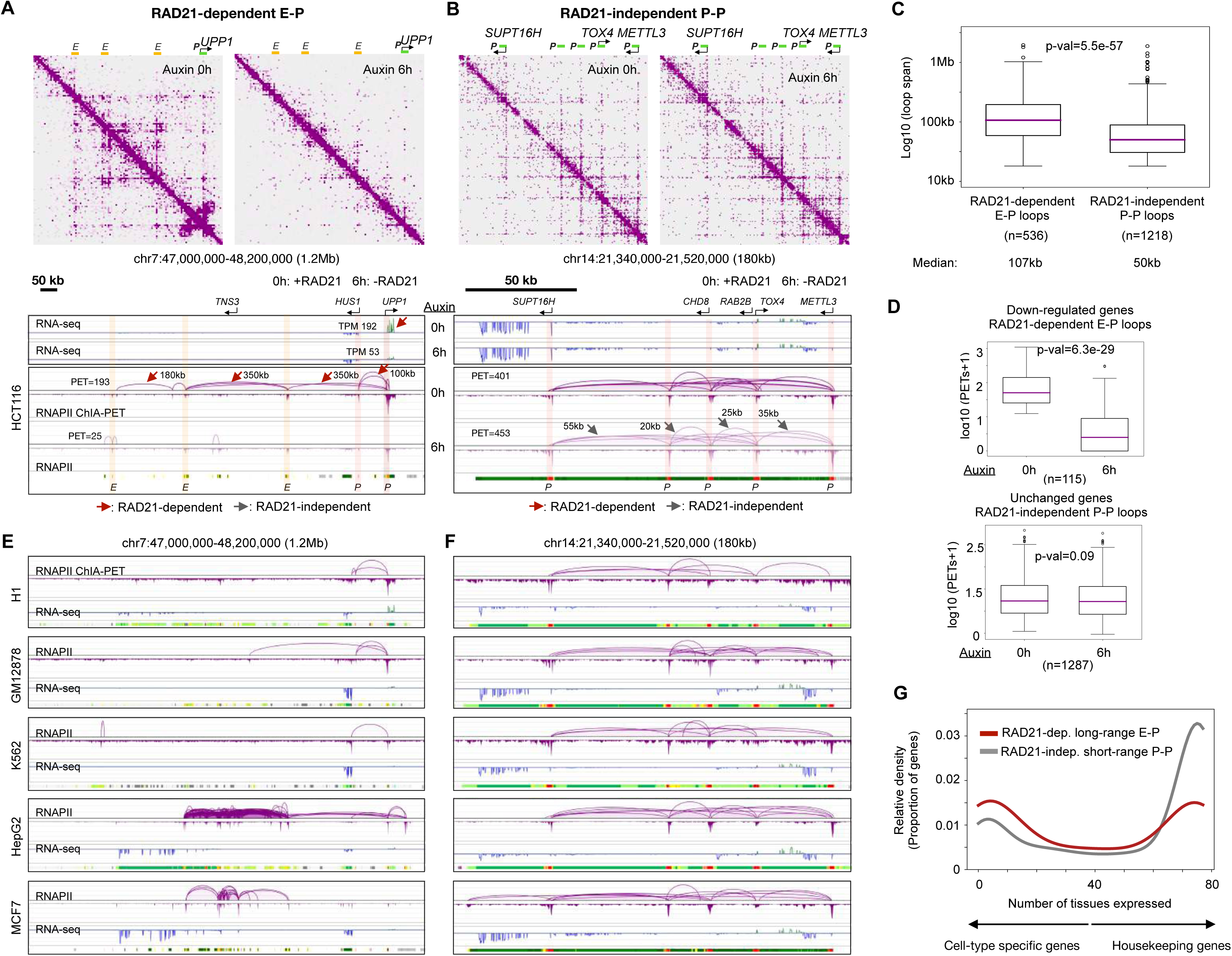
Distinct roles of RAD21-dependent E-P and RAD21-independent P-P RNAPII loops in gene regulation. **(A)** An example of RNAPII-associated chromatin loops attenuated by RAD21 depletion. In a large chromatin domain (1.2 Mb) harboring *UPP1* gene and associated regulatory elements enhancer (*E*) and promoter (*P*) demarcated by ChromHMM, 2D contact maps of RNAPII ChIA-PET before (0h) and after (6h) depleting RAD21 are shown. Below, tracks of RNAPII ChIA-PET loops/peaks views in HCT116 cell line connect *UPP1* gene promoter (*P*) to many distal enhancers (*E*), with RAD21-dependent attenuated loops marked by red arrows. TPM: transcripts per kilobase million; PET n: number of paired-end tags. **(B)** An example of RAD21-independent RNAPII loops connecting active gene promoters. In a 180 kb region harboring *METTL3* and other genes, 2D contact maps of RNAPII ChIA-PET before (0h) and after (6h) depleting RAD21 are shown. Below, the tracks of RNA-seq and RNAPII ChIA-PET data show connections of active gene promoters that are between 20 kb and 55 kb apart, and these short-range P-P loops are not affected by the RAD21 depletion (grey arrow). Active gene promoters as annotated with ChromHMM states are highlighted in red. **(C)** Boxplot of loop span of RAD21-dependent enhancer-promoter (E-P) and RAD21-independent promoter-promoter (P-P) loops. Number of loops (n) used for plotting and the median of loop span are provided. **(D)** Boxplots of PET numbers of RAD21-dependent E-P loops involving down-regulated genes (top panel) and of RAD21- independent P-P loops connecting unchanged genes (bottom panel). **(E)** In the same region as panel **A**, the RNAPII ChIA-PET loops and peaks, RNA-seq coverage, and ChromHMM states are also shown for 5 other cell lines: H1, GM12878, K562, HepG2, MCF7. **(F)** Same region as panel **B** with the same annotation as panel **E**. **(G)** Density plot of the number of tissues (out of 76 tissues), in which each gene is expressed, for those connected in RAD21-dependent E-P loops (red) and those in RAD21- independent P-P loops (grey) (see **Methods**). See also **Figure S6**.

The long-range (median of 101 kb) chromatin loops that are sensitive to RAD21 depletion are important for down-regulating a handful of genes (**Figure S4D**). However, most RNAPII-associated chromatin loops are in short-range (median of 26 kb) and are not affected by RAD21 depletion, i.e., cohesin-independent (**Figure S4D, S4F**). We investigated the composition of the short-range chromatin loops that are associated with genes that do not significantly change in expression levels after RAD21 depletion. In 1,802 unchanged genes with measurable chromatin looping in RNAPII ChIA-PET data, 93% of the chromatin loops were cohesin-independent, and the majority (75%) of these loops were involved in connecting promoters of active genes (P-P interactions). For instance, in a 180 kb window on chromosome 14 harboring multiple active genes, the P-P chromatin loops connect the promoters of 5 genes (*SUPT16H*, *CHD6*, *RAB2B*, *TOX4*, and *METTL3*) and are in short-range from 20 kb to 55 kb (**Figure 6B**). In line with the notion that long-range (median of 101 kb) loops disappear, and short-range (median of 26 kb) loops remain after RAD21 depletion (**Figure S4D**), these short daisy-chain-like RNAPII loops and gene expression seem to persist without RAD21 in both 2D contact maps and loop views. A similar example is included (**Figure S6B**).

Genome-wide statistics support a general principle that RAD21-dependent enhancer-promoter loops are long with a median of 107 kb, while RAD21-independent promoter-promoter loops are short with a median of 50 kb (**Figure 6C**). Furthermore, down-regulated genes are often connected to enhancer-promoter loops that are RAD21-dependent (**Figure 6D**, top panel), whereas genes showing little or no changes in transcription upon RAD21 depletion are often connected by cohesin-independent promoter-promoter (P-P) loops (**Figure 6D**, bottom panel).

### RAD21-dependent loops and genes are cell-type specific while RAD21-independent loops and genes are constitutive

To explore whether our results generalize to other cell types beyond HCT116, we analyzed RNAPII ChIA-PET and RNA-seq data in 5 additional human cell lines from the ENCODE consortium (Luo et al., 2020), including embryonic (H1), lymphoblastoid (GM12878), erythromyeloblastoid (K562), hepatocyte (HepG2), and epithelial of breast cancer patient (MCF7), just as we had previewed in **Figure S5J**. We found that at chromatin domains harboring differentially expressed genes (e.g. *UPP1*, *CDKN2B*), chromatin loop and gene expression profiles were highly variable among the 6 cell lines (**Figure 6E** and **Figure S6A**, bottom panels). By contrast, for genes associated primarily with cohesin-independent loops, the patterns of gene expression, chromatin loop, and the chromatin states (ChromHMM) were largely invariable across different cell lines (**Figure 6F, Figure S6B**, bottom panels). A quantitative way to measure the heterogeneity is by the normalized Shannon entropy, which indicates that RAD21-dependent E-P genes and loops tend to be more heterogeneous than RAD21-independent P-P genes and loops (**Figure S6C**; see **Methods**). Furthermore, genome-wide statistics support the idea that genes relying on cohesin-dependent loops tend to be cell-type-specific (**Figure S6D**), while genes that are organized by short-range P-P chromatin loops are constitutive active across tissues (**Figure S6E**).

We analyzed an RNA-seq data collection from 76 human tissues (Papatheodorou et al., 2019), and quantified the number of tissues in which a given gene is expressed. The higher the number, the more likely a gene carries out “housekeeping” functions; a low number would imply that a gene is highly tissue- or cell-type-specific. As a result, genes involved in RAD21-dependent long-range E-P interactions were predominantly cell-type-specific, while those connected by RAD21-independent short-range P-P loops were in general ubiquitously expressed, and therefore, may potentially function as “housekeeping” genes (**Figure 6G**; see **Methods**). Recall that GO analysis (**Figure S5D**) also suggested that the down-regulated genes by RAD21 depletion were enriched in functions for transcriptional regulation in response to stress, while the unchanged genes had general housekeeping functions such as RNA processing and translation. Collectively, our analyses of gene expression and function support a cohesin-associated topological mechanism in which cohesin-dependent long-range E/SE-P loops regulate cell-type-specific genes, while cohesin-independent short-range P-P interactions maintain constitutively expressed housekeeping genes.

### DNA replication signals of up-regulated genes drastically amplify upon RAD21 depletion

Intriguingly, the up-regulated genes (n=361; **Figure 5A**) were enriched in DNA replication term in the GO analysis (**Figure S5D**). Many of the genes in this class are indeed well-known for possessing DNA replications functions, including the origin recognition complex subunit 1 (*ORC1*), DNA replication complex GINS protein PSF1/2 (*GINS1/2*), and G1/S-specific cycline-E1/2 (*CCNE1/2*) (**Figure S7A**). This result motivated us to re-analyze the published Repli-seq data with early (P02) to late (P17) DNA replication stages in HCT116 cells before (0h) and after (6h) depleting RAD21 (Emerson et al., 2022).

The genome-wide statistics of taking the median replication signal in the gene body (see **Methods**) revealed distinct properties of up-regulated genes (**Figure 7A**). While all three categories of up-regulated, down-regulated, and unchanged genes generally had higher signal in the early replication stage than late in HCT116 cells before RAD21 depletion (0 hour), the median signal of P04 was highest in up-regulated genes (0.7) compared to down-regulated genes (0.6) and unchanged genes (0.64). More prominent features are in the RAD21-depleted cells (6 hour): only up-regulated genes have close to 2-fold increase in P04 and P05, and all values from P04 to P17 are higher in 6 hour than 0 hour (**Figure 7A**). Unchanged genes have similar patterns as down-regulated genes, while random control regions have constant median replication signals around 0.3 throughout all stages both before (0h) and after (6h) RAD21 depletion.

**Figure 7:**
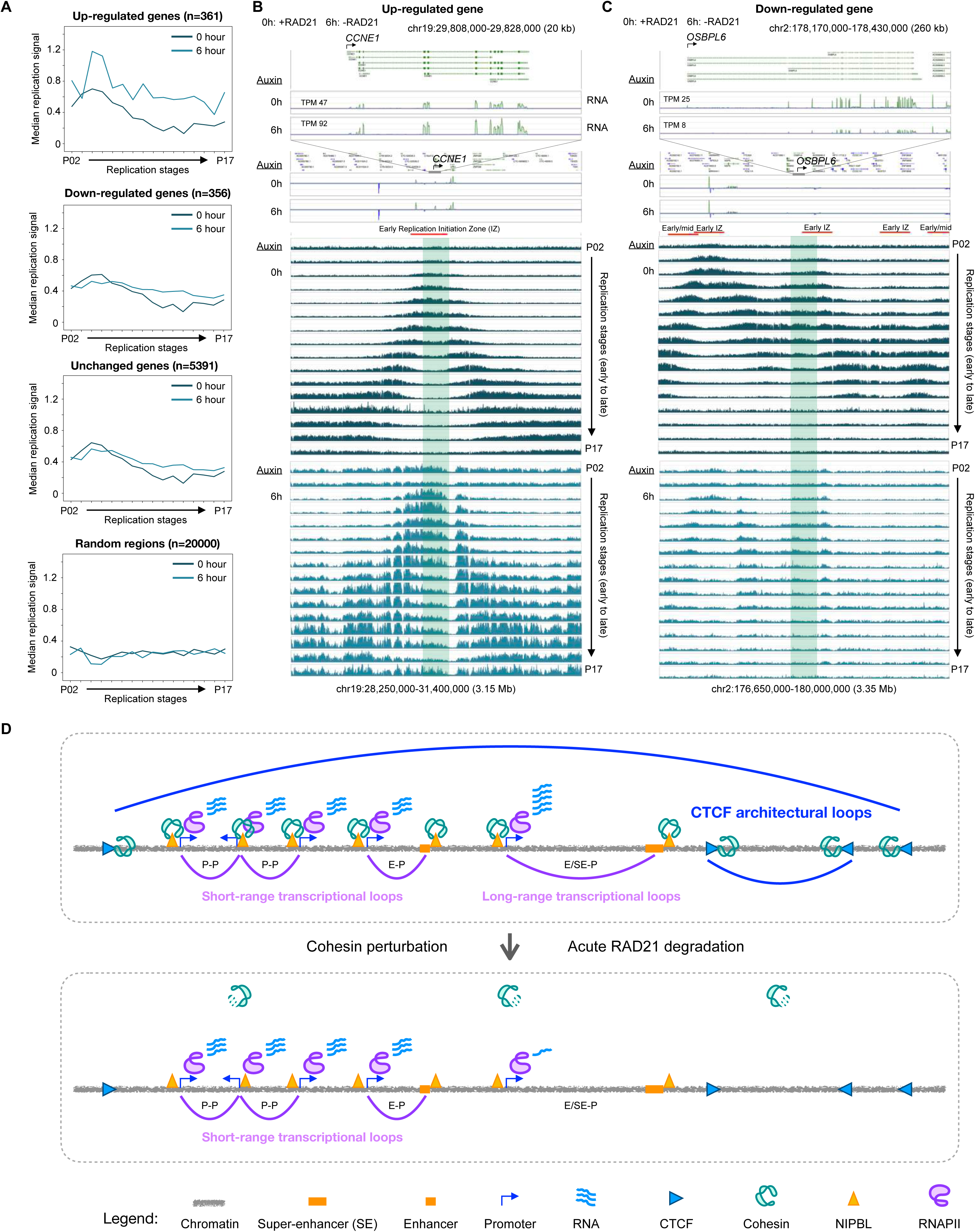
DNA replication signal patterns in differential genes, and proposed model. **(A)** The median replication signal of gene body is plotted for each of the 16-stage Repli-seq data before (0 hour) and after (6 hour) depleting RAD21 for up-regulated, down-regulated, and unchanged genes, along with 20000 random regions (see **Methods**). **(B)** An up-regulated gene *CCNE1* with TPM (transcript per kilobase million) computed from RNA-seq data before (0h: 0 hour) and after (6h: 6 hours) depleting RAD21. A larger 3.15 Mb region encompassing *CCNE1* is shown with 16-stage Repli- seq data (Emerson et al., 2022) from early P02 to late P17 replication stages with auxin treatment denoting cell with (0h) or without (6h) RAD21. Red bar indicates the early initiation zone defined by Emerson et al. using HCT116 0h Repli-seq data. **(C)** Similar to panel **B**, for a down-regulated gene *OSBPL6*. **(D)** Chronologically, cohesin first loads to chromatin at NIPBL binding sites that are usually co-localized with RNAPII and active transcriptional elements (promoters, enhancers), then it goes along with RNAPII in the direction of transcription and establish short-range transcriptional interactions for local constitutive genes (P-P and E-P) and in long-range loops for connecting distal (super-)enhancers to target gene promoters (SE-P); after arriving at CTCF binding sites, cohesin is interlocked with CTCF, anchors itself there, and actively reels in DNA string in accordance with the CTCF motif orientation, thereby constituting large architectural loops. The architectural loops (blue), the long-range transcriptional loops (purple for RNAPII), and the associated genes are sensitive to RAD21 depletion (cohesin-dependent), whereas the short-range transcription loops (purple for RNAPII) and associated genes are cohesin-independent. See also **Figure S7**.

For example, one of the up-regulated genes is *CCNE1*, which had a 2-fold increase in expression (TPM of 47 vs. 92) (**Figure 7B**, top panel). In a 3.15 Mb region centered around *CCNE1*, the replication signal exhibits a typical pattern of early initiation zone (EIZ) with the center enriched signals extending symmetrically outward as the stage of DNA replication timing progresses in control cells (**Figure 7B**, middle panel). Remarkably, upon RAD21 depletion (6h), the replication signals in all stages were drastically increased and the pattern seems to be disrupted over a megabase window (**Figure 7B**, bottom panel). Another up-regulated gene *RRM2* with 1.9-fold increase (TPM 193 vs. 367) also has a clear enrichment of early replication signal in 0h and such pattern is abolished in 6h with magnitudes of higher signals surrounding the gene (**Figure S7B**).

A stark contrast is shown for a down-regulated gene *OSBPL6* (**Figure 7C**, top panel). The replication signal in 0h also has enriched signal in early stages, but unlike up-regulated genes, it does not have the ‘upside down V’ shape in the 3.35 Mb region surrounding the gene (**Figure 7B**, middle panel). More importantly, upon depleting RAD21 (6h), the replication signals at the gene body remain similar and the broader 3.35 Mb region is lacking replication signal. An additional example is shown for the *FRAS1* gene (**Figure S7C**). These two examples demonstrate that the amplification of replication signal in up-regulated genes for 6h (**Figure 7B, S7B**) is not observed in down-regulated genes.

Most studies could not fully explain the molecular mechanism by which a subset of genes is up-regulated upon depleting RAD21 (Rao et al., 2017; Hsieh et al., 2022). Our analyses conclude a comprehensive characterization of up-regulated, down-regulated, and unchanged genes with respect to multiple dimensions of genome organization from chromatin looping to DNA replication.

## DISCUSSION

In this study, we investigated the intricate interplays between cohesin and RNAPII. By applying single-molecule ChIA-Drop technique with chromatin immunoprecipitation, we obtained highly specific multiplex chromatin interaction maps in the human genome. Our results showed that cohesin and RNAPII are highly correlated in establishing transcriptional loops, suggesting multifaceted roles of cohesin: 1) forming architectural loops with CTCF at convergent motif sites; 2) mediating and maintaining long-range (super-)enhancer to promoter interactions with RNAPII; 3) possibly mediating but not maintaining short promoter-promoter loops; 4) organizing gene body loops locally from TSS to TES. More specifically, with ChIA-Drop data we are able to capture a large number of multiplex transcriptional loops that connect 3 or more distal regulatory elements, providing evidence with single molecule mapping resolution to support a complex topological framework for co-transcription regulation.

Considering that both cohesin and RNAPII have their own motors to move along DNA templates, a particular unresolved question is how these two different types of molecular motors reconcile with each other to coordinate their action. We provided evidence that cohesin follows the direction of transcription at active gene promoters in a loop reeling process similar to that we had observed in *Drosophila* (Zheng et al., 2019). In line with this idea, previous studies showed that knocking out both CTCF and WAPL in mouse cells results in cohesin accumulation at sites of convergent transcription from ChIP-seq experiments, thereby creating ‘islands’ (Busslinger et al., 2017), and that RNA Polymerase can be a barrier to cohesin loop extrusion (Banigan et al., 2023). Based on the single-molecule imaging experiments supporting the two-sided extrusion of cohesin *in vitro* (Kim et al., 2019), we hypothesize that cohesin itself does not have a directionality and can be inherently bidirectional, but with the influence of external factors such as CTCF and RNAPII it may be biased towards one direction. One way to directly validate this idea is to deplete RNAPII and perform RAD21 ChIA-Drop or ChIA-PET experiments. If RNAPII is indeed the driving force determining cohesin’s direction of movements, then cohesin should lose directionality at gene promoters and not follow transcription, similar to its behavior at loading sites without TSS (**Figure S2D**). To measure the effect of CTCF on cohesin, one may genetically introduce CTCF binding sites at cohesin loading sites and quantify the changes in directionality. Cohesin and RNAPII are also highly correlated in connecting distal enhancers to promoters, implying that cohesin may cooperate with RNAPII in establishing transcriptional loops; in parallel, a recent single-cell Micro-C experiments show that cohesin forms transcription elongation loops (Wu et al., 2023).

The latest major conundrum in chromatin biology is that acute depletion of chromatin architectural proteins has only a marginal effect on gene expression (Rao et al., 2017; Hsieh et al., 2022). Conversely, depletion of RNAPII eliminates transcription but results in only subtle or no changes in chromatin folding (El Khattabi et al, 2019; Jiang et al., 2020). These observations raise valid concerns: namely, if the chromatin folding topology is truly relevant to the function of gene transcription. It is possible that Hi-C or Micro-C (Nora et al., 2017; Xu et al., 2021) experiments lacked resolution or were not specific enough to enrich protein-mediated chromatin contacts necessary for tackling this question, even though Micro-C could identify some enhancer-promoter interactions (Hsieh et al., 2022; Zhang et al., 2023) at the cost of deep sequencing lack of protein specificity. Here, we demonstrated that cohesin is critical for mediating long-range transcriptional interactions between distal enhancers and promoters of cell-type specific genes, but it is not required for maintaining short-range transcriptional loops connecting promoters of constitutively expressed genes with housekeeping functions. Given that most active genes in a cell are constitutive and most are organized via short-range (around 50 kb) transcriptional loops, our results resolve the puzzle that transcriptional landscape does not change dramatically despite the loss of large loop domains. However, the molecular mechanism by which cohesin-independent short-range transcriptional loops are created is yet to be determined. One hypothesis is that due to the polymer property of chromosomes, once established, short-ranged transcriptional loops are self-sustainable in the absence of cohesin. An alternative scenario is that other specific protein factors facilitate stable contacts of housekeeping genes for constitutive transcription, while the long-range loops for cell-type specific and/or regulatory genes may require a rather flexible mechanism to stay responsive. Cohesin has been extensively studied in terms of biophysical properties (Banigan et al., 2017), protein structures (Li et al., 2020), and energetics (Vian et al., 2018). Yet our results show that cohesin is responsible for maintaining only long-range chromatin interactions. We are optimistic that a similar level of enthusiasm will emerge for studying properties of short cohesin-independent loops.

It is known that transcription and DNA replication are tightly correlated and that cohesin has also been suggested to have roles in DNA replication as well, with an idea that cohesin is recruited to DNA replication origins in *Drosophila* (Pherson et al., 2019). Alongside chromatin loops formed by various protein factors, the DNA replication and cell cycle are also implicated in organizing the mammalian genome in 3-dimensional space (Hand, 1978; Nagano et al., 2017; Zhang et al., 2019; Emerson et al., 2022). By re-analyzing publicly available Repli-seq data with respect to our differential genes before and after depleting RAD21, we reported an interesting distinct pattern for up-regulated genes: the replication signal is highly amplified and the ‘upside down V’ pattern is disrupted without RAD21 (**Figure 7A, S7A**). One reason for such an exclusive outcome for up-regulated genes is that they were enriched in genes with functional roles in DNA replication and hence lost key molecules to maintain proper DNA replication in the absence of RAD21, while down-regulated and unchanged genes were not related to replication in GO analysis. Moreover, given that up-regulated genes were two-fold shorter than down-regulated genes, we speculate that these short genes are more vulnerable and prone to be exposed for ectopic gene activation. However, rigorous follow-up studies are required to identify the precise causal impact of RAD21 depletion on DNA replication of up-regulated genes. With a new technique Repli-HiC (Liu et al., 2024) to capture chromatin interactions involving nascent DNA, we envision the field to further dissect the relationship between DNA replication and genome topology.

Together, with the novel insights provided in this study, an integrated perspective is proposed for the interconnected dual roles of cohesin in transcription regulation and chromatin loop formation (**Figure 7D**). It is speculated that at its loading (and transcriptionally active) sites, cohesin translocates along chromatin together with RNAPII to establish transcription loops connecting promoters and enhancers in both short-range for constitutive genes and long-range for regulatory and cell-specific genes; at its anchoring site, cohesin interacts with CTCF to robustly reel in chromatin to create loops. However, cohesin is required for maintaining only the long-range interactions and not short loops. There are still many unanswered questions. For example, how exactly cohesin and RNAPII act together at establishing transcriptional loops; why cohesin is more critical for long-range transcription loops than the short-range ones; what conformational changes occur when cohesin and CTCF are connected at the anchoring sites, which lead to chromatin reeling. We anticipate that additional research efforts will be invested in these and related directions in the near future.

## ACKNOWLEDGEMENTS

This study was supported by the Jackson Laboratory Director’s Innovation Fund (DIF19000- 18-02 to Y.R.), 4DN (U54 DK107967 to Y.R.) and ENCODE (UM1 HG009409 to Y.R.) consortia, Human Frontier Science Program (RGP0039/2017 to Y.R.), the National Human Genome Research Institute (R01-HG009900 to A.W.C., R01-HG011253 to C.L.W., R01-GM127531 to C.L.W., K99-HG011542 to M.K.), and National Science Foundation (CCF-1955712 to O.M., CIF 1956384 to O.M.). The authors acknowledge Zhihui Li, Xiaoan Ruan, Meizhen Zheng, and Simon Tian for the preliminary ChIA-Drop data generation and analyses and thank the Casellas group members for critical feedback on the manuscript.

## AUTHOR CONTRIBUTIONS

Y.R. conceptualized the project. Y.R., M.K., P.W. designed experiments. P.W., H.C., X.L., C.N., C-L. W. generated mapping data. M.K., I.C., X.W., J.P., B.L. performed computational analyses. F.Y., O.M., J.H.C., C.L.W., R.C., Y.R. assisted data analysis and interpretation. A.W.C., Y.R., P.A.C., and M.K. designed the imaging experiments. P.A.C. and A.W.C. performed the imaging experiments and analyzed the data. Y.R., B.L., and M.K. assisted imaging data analysis and interpretation. Y.R. and M.K. wrote the manuscript with inputs from P.W., R.C., P.A.C., and A.W.C.

## RESOURCE AVAILABILITY

Further information and requests for resources and reagents should be directed to and will be fulfilled by the Lead Contact, Yijun Ruan.

## Materials availability

This study did not generate new unique reagents.

## Data and code availability

The accession number for the deep-sequencing data reported in this paper is GEO: GSE158897.

## CONTACT FOR REAGENT AND RESOURCE SHARING

Further information and requests for resources and reagents should be directed to, and will be fulfilled by the Lead Contact, Yijun Ruan.

## KEY RESOURCES TABLE

| REAGENT or RESOURCE | SOURCE | IDENTIFIER |
| --- | --- | --- |
| **Antibodies** |  |  |
| RAD21 | Abcam | Cat# ab922 |
| WAPL | Abcam | Cat# ab70741 |
| RNA Polymerase II | BioLegend | Cat# 664912 |
| CTCF | Abclonal | Cat# ab70303 |
| NIPBL | Bethyl Laboratories | Cat# A301-779A |
| SMC1 | Bethyl Laboratories | Cat# A300-055A |

## CRITICAL COMMERCIAL ASSAYS

|  |  |  |
| --- | --- | --- |
| Nextera DNA sample preparation kit | Illumina, Inc | Cat# FC-121-1030 |
| Nextera Index kit | Illumina, Inc | Cat# FC-121-1011 |
| LightCycler 480 SYBR Green I Master | Roche | Cat# 04887352001 |
| DNA Clean & Concentrator-5 | Zymo research | Cat# 4013 |
| ChIP DNA clean & Concentrator | Zymo research | Cat# D5205 |
| QIAquick PCR purification kit | Qiagen | Cat# 28104 |
| Amicon Ultra-0.5 Centrifugal Filter Unit | Sigma-Aldrich | Cat# UFC510024 |
| Chromium™ Genome HT Library Kit & Gel Bead Kit v2 | 10X Genomics | Cat# PN-120261 |
| Chromium™ Genome Chip Kit v2 | 10X Genomics | Cat# PN-120257 |
| Chromium™ i7 Multiplex Kit | 10X Genomics | Cat# PN-120262 |
| QIAquick Gel Extraction Kit | Qiagen | Cat# 28706 |
| KAPA mRNA HyperPrep Kit | Roche | Cat# 08098123702 |

## CHEMICIALS, REAGENTS, AND PROBES

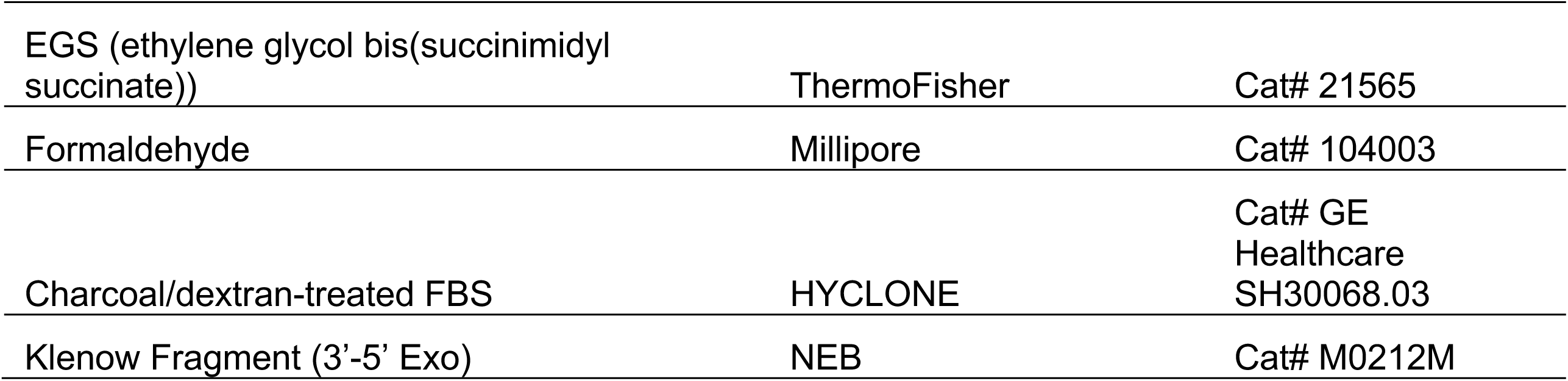
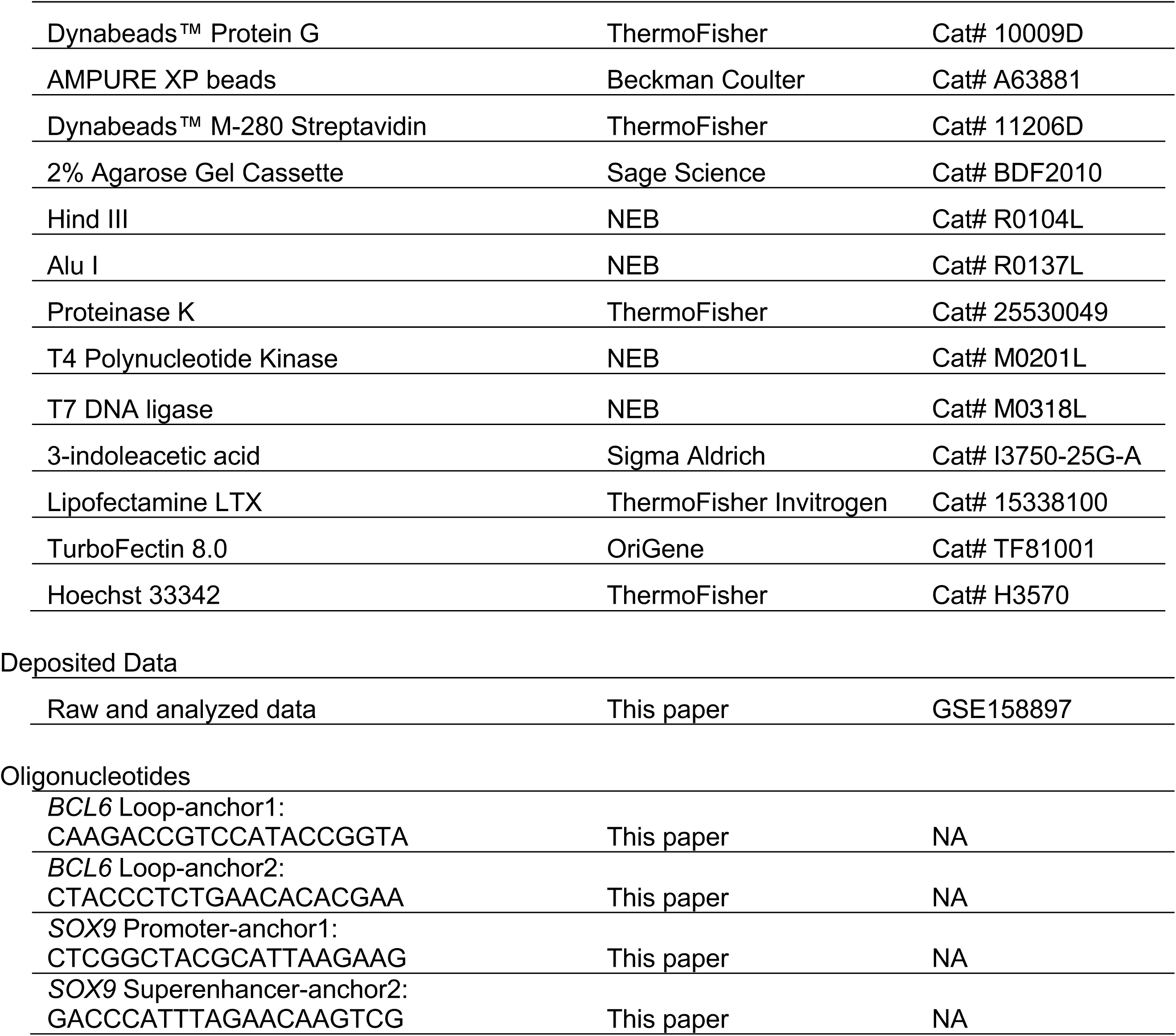

## EXPERIMENTAL MODEL AND SUBJECT DETAILS

### Cell Lines and Culture Conditions

Human GM12878 cells (Coriell Institute for Medical Research), B-lymphoblastoid cell line, were cultured in RPMI 1640 (ThermoFisher, A10491), supplemented with 15% fetal bovine serum (ThermoFisher, 10082147). These cells were cultured at 37 °C, 5% CO2, and ambient oxygen levels as described by the Coriell Institute of Medical Research.

These cells at exponential growth phase were harvested for chromatin preparation.

### Auxin-inducible degron system (AID) in HCT-116 cells

Human HCT-116-RAD21-mAID-mClover cells (HCT-116 RAD21-mAC) (Natsume et al., 2016) were cultured in McCoy’s 5A medium (ATCC, 30-2007) supplemented with 10% charcoal/dextran-treated FBS (HYCLONE, GE Healthcare SH30068.03), 2 mM L-glutamine, 100 U/ml penicillin, and 100ug/ml streptomycin at 37C with 5% CO_2_.

To induce rapid RAD21 protein degradation, 500uM indole-3-acetic acid (IAA; Sigma Aldrich, I3750-25G-A) dissolved in ethanol were added to cell culture and incubate for 6 hours, 9 hours, 12 hours or 24 hours.

For our ChIP-Seq and in situ ChIA-PET experiments on untreated cells and cells treated for 6 hours, 12 hours or 24 hours, medium was replaced with fresh medium (untreated) or medium with 500uM IAA. The cells were washed with DPBS, and then crosslinked with 1% formaldehyde for 10 minutes or 1% formaldehyde for 20 minutes followed by EGS fixation for another 45 minutes at room temperature. The cells were collected and stored at -80 °C for these experiments.

## METHOD DETAILS

### *In situ* ChIA-PET library preparation

ChIA-PET libraries with antibody against RNAPII, RAD21, SMC1, and CTCF were constructed using approximately 10^7^ input cells from GM12878 and HCT-116 cell cultures, following the *in situ* ChIA-PET protocol (Wang et al., 2021). The ChIA-PET libraries were sequenced with 150 bps long paired-end reads using NovaSeq 6000 instrument (Illumina).

### ChIP-Seq

We performed ChIP-Seq experiments for CTCF, RNAPII and RAD21 following the standard protocol with 1% formaldehyde for 10 minutes at room temperature. For WAPL and NIPBL ChIP-Seq experiments, we fixed the cells with 1% formaldehyde for 20 minutes followed by EGS fixation for another 45 minutes at room temperature, and then fixation was quenched by the addition of glycine to a final concentration of 125mM for 10 minutes. Ten million fixed cells were washed with PBS and stored at -80 °C until further processing or resuspended in 1 mL of RIPA buffer (10 mM Tris pH 7.6, 1 mM EDTA, 0.1% SDS, 0.1% sodium deoxycholate, 1% Triton X-100, Complete Mini EDTA free proteinase inhibitor (Roche)). Sonication was performed using Sonics sonifier at 4°C. Twenty microgram of respective antibody was incubated with 100 uL of Dynabeads Protein A (or G) overnight at 4°C or for at least 6 hours. Antibody-bound beads were added to 1 mL of sonicated chromatin, incubated at 4°C overnight, and then washed twice with RIPA buffer, twice with RIPA buffer containing 0.3M NaCl, twice with LiCl buffer (0.25 M LiCl, 0.5% Igepal-630, 0.5% sodium deoxycholate) and twice with TE (pH 8.0). Crosslinking was reversed by incubating the beads at 65°C for 4 hours in the presence of 1% SDS and 1 mg/mL Proteinase K. ChIP DNA was purified by ChIP DNA clean and concentrator column (Zymo research). Libraries were prepared using the tagment DNA Enzyme and Buffer Large Kit (Illumina), and size-selected libraries (300-500 bp) were sequenced by 1x 50bp single reads using NovaSeq 6000 instrument (Illumina). Antibodies used for ChIP-Seq are listed in the Key Resources Table.

### RNA-Seq

Total RNA from HCT-116-RAD21-mAID-mClover cells treated with IAA for 6, 9, 12 hours and those not treated with IAA (0 hour) was isolated by Trizol extraction. mRNA was then isolated and RNA-Seq library was prepared following the KAPA RNA HyperPrep Kit protocol (Roche, Cat. No: 08098123702). The RNA-seq libraies were sequenced by NovaSeq 6000 (Illumina) to either 2 x 50 bp or 2 x 150 bp reads.

### Protein factor-enriched chromatin sample preparation for ChIA-Drop

The overall factor-enriched chromatin sample preparation for ChIA-Drop followed the non-enriched protocol as descripted below except that the specific antibody (CTCF, RNAPII, RAD21, or SMC1) pull-down steps were added. The fragmented chromatin material (4000-6000 bp) was incubated with 20 µg of antibody bound on Dynabeads™ Protein G beads at 4 °C overnight with rotation. Antibody-enriched chromatin was released from Protein G beads by incubating with EB Buffer containing 1% SDS at 37 °C for 30 min with constant agitation. The elution supernatant was passed through Amicon Ultra-0.5 Centrifugal Filter with Ultracel-100 regenerated cellulose membrane (Millipore) to remove the remaining SDS. The quality control step was performed using ChIP-qPCR with the primers of specific transcription factor binding sites.

Qualified factor-enriched chromatin samples were proceeded for ChIA-Drop library construction. If not immediately used, the chromatin sample should be stored at -80 °C.

The overall chromatin sample preparation for ChIA-Drop is similar to that described in (Zheng et al., 2019), except for the following changes. First, 10 million cells were crosslinked with 1% formaldehyde at room temperature for 20 min followed by EGS fixation for another 45 minutes at room temperature, and quenched with 0.125 M Glycine (Promega) for 5 min, and then washed twice with DPBS. The crosslinked cells can be stored at -80 °C for later use or they can immediately proceed to cell/nuclei lysis. Second, the crosslinked cells were suspended in 1 ml of cell lysis buffer (50 mM HEPES-KOH pH 7.5, 150 mM NaCl, 1 mM EDTA, 1% Triton X-100, 0.1% Sodium Deoxycholate, 0.1% SDS, 1× Protease Inhibitor cocktail, Roche) and incubated at 4 °C for 1 hour with rotation. Then, the nuclei were isolated by centrifugation at 4 °C for 5 min at 2,500 relative centrifugal force (rcf). The nuclei pellet was suspended in 100 µl of 0.5% SDS and incubated at 62 °C for 5 min to permeabilize the nuclear membrane. After that, 285 µl of nuclease-free water and 25 µl of 20% triton X-100 were added for further incubation at 37 °C for 15 min to neutralize SDS. The permeabilized nuclei were then ready for in situ chromatin digestion. For the digestion step, 80 µl of nuclease-free water and 30 µl of HindIII (20 U/µl) were added to set up the reactions. The incubations took place at 37 °C overnight with constant agitation. Then the nuclei with digested chromatin materials were resuspended in 400 µl cell lysis buffer with 1× Protease Inhibitor cocktail and sheared by sonication to further fragment the chromatin fragments. The DNA size range of the chromatin fragments was generally 4000-6000 bp. The fragmented chromatin materials were filtered with Amicon Ultra-0.5 Centrifugal Filter with Ultracel-100 regenerated cellulose membrane (Millipore) to remove the remaining SDS and be concentrated, and ChIA-Drop library construction followed.

### ChIA-Drop library construction and sequencing

Each of the fragmented protein factor-enriched chromatin sample was mixed with 50 µg/ml of BSA (cat# B9000S, NEB) to prevent chromatin aggregation. To estimate concentration, an aliquot of the chromatin sample was taken and the DNA was then decrosslinked and purified (referred to as “pure DNA”) for quantification. Then, the chromatin sample was adjusted to a concentration of 0.5 ng/µl. The ChIA-Drop library construction and sequencing followed our previously published paper (Zheng et al., 2019) except the optimized time for the Gel bead in Emulsion (GEMs) amplification. In this study, the GEMs were subjected to a 30 °C isothermal incubation to amplify and barcode the chromatin DNA templates only for 3 hours instead of overnight.

### Casilio Experiment

#### Guide RNA design

To avoid interference of dCas9 binding on loop formation, we selected probes at least 5kb away from the ChIA-PET shores. A design window of 2kb was selected and overlapped with unique sites using the JACKIE pipeline v1.0 (Zhu et al., 2022) and off-target prediction software Cas-OFFinder v2.4 (Bae et al., 2014). The gRNAs were selected per design window minimizing, in order, 1-mismatch, 2-mismatch and 3-mismatch predicted off-target sites and the activity of gRNA is further optimized by selecting within 40∼60 GC%.

#### gRNA spacer sequences

Oligonucleotide used for gRNA spacer sequences are listed in the **Key Resources Table**.

#### Cloning

Clover fused with PUF RNA-binding domain were previously described: pAC1447 (Clover_PUFc) (Addgene #73689). Cloning of dCas9 expression plasmid (lenti-dCas9-Blast) and PUF9R_iRFP670 are described in (Clow et al., 2022). These PUF-fluorescent protein fusions contain nuclear localization signal (NLS) for their localization in the nucleus. gRNA spacer sequences were cloned into sgRNA-PBS expression vectors pCR8-sgRNA-15xPBSc or pCR8-sgRNA-15xPBS9R via an oligo-annealing protocol (Cong et al., 2013). All plasmids were subjected to restriction diagnostic tests and sequenced to ensure correct cloning.

#### Transfection

HCT116/RAD21-mAID/dCas9 cells were seeded at density of 60,000 cells/compartment in 35 mm 4-compartment CELLview™ cell culture dish the day before transfection. For each well, cells were transfected with 300 ng of each sgRNA-15xPBS plasmid and 40 ng of each fluorescent protein plasmid using 3.5 µL TurboFectin 8.0 (OriGene). Media was changed at 24 hours post-transfection.

#### Confocal microscopy

Imaging was performed at 42-52 hours post-transfection. Prior to imaging, cells were stained with 1.0 μg/ml Hoechst 33342 prepared in cell culture media for 30-60 minutes, followed by two media washes. Images were acquired with the Dragonfly High Speed Confocal Platform 505 (Andor) using an iXon EMCCD camera and a Leica HCX PL APO 40x/1.10 W CORR objective or Leica HC PL APO 63x/1.47NA OIL CORR TIRF objective mounted on a Leica DMi8 inverted microscope equipped with a live-cell environmental chamber (Okolab) at humidified 37°C and 5% CO_2_. Imaging mode was Confocal 25 μm. Hoechst images were acquired with a 200 mW solid state 405 nm laser and 450/50 nm BP emission filter. Clover images were acquired with a 150 mW solid state 488 nm laser and 525/50 nm BP emission filter. iRFP670 images were acquired with a 637 nm laser and 700/75 nm BP emission filter. Z-series covering the full nucleus was acquired at 0.27 μm step size for 40x objective, and 0.13 μm step size for 63x objective. For time-lapse imaging, the Z-series was acquired at 0.32 µm step size for 40x objective or 0.16-0.2 μm step size for 63x objective. Images are a maximum intensity projection of Z-series. Images were processed in Fusion software using ClearView-GPU deconvolution with the Robust (Iterative) algorithm (pre-sharpening 0, 5 iterations, and de-noising filter size 0.1). Linear adjustments in maximum and minimum levels were applied equally across the entire image to display only nuclear spot signals, and not background nuclear fluorescence.

#### Image analysis

For measuring two targeted genomic loci distance in time-lapse images, Fiji image analysis software was used (Schindelin, 2012). Z-series acquired at 0.32 μm step size for 40x objective, and 0.16-0.2 μm step size for 63x objective was used. If the nucleus drifted over time, the Correct 3D Drift plugin was used (Parslow, 2014). For segmenting and tracking spots, the TrackMate plugin was used (Tinevez, 2017). Blob diameter was set at 1.5-2.0 μm. For each channel, threshold was set to include two spots with the maximum intensity in the 3D volume of the nucleus. Simple LAP Tracker used 2 μm linking max distance, 2 μm gap closing max distance, and 2 gap closing max frame gap. Analysis produced “Spots in track statistics” file which was used to run python script to calculate 3D distances between spots, generate 3D tracks, and calculate speeds. To determine and correct for potential chromatic aberrations, Invitrogen Tetraspeck 0.1 µm Microspheres (ThermoFisher T7279) were imaged. On the Dragonfly High Speed Confocal Platform 505 (Andor), Z-series were acquired for both the Leica HCX PL APO 40x/1.10 W CORR objective and the Leica HC PL APO 63x/1.47NA OIL CORR TIRF objective. TrackMate was then used to localize the positions of the microspheres. iRFP670 chromatic aberrations in each X,Y,Z dimension were calculated relative to Clover, and corrected by offsetting the shifts in each X,Y,Z dimension as follows:

| Objective | Fluorescent Channel | X (mm) | Y (mm) | Z (mm) |
| --- | --- | --- | --- | --- |
| 40x | iRFP670 | -0.035 | -0.007 | +0.691 |
| 63x | iRFP670 | -0.001 | -0.012 | +0.147 |

### Bioinformatics

Software packages used:

- BASIC browser (Lee et al., 2020)
- Bedtools/2.27.0 (Quinlan and Hall, 2010)
- ChIA-PIPE (Lee et al., 2020)
- ChIA-DropBox (Tian et al., 2019)
- ChIA-View (Tian et al., 2019)
- Fiji/2.1.0 (Schindelin et al., 2012)
- Fiji Correct 3D Drift (Parslow et al., 2014)
- Fiji TrackMate/6.0.1 (Tinevez et al., 2017)
- HiCRep/1.11.0 (Yang et al., 2017)
- JACKIE (Zhu and Cheng, 2022)
- Juicer/1.6.2 (Durand et al.,2016)
- Juicebox/1.11.08 (Durand et al., 2016)
- MIA-Sig/0.1 (Kim et al., 2019)
- Python/3.6.6; 3.7.4

`o` numpy/1.18.1, pandas/1.0.1, matplotlib/3.1.3, pybedtools/0.8.1, pyBedGraph/0.5.42, scipy/1.3.2; 1.4.1, ipdb/0.12.3, Seaborn/0.10.0, deeptools/3.4.3, glob, functools, argparse, os, math, warnings
- R/3.4.0
- RStudio/1.1.442

### Previously published datasets used in this study

NIPBL ChIP-seq (Gene Expression Omnibus; accession GSM2443453), H3K27ac ChIP-seq (ENCODE data portal (Davis et al., 2018); accession ENCFF340JIF), H3K4me1 ChIP-seq (ENCODE data portal: accession ENCFF831ZH), RNA-seq (ENCODE data portal: accession ENCLB555AQG), GM12878 Hi-C (4DN data portal (Dekker et al., 2017): accession 4DNFI7J8BQ4P, 4DNFI1UEG1HD), list of GM12878 super-enhancers and constituents (Hnisz et al., 2013), HCT116 Hi-C (4DN data portal: accession 4DNFIP71EWXC, 4DNFIBIV8OUN), HCT116 0h Repli-seq (4DN data portal: accession 4DNFIR6ZS4LY, 4DNFID2WWTSC, 4DNFIH4B6I1S, 4DNFIFBWQ3QC, 4DNFI6FRVLDB, 4DNFINSRFNDX, 4DNFI3JLMX17, 4DNFIFS513KB, 4DNFIB697UQV, 4DNFIED8FHGM, 4DNFIJ88Z7MW, 4DNFI4FWA2X9, 4DNFIYQQ72X9, 4DNFIPWM5DS1, 4DNFIRBZUG62, 4DNFIQXJN452), HCT116 6h, Repli-seq (4DN data portal: accession 4DNFIXFHUPTI, 4DNFIX6NTFM4, 4DNFISBPS2ZV, 4DNFIMMM331D, 4DNFIPSBAONE, 4DNFIKESZXXD, 4DNFIZT1GRIL, 4DNFIHIHNQSN, 4DNFIQ573AKW, 4DNFI89FPVRX, 4DNFIST28EMP, 4DNFIA9QXIDF, 4DNFIRWK243V, 4DNFI6V9EXOM, 4DNFI1WWTVBY, 4DNFIL37I65A), list of HCT116 super-enhancers (https://asntech.org/dbsuper/download.php), GM12878 chromHMM states (https://hgsv.washington.edu/cgi-bin/hgFileUi?db=hg18&g=wgEncodeBroadHmm), and HCT116 chromHMM states (ENCODE data portal: accession ENCFF513PJK). CTCF *in situ* ChIA-PET data from HFFc6 (4DN data portal: accession 4DNESCQ7ZD21) and MCF10A (ENCODE data portal: accession ENCSR403ZYJ) cells. RNAPII ChIA-PET data, RNA-seq data and chromHMM states from H1 (4DN data portal: accession 4DNEXF93AC6Q; ENCODE data portal: accession ENCLB555AMA; UCSC Broad chromHMM), GM12878 (this study; ENCODE data portal: accession ENCLB555AQG; UCSC Broad chromHMM), K562 (ENCODE data portal: accession ENCSR880DSH, ENCLB555AKN; UCSC Broad chromHMM), HepG2 (ENCODE data portal: accession ENCSR857MYZ, ENCLB555AQD; UCSC Broad chromHMM), and MCF7 (ENCODE data portal: accession ENCSR059HDE, ENCLB555AQN, ENCFF506GEX). Gene expression profile in 76 human tissues from the Expression Atlas (Papatheodorou et al., 2019; https://www.ebi.ac.uk/gxa/home).

## QUANTIFICATION AND STATISTICAL ANALYSIS

### Definitions, Abbreviations, and Notations

Throughout the manuscript, the following set of definitions hold unless stated otherwise. Pearson’s r: Pearson’s correlation coefficient; Spearman’s r: Spearman’s correlation coefficient; TAD: topologically associating domain; ECDF: empirical cumulative distribution function; K-S test: Kolmogorov-Smirnov test; M-W test: Mann-Whitney U test; Mb: megabasepairs; kb: kilobasepairs. In all boxplots, the central line inside the box is the median, the edges of the box are the 25^th^ and 75^th^ percentiles, and whiskers extend to the most extreme data points not considered outliers. Seeds are set when generating random objects or numbers. The reference genome is hg38.

### ChIP-seq and ChIA-PET Data Processing

ChIP-seq reads were mapped to the hg38 genome with bwa v0.7.7 (bwa mem -t 20) and the resulting sam file was converted to bam file with samtools v1.5 (samtools view -S -b). Of the uniquely mapped and non-supplementary reads, only those with mapping quality greater than or equal to 30 are retained (samtools view -F 4 -F 2048 -q 30). De-duplicated reads (samtools rmdup -s) are used to generate bedgraph files via bedtools v2.26.0 (bedtools genomecov -ibam -bg). To filter out any mapping artefacts, ENCODE blacklist v2 regions are excluded from the coverage file (bedtools subtract). Both NIPBL and WAPL ChIP-seq data utilized GM12878 input control to call peaks on bam files via SPP (v1.13; R v3.2.1; Kharchenko et al., 2008) with z_thresh of 6.

ChIA-PET data were processed using ChIA-PIPE pipeline (Lee et al., 2020) on hg38 reference genome, ENCODE blacklist v2 regions, and peak caller SPP with input control. The resulting bedgraph file for binding intensity, bedpe files for loops, and bed files with significant peaks (z_thresh of 6) are used for downstream analyses. ChIA-PET terminologies are defined as follows. A loop has two anchors, left and right anchor, each with chromosome name, start, and end position. An anchor size is the distance between start and end position, anchor boundary ranges from the start position of the left anchor to the end position of the right anchor, and the loop span is defined as the distance between midpoint of the left anchor and midpoint of the right anchor. The PET count is the number of paired-end tags that link left and right anchors.

### ChIA-Drop Data Processing

The raw reads from all ChIA-Drop experiments were processed by the ChIA-DropBox pipeline (Tian et al., 2019) to map reads to the hg38 reference genome and using a parameter of 8 kilobasepairs (kb) to merge all reads therein to form a fragment. Two or more fragments with the same barcode within the same chromosome constitute a putative complex.

To resolve potential multiplets resulting from a droplet encapsulation of more than 1 chromatin complexes, we modified the distance test of the MIA-Sig algorithm (Kim et al., 2019) and implemented the entropy filter (entropy_filter.py v0.1) as follows. The upper distance threshold tau is a uniformly randomly selected integer between 1 megabasepairs (Mb) and 5 Mb. If the smallest fragment-to-fragment (F2F) distance is larger than tau, then the putative complex is separated into sub-complexes, each containing a single fragment. If largest F2F distance is smaller than or equal to tau, then the putative complex is kept as is. Otherwise, the smallest-to-largest sorted F2F distance is converted into a probability vector, which is used for computing normalized Shannon entropy (Shannon, 1948) iteratively using the first 1,2,…,n-1 entries, where n is the number of fragments; the distance corresponding to the smallest entropy is used as a threshold to separate a putative complex into multiple sub-complexes. The normalized Shannon entropy is defined as 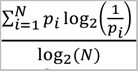 , where *N* is the number of outcomes and *p_i_* is the probability of *i*th event. The resulting union of putative complexes and sub-complexes with 2 or more fragments are referred to as entropy-filtered complexes, while those with only 1 fragment are entropy-filtered singletons.

Entropy-filtered complexes from non-enriched ChIA-Drop data are used in the downstream analyses. However, intra-chromosomal putative complexes from protein-enriched ChIA-Drop experiment have additional workflows to reflect protein-specific enrichment. The fragments mapped to the repetitive regions of the genome (UCSC simple repeats annotation file; Haeussler et al., 2019) are excluded via MIA-Sig (filter_repeat.py), and then go through the entropy filter. Next, the enrichment test of MIA-Sig (freq_enrich_sigtest.py v1.0) is run with parameters fdr_thresh=0.2, samp_size=500 on entropy-filtered complexes, and with fdr_thresh=0.2 and samp_size=100 on entropy-filtered singletons to obtain statistically significant complexes in protein-enriched regions. Fragments from complexes and singletons that pass the enrichment test are piled-up to generate protein binding intensity. Statistically significant complexes from the enrichment test are used in the downstream analyses. These processed and filtered ChIA-Drop data are collectively called chromatin complexes hereafter.

### Visualization of ChIA-PET and ChIA-Drop Data

The protein binding coverage bedgraph files from ChIP-seq and ChIA-PET data are visualized on the BASIC browser (Lee et al., 2020) as continuous integers on the y-axis denoting the number of reads piled-up. ChIA-PET loops are drawn as arcs with number of paired-end-tags (PETs) as heights indicating the strength of interaction between two genomic loci. To obtain similar number of loops for 3 ChIA-PET datasets from GM12878 cell line, the following criteria were applied before the visualization: 1) 172,635 CTCF loops with >=6 PETs and at least 1 anchor overlapping a peak; 2) 161,911 cohesin loops with >=9 PETs and at least 1 anchor overlapping a peak; 3) 147,737 RNAPII loops with >=5 PETs and at least 1 anchor overlapping a peak.

Both ChIA-PET and ChIA-Drop interaction data can be visualized on Juicebox as 2D contact maps by using Juicer software v1.7.5 to generate *.hic files of resolutions ranging from 1 kb to 2.5 Mb. The multiplex interactions from ChIA-Drop data are converted to pairwise interactions by simply enumerating over all pairs for each entropy-filtered complexes (but without applying the enrichment test). ChIA-Drop chromatin complexes are converted to subrds files, which are inputs to ChIA-View (RStudio v1.1.442) for the visualization of intricate multiplex chromatin interactions. Two modes are available on ChIA-View: cluster view for large segments of binned complexes, and fragment view for a collection of fragments connected with a horizontal line to constitute a chromatin complex.

To facilitate the single-molecule visualization and analyses of bulk ChIA-PET dataset, the *bsorted.pairs.gz files encoding an individual paired-tag interactions are treated as a list of putative complexes akin to those resulting from a ChIA-DropBox pipeline. Specifically, we extend the midpoint genomic coordinate of each pair (columns 3 and 5 of the pairs file) by 250 bps both directions and only retain the intrachromosomal pairs with the midpoints separated by more than 8 kb (as means of treating those less than 8kb as self-ligated pairs). We then assigned a tentative “GEM ID” by concatenating a library ID with “-100-”, an integer counting the pairs+100000000, and “-HEA-7-4-sub-1-1”. The final *region file is a list of chromosome, start, end, 2 (denoting the number of fragments), and GEM ID.

### Reproducibility assessment of ChIA-Drop datasets

To assess the reproducibility of ChIA-Drop experiments, we computed stratum-adjusted correlation coefficient (SCC) by running HiCRep (v1.11.0) (Yang et al., 2017) on 3 CTCF, 2 RNAPII, 1 SMC1A, 2 RAD21, and cohesin (combining 1 SMC1A and 2 RAD21) ChIA-Drop datasets and respective CTCF, RNAPII, cohesin (combining 1 SMC1A and 1 RAD21) ChIA-PET data. Parameters for HiCRep are 100 kb resolution, maximum distance of 2 Mb, and smoothing factor of 3, with raw matrices for enriched ChIA-Drop and ChIA-PET data. To account for the varying sequencing depths, contact matrices were downsampled to 800,050 contacts (e.g., the number of complexes in the minimum dataset, CTCF dataset 3) when comparing individual and pooled ChIA-Drop datasets. However, SCCs between pooled ChIA-Drop and ChIA-PET data were computed without downsampling due to similar level of high depth in sequencing between the pairs.

### Identifying confident CTCF loops for GM12878 cell line

From 1,181,427 loops with PET count >=3 in the GM12878 CTCF ChIA-PET data, highly confident CTCF loops were selected based on a few filtering criteria. The first set of filters require that 1) both left and right anchor sizes are less than 10 kb; 2) the distance between anchor boundaries are greater than 8 kb; 3) the loop span is between 50 kb and 3 Mb. These 609,634 loops satisfying the criteria are further filtered with additional requirements that the maximum binding intensity within both anchors exceed 200 read counts and that the PET count is greater than 9. Total 13,549 loops passing these stringent criteria are referred to as a set of ‘confident CTCF loops’ in the downstream analyses.

### Calling CTCF-mediated chromatin domains and CTCF binding motifs

CTCF binding motifs were called in Tang et al., 2015, using the software package STORM (Search Tool for Occurrences of Regulatory Motifs) and the resulting 22,648 motifs are overlapped with loops using bedtools. Of the 13,549 loops from GM12878 CTCF ChIA-PET data, 6,385 loops had convergent motifs at their anchor sites, where left anchor overlapped with a forward (+ or >) CTCF motif and the right anchor overlapped with a reverse (- or <) motif. Following the approach described in Tang et al., contiguous convergent loops were merged to form a larger domain, yielding 2,289 CTCF-mediated chromatin domains (or simply referred to as ‘domains’). Median domain size is around 478 kb. To facilitate downstream analyses, motifs were first extended by 4 kb both directions. If two or more motifs with same orientation overlapped after the extension, they were merged to form a ‘multi-motif’. If two motifs with different orientation overlapped, then the one with higher CTCF ChIA-PET binding was picked unless one of the motifs reside in the domain boundary, in which case it received the priority. This merging scheme resulted in 20,251 motifs for statistical analyses in the following sections including the HCT116 cell line data analysis.

### Comparing cohesin and RNAPII at interaction loci among enhancers and promoters

To extract the interactions between regulatory elements mediated by cohesin and RNAPII, we first defined a set of elements by using peaks called from CTCF, cohesin, RNAPII ChIA-PET and NIPBL ChIP-seq experiments performed in this study. Similar to CTCF domains, we defined RAIDs (RNAPII-associated interaction domain) as follows: from 147737 RNAPII ChIA-PET loops with PET counts >=5 and having at least one anchor with peak support and loop span less than 700kb, merge any overlapping loops and obtain 1213 RAIDs that have more than 5 loops merged.

Of 6270 cohesin peaks not overlapping CTCF peaks, 5400 overlapped with RNAPII peaks. Further filtering by NIPBL peak overlap and merging any overlapping peaks (bedtools merge) yielded 4236 peaks annotated with chromHMM states (bedtools groupby -o max): active promoter (n=932), weak promoter (n=113), enhancer (n=2378), a hybrid of enhancer and promoter (n=429), others (n=384). There are total 3852 enhancer or promoter peaks (E/P). Of 147737 RNAPII ChIA-PET loops with PET count >=5 and peak support, 12654 have both anchors overlapping E/P; from 161911 cohesin ChIA-PET loops with PET count >= 9 and peak support, 7978 have both anchors overlapping E/P. These loop anchors are adjusted to only the overlapping portion of E/P. By further requiring loops to be within the same RAID with more than 1 E/P and with loop span greater than 100kb, 1706 loops remained. Promoter definition in this section encompasses ‘active promoter’, ‘weak promoter’, ‘hybrid of enhancer and promoter’.

For each of these 1706 loops, ChIA-Drop complexes are sorted: 1) from the first element to the second element, 2) second element to the first element, 3) overlapping both first and second elements, 4) from the first element to the left by the same loop span, 5) from the second element to the right by the same loop span. The number of complexes in the first three categories are plotted for CTCF, cohesin and RNAPII in a boxplot, with an additional scatterplot to compare cohesin and RNAPII. A subset of 1667 loops with at least 1 CTCF, RNAPII, and cohesin complexes in any of the 5 categories are retained for the following analysis. The Jensen-Shannon divergence (JS div.) is computed for a pair of probability vectors, each derived by dividing the number of complexes in 5 categories by the total number of complexes; JS div. close to 1 implies dissimilar, and towards 0 is considered to be similar. Boxplots of JS div for RNAPII vs. cohesin, RNAPII vs. CTCF, and cohesin vs. CTCF are presented.

### Characterizing multiplexity of ChIA-Drop complexes with respect to the expected distribution

A unique feature of ChIA-Drop experiment is its ability to capture multiplex (2 or more) interactions. We sought to quantify the multiplexity of ChIA-Drop complexes with respect to the promoters and enhancers (for RNAPII). Genome-wide coverage is as follows: 49,004,649 bps of promoter chromHMM states; and 136,227,664 bps of enhancer states. Given that the total size of the hg38 reference genome is 3,088,286,401 bp, P(promoter)=0.0159, P(enhancer|no promoter)=0.0448. Then the expected number of complexes with *k* elements (promoter) is 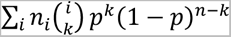, where *i* ranges from 2 to maximum fragment number, *n_i_* is the observed number of complexes with *i* fragments, and *p* = 0.0159. Applying this formula to the RNAPII ChIA-Drop complexes, we obtain the expected number of complexes with 0, 1, 2, 3, or >=4 motifs. The number of RNAPII complexes with 0,1,2,3, or >=4 promoters are: 1086441, 1073944, 229688, 8695, 964 (observed); 2316500, 82282, 941, 8, 1 (expected). Of 1086441 complexes with no promoter, 738877, 307255, 38854, 1339, 116 complexes have 0,1,2,3, or >=4 enhancers, respectively (observed), whereas expected counts are 1069072, 17282, 87, 0, 0 (expected). These pairs of observed and expected distributions are compared against each other via the two-sided K-S test.

### Aggregating contact regions

It is informative to aggregate over many 2D contact regions of interest to obtain genome-wide patterns. First, .hic files are converted into .cool files (Abdennur and Mirny, 2019) with 5 kb resolution using hic2cool v0.8.3 (hic2cool convert -r 5000). The observed raw counts are normalized by the average counts over all bins of the same distance to account for the fact that genomic loci in close proximity are more likely to interact by chance than those far apart. These expected counts are averaged over a set of regions that are greater than ‘min_size’ and less than ‘max_apart’, where ‘min_size’ and ‘max_apart’ are user-defined parameters. When aggregating cohesin and CTCF ChIA-Drop contact maps, those in the reverse motifs or promoters are flipped along the 45 degree diagonal line and are plotted together with the forward motifs.

### Calling ChIA-Drop peaks and generating binding heatmap

One of the benefits of the MIA-Sig enrichment test is the de-noising of bedgraph binding tracks by retaining only the fragments in high binding thereby removing spurious data. Therefore, the fragments of significant singletons and complexes from enrichment test are piled up to generate binding coverage, which is used to call peaks in CTCF, cohesin, and RNAPII ChIA-Drop data. By scanning through the genome with 500 bp non-overlapping windows, those with binding intensity greater than a threshold are called as enriched bins, which are then merged if two enriched bins are within 3 kb. Thresholds are 70, 400, and 90 for CTCF, cohesin, and RNAPII, respectively, which have been selected based on genome-wide average binding intensity of all 500 bp bins. This approach yielded 19117, 26652, and 14608 peaks for CTCF, cohesin, and RNAPII ChIA-Drop data, respectively. Common peaks between CTCF and cohesin are identified via bedtools (Quinlan and Hall, 2010) v2.27.1 (bedtools intersect -wa) and factor specific peaks by bedtools intersect -wo.

### Identifying cohesin loading regions and sorting complexes

As means of characterizing the process by which cohesin loads and translocates to the anchor sites, 10239 cohesin-specific peaks are further filtered by the criteria that NIPBL binding (of GSM2443453 downloaded track) is greater than 5 counts, yielding 5,467 regions. A total 3,014 convergent loops include 1 or more of these cohesin loading regions; since convergent loops may overlap (e.g., a bigger loop enclosing a smaller loop), some loading regions are recorded for two different convergent loop. These 3014 loops enclosed 8777 loading regions.

### Characterizing RNAPII-associated loading and anchoring regions by cohesin

One of the objectives is to characterize the RNAPII interactions with respect to cohesin at loading sites and CTCF anchor sites, further categorized by the transcription direction and motif orientation.

To define the RNAPII-associated anchor regions, RNAPII ChIA-PET peaks overlapping cohesin ChIA-PET peaks (bedtools intersect -u) are retained if they are within 25 kb of CTCF ChIA-PET peaks overlapping the binding motif (bedtools window -w 25000). These 5521 peaks are then annotated by the closest 50 genes and their expression levels using ENCODE RNA-seq data (ENCFF879KFK); only those within 5 kb to the promoter with TPM (Transcripts Per Kilobase Million) greater than 0.5 and gene body length > 5 kb are considered to be ‘TSS’ and all others are ‘non-TSS’. If a peak overlaps multiple ‘TSS’, then those with highest TPM are selected. Subsequently, the peaks are further categorized into: 1) forward motif and forward TSS (n=423); 2) forward motif and reverse TSS (n=339); 3) forward motif and non-TSS (n=1779); 4) reverse motif and forward TSS (n=456); 5) reverse motif and reverse TSS (n=418); 6) reverse motif and non-TSS (n=2106).

Likewise, RNAPII ChIA-PET peaks overlapping cohesin ChIA-PET peaks (bedtools intersect -u) are kept if they do not overlap CTCF ChIA-PET peaks nor do they overlap CTCF motif (bedtools intersect -v); finally, those overlapping NIPBL ChIP-seq binding peaks (generated in this study) amount to 3638 “RNAPII at cohesin loading sites”. Using the same criteria described above for the anchor regions, TSS is assigned based on the highest TPM on each of the forward and reverse orientations (if any). If there are two genes transcribed in both directions, the one with more than twice TPM is selected and the peak is considered to have transcriptions to the both directions (‘both TSS’) if TPMs do not differ by more than a factor of 2. Finally, loading sites are considered to have: 1) forward TSS (n=556), 2) reverse TSS (n=423), 3) non-TSS (n=363), 4) both TSS (n=19).

Each of these peaks have been extended to the both sides by the gene body length if they are associated with TSS, and by 150 kb both directions if they are non-TSS. Finally, only the regions larger than 100 kb are retained for aggregation plots and statistics on directionality of RNAPII and cohesin complexes. In particular, the ChIA-Drop complexes in these regions are extracted if they overlap the peak, which are next sorted from the peak to the left and to the right ends of the region. The number of complexes to the left and to the right are recorded for: TSS in concordance with motif (n=264) and TSS in discordance with motif (n=199), TSS at loading (n=445) and non-TSS at loading (n=2640).

### Super-enhancer (SE) annotation and intra-SE contacts

A list of 257 super-enhancers with 1640 SE constituents in GM12878 cell line was downloaded from Hnisz et al., 2013. 158 SEs greater than 33 kb are selected and are extended by its span to the both directions, and 1000 random regions of 300 kb are sampled via bedtools v 2.27.0 (bedtools random -l 300000 -n 1000 -seed 1234). Aggregating the 2D contact maps in these regions are performed with distance normalization. Likewise, the number of complexes with 2 or more fragments within each of these regions are recorded for CTCF, cohesin, and RNAPII ChIA-Drop data.

### Constructing SE graphs and computing node degrees

By taking the union of RNAPII ChIA-PET and ChIA-Drop peaks and merging those within 500 bps and concatenating with super-enhancer constituents, we obtained 34107 peaks annotated with chromHMM state and gene expression level if it is within a promoter region. 71,906 RNAPII ChIA-PET loops with PET count >=5 and both anchors overlapping these peaks are used as reference interaction map involving super-enhancers and other regulatory elements. To identify the target gene(s) of SE, loops with one anchor overlapping SE and another anchor overlapping a promoter of a gene with TPM > 1 of span larger than 150 kb and smaller than 6 Mb, with at least 1 RNAPII complex are candidates, among which the one with highest TPM is selected as target gene promoter (TP; simplified as P in the main text and figures). As a result, there are 188 SE-TP pairs.

We constructed a graph for each of these 188 pairs by taking all elements from SE to TP as nodes and the number of ChIA-Drop complexes connecting a pair of nodes as weighted edges. A node degree is defined as the number of weighted edges incident to the node, e.g., the sum of all ChIA-Drop complexes connecting a given node to any other nodes. The normalized degree is defined as a node degree divided by the sum of all node degrees (i.e., 2*|E|, where E is the set of all weighted edges). The number of direct contact denotes the number of ChIA-Drop complexes connecting SE and TP, whereas the indirect contact is the sum of interactions between SE and intermediary enhancer or intermediary promoter—all but TP. In other words, the number of direct and indirect contacts sum up to the degree of TP. There are 75 pairs in which the number of direct contact is greater than or equal to the indirect contact, 80 pairs with direct < indirect; 33 pairs have another SE along the path to TP.

These analyses are first performed for RNAPII ChIA-Drop data, and repeated for cohesin ChIA-Drop data on the same 188 pairs. SE graphs are constructed and plotted via the networkx v1.11 python package (Hagberg et al., 2008).

### Processing HCT116 ChIP-seq, ChIA-PET, and Hi-C data and obtaining related files

The HCT116 cells are treated with auxin for RAD21 depletion at timepoints 0hrs (i.e., no treatment), 6h, 9h, and 12h and were subsequently subject for CTCF, RAD21, and RNAPII ChIP-seq experiments.

ChIP-seq data were processed as described earlier, resulting in mapped reads (bam), peaks and coverage (in bedgraph / bigwig). For obtaining genome-wide binding pattern changes over timepoints and to ensure that RAD21 is indeed depleted, the binding coverage is aggregated at peak regions as follows. First, the bam file is converted into a coverage file by deeptools command bamCoverage with options --binSize 60 --normalizeUsing RPGC --effectiveGenomeSize 2913022398 -- ignoreForNormalization chrY chrM --extendReads --outFileFormat bigwig. The aggregate binding intensity at 32252 CTCF ChIP-seq peaks, 27334 RAD21 ChIP-seq peaks, and 13999 RNAPII ChIP-seq peaks at 0 hours are plotted for 0h, 6h, 9h, and 12h by deeptools functions computeMatrix and plotHeatmap.

After confirming that 6 hours of auxin treatment is sufficient to deplete RAD21, we generated non-enriched ChIA-PET (referred to as Hi-C), CTCF ChIA-PET and RNAPII ChIA-PET data at 0h and 6h and the replicates are combined for subsequent analyses. The 2D contact maps in *.hic format are converted into *.cool format with 5kb resolution and the values are extracted at 6385 convergent loops with coolpup.py parameters --nshifts 10 --unbalanced --coverage_norm. Final plots are generated with plotpup.py options --enrichment 3 --scale linear --vmin 0 --vmax 8 --cmap seismic.

Subsequent analyses focus on CTCF ChIA-PET and RNAPII ChIA-PET data generated at 0h and 6h. One of the outputs of ChIA-PIPE is the *bsorted.pairs.cis file which encodes all intra-chromosomal paired-end-tags. The relative density of the log10 of the distance between the pairs is plotted for both CTCF and RNAPII ChIA-PET data with x-axis limited to be between 3 and 6.5; python function gaussian_kde is imported from scipy.stats.kde.

The super-enhancer annotation for HCT116 was available only in hg19 reference genome, so we lifted over the coordinates via liftOver tool and obtained 2588 super-enhancer constituents and 387 super-enhancers in hg38. There were 16 chromHMM states for HCT116, which is further simplified to ‘Enh’ if only overlapping enhancers, ‘Tss’ if only overlapping Tss, ‘Enh/Tss’ if overlapping both enhancers and Tss, and ‘Other’ otherwise.

### HCT116 RNA-seq data processing and analysis of differential gene and gene ontology

Raw fastq files from each of the RNA-seq experiments are processed as follows. The reads are first aligned to the reference genome hg38 via STAR (v 2.5.3) aligner with options --outSAMtype BAM and SortedByCoordinate. Subsequent bam files are de-duplicated with picard using MarkDuplicates command and reads with mapping quality less than 30 are filtered out using samtools (v 1.5) view command.

Finally, bam files are converted into strand-specific bedgraph files by bedtools (v 2.27.0) bamtobed and genomecov commands; these bedgraph files are visualized on the BASIC browser tracks with green denoting the positive forward strand and blue representing the negative reverse strand.

There are RNA-seq data from 0h, 6h, 9h, and 12h timepoints in HCT116 cells treated with auxin, each with two replicates. To quantify the transcripts, each of the bam files are subject to featureCounts function in the subread package with respect to the gtf file version Homo_sapiens.GRCh38.94.gtf. The resulting count files are concatenated such that each row is a gene, and the columns denote a gene ID, length, and counts of 0h rep1, 0h rep2, 6h rep1, …, 12h rep2. This information is stored in a *tsv file. For differential gene analysis, only 0h and 6h data of both replicates are retained and are read through DESeq2 library in R. Performing DESeqDataSetFromMatrix function followed by DESeq resulted in p-values and adjusted p-values assigned to each gene. Using the threshold of 0.05, there were 2474 genes with adjusted p-value less than 0.05 and a negative log2 fold-change (i.e., significant reduction in expression) and 1568 genes with adjusted p-value less than 0.05 and a positive log2 fold-change. An additional criteria is enforced by first computing the transcripts per million (TPM) for each gene for each sample (normalizing the transcript counts by gene length, then dividing it by the sum of normalized counts and multiplying by 1000000). The average TPM for each condition is the mean TPM of replicate 1 and replicate 2. Finally, there are 361 up-regulated genes with adjusted p-value < 0.05 and log2fc > 1 and average TPM at 6h > 20; 356 down-regulated genes with adjusted p-value < 0.05 and log2fc < -1 and average TPM at 0h > 20; 5391 unchanged genes with p-value >= 0.05 and -1 < log2fc < 1 and average TPM at 0h > 20; the other 10315 genes did not meet any of these criteria. The volcano plot labeled up- regulated genes as red, down-regulated genes as blue, unchanged genes as green, and others in grey.

Our RNA-seq data are compared to the publicly available PRO-seq data before (0h) and after (6h) depleting RAD21 in HCT116 cells (Rao et al., 2017). As authors have also performed DESeq2 to identify 235 up-regulated genes and 64 down-regulated genes, we used it as a reference set to compare the log2(fold-change) via boxplots for both PRO-seq and RNA-seq. p-values were computed by Mann-Whitney U test. For the 4761 unchanged genes, a scatterplot of log2 fold-change from RNA-seq and PRO-seq is plotted with colors denoting the high density of datapoints (red highest, blue lowest).

The RNAPII ChIA-PET binding peak intensity at promoters (+/- 2kb from TSS) of down-regulated and up-regulated genes are recorded as maximum value of 0h and 6h (combined replicates) data and are plotted in a scatterplot. An alternative aggregation of peaks are plotted with deeptools for a region extending 25 kb from the gene promoter.

As an input to the gene ontology (GO) analysis (http://great.stanford.edu), the gene coordinates in the each category of down-regulated, up-regulated and unchanged genes are extracted from Homo_sapiens.GRCh38.94.gtf and run on the web server with a whole genome background and human: GRCh38 (UCSC hg38, Dec. 2013) assembly.

### Identification and characterization of RAD21-dependent and RAD21-independent CTCF loops

Using two replicates of 0h HCT116 CTCF ChIA-PET data (library IDs LHH0157 and LHH0172) and two replicates of 6h data (library IDs LHH0158 and LHH0173), the differential loops are identified as follows. The 0hr combined data had 131361 peaks called by ChIA-PIPE, of which 34108 had maximum peak value greater than 240 (median x 1.8). After merging peaks within 3kb, 28407 peaks remained. Similarly, 6hr data had 127816 peaks, which were filled to 31333 peaks greater than a threshold of 200 and finally yielded 25495 peaks. Taking the union (bedtools intersect -v) of 28407 peaks in 0hr and 25495 peaks in 6hr amounted to 30071 union peaks. Each of the loops with PET count >= 3 is annotated with its anchor(s) overlapping a union peak, if any: 415937 loops 0h rep1, 579746 loops 0h rep2, 273821 loops 6h rep1, 245261 loops 6h rep2. Now each loop is labeled as unique peak ID of the left anchor and that of the right anchor and PET counts are collated for each sample (rows are loop labels and columns are PET counts for 0h rep1, 0h rep2, 6h rep1, 6h rep2). This table of 90434 loops with nonzero counts is treated as a feature count and undergoes DESeq2 with false discovery rate of 0.2: 521 loops with adjusted p-value < 0.2 and log2fc > 0.5 had significant increase in loop strengths, 18897 loops with adjusted p-value < 0.2 and log2fc < -0.5 had significant decrease, and 4407 loops with adjusted p-value >= 0.2 and -0.5 < log2fc < 0.5 remained unaffected. Since the last two classes make up the majority of the population, we subsequently focus on the 18897 RAD21-dependent loops (as these loops form strongly only in the presence of RAD21) and 4407 RAD21-independent loops that do not significantly change the loop strengths.

Of 90434 total loops, 42083 loops (46.5%) had both anchors overlapping CTCF binding motif, 39019 loops (43.1%) had one anchor overlapping CTCF motif, and 9332 loops (10.3%) had no anchor overlapping CTCF motif. 42083 loops are further categorized into 19644 convergent (><) loops, 8742 right tandem (>>) loops, 8787 left tandem (<<) loops, and 4910 divergent (<>) loops. The numbers for 18897 RAD21-dependent loops were 12249 loops (64.8%) with both anchors overlapping motif, 5883 with only one anchor, and 765 with none. There were 8974 convergent, 1451 right tandem, 1638 left tandem, and 186 divergent loops. Interestingly, only 34.9% (n=1537) of the 4407 RAD21-independent loops had both anchors overlapping motif, with a fewer convergent loops (n=190) than right tandem (n=480), left tandem (n=449) and divergent (n=418) loops. 2241 loops had one anchor overlapping motif, and 629 loops had none.

The 0h (LHH0157) and 6h (LHH0158) CTCF data are represented as single-molecule data as described above, and are sorted at 6385 convergent loops. As expected, 0h data had higher proportion of complete looping (median = 0.07) than 6h data (median = 0.0).

A particular interest is to characterize the small RAD21-independent loops around CTCF binding sites. The union peak of 0h and 6h CTCF data are further filtered to overlap CTCF motif, have binding intensity greater than 1000 in both 0h and 6h data, and to not have another CTCF motif within +/- 25 kb. There were 2168 such peaks, which is referred to as ‘strong CTCF binding’ in this section. To compute the stalling rate, the region files after MIA-Sig enrichment test FDR 0.2 for each of the 0h and 6h data are intersected with 2168 strong CTCF binding motifs extended by 4kb both directions (bedtools intersect - wao). For each loop, the stalled complexes (within 25kb from motif) and extruding complexes (distance greater than 25kb and less than 3 Mb following the motif orientation) are recorded. The stalling rate is then defined as the number of stalled complexes divided by the sum of the number of stalled complexes and extruding complexes. Stalling rate is lower in 0h (median =0.20) than in 6h (median=0.45) data.

Finally, CTCF complexes are aggregated at these 2168 strong CTCF binding sites extended by 25 kb both directions in arc plot representing the number of complexes following the motif orientation vs. those against the motif.

### Composition of RAD21-dependent and RAD21-independent RNAPII loops

A similar procedure from CTCF ChIA-PET differential loop analysis is applied to RNAPII ChIA-PET loops. At the peak level, HCT116 0h RNAPII ChIA-PET data (combined LHH0170, LHH0159, LHH0159V) had 129566 original peaks, 28679 peaks greater than intensity of 200, and 19563 merged within 3kb.

Likewise, HCT116 6h RNAPII ChIA-PET data (LHH0171, LHH0160, LHH0160V) had 128199 peaks, 27700 peaks greater than 240, and 18060 3kb-merged peaks. These two groups had a union (bedtools intersect -v) of 21667 peaks and each of the following loops with PET count greater than or equal to 3 are annotated with anchors overlapping union peaks: 421918 loops 0h rep1, 230530 loops 0h rep2, 281940 loops 6h rep1, 196384 loops 6h rep2. A count table for RNAPII ChIA-PET data is created for each pair of unique loop and corresponding PET counts in each of the 4 samples (69066 rows and 5 columns). Using the same threshold, out of 69066 loops with nonzero total read count, 113 loops with adjusted p-value < 0.2 and log2fc > 0.5 had strengthened, 1201 loops with adjusted p-value < 0.2 and log2fc < -0.5 had weakened, and 4605 with adjusted p-value >= 0.2 and -0.5 < log2fc < 0.5 were unaffected. As a result, there are 1201 RAD21-dependent RNAPII loops and 4605 RAD21-independent RNAPII loops.

Of 69066 total loops, 7460 loops had both anchors overlapping CTCF motifs in convergent (n=2630), right tandem (n=1783), left tandem (n=1796) and divergent (n=1251) orientation. 37823 loops did not overlap any CTCF motifs, but instead were categorized into promoter-promoter (P-P; n=9235), enhancer-promoter (E-P; n=11224) and enhancer-enhancer (E-E; n=6066) loops. The RAD21-dependent RNAPII loops (n=1201) were composed of 306 convergent, 55 right tandem, 62 left tandem, and 9 divergent loops. 340 loops were devoid of CTCF motifs and constituted 41 P-P, 150 E-P, and 88 E-E loops. Only a small portion (8.6%; n=396) of 4605 RAD21-independent loops were associated with CTCF, with a small number (n=79) of convergent loops, and the rest of right tandem (n=125), left tandem (n=134) and divergent (n=58) loops. The majority (58.1%; n=2674) of the RAD21-independent loops were CTCF-free and were mostly P-P (n=857), E-P (n=706), and E-E (n=493) loops.

### Comparison of RAD21-dependent E-P loops and RAD21-independent P-P loops

After observing that the majority of RAD21-dependent loops (n=1201) connecting enhancers to promoters and that RAD21-independent loops (n=4605) were connecting active gene promoters, we sought to define RAD21-dependent E-P loops and RAD21-independent P-P loops regardless of the presence or absence of CTCF binding motifs. Recall that differential loops are labeled with pairs of unique peak ID that overlap loop anchors. Parsing it out, if the peak overlaps a gene promoter (defined as +/- 1kb) with its expression TPM greater than 20, then it is considered a promoter ‘P’. Of those not considered a promoter, if the peak has chromHMM state annotated as ‘Enh’ or ‘Enh/Tss’, then the anchor is labeled as an enhancer ‘E’. This approach resulted in 548 RAD21-dependent E-P loops and 1586 RAD21- independent P-P loops, from which only those with the distance between the end of left anchor and the start of righ anchor greater than 10kb are retained, resulting in 536 and 1218 loops, respectively.

The genes in each category are subject to the quantification of cell-type-specificity. To do so, we downloaded the gene expression data (TPM) in 76 human tissues and further annotated the number of tissues for which a gene is expressed—with TPM greater than 1 considered as being expressed, for each gene (Expression Atlas, Papatheodorou et al., 2019). There were 1839 unique genes in 1218 RAD21- independent P-P loops, of which 1704 had been annotated in the Expression Atlas. Likewise, 361 unique genes in 536 RAD21-dependent E-P loops had a subset of genes (n=332) annotated in the database.

The relative density function of the number of tissues for these 1704 genes in RAD21-independent P-P loops (median # of tissues 74 and mean 52) and 332 genes in RAD21-dependent E-P loops (median # of tissues 34 and mean 38.1) are plotted.

Using the HCT116 super-enhancer annotation file, the super-enhancer to target gene promoter pair is assigned from the union of RNAPII 0h and 6h loops following the same rule applied to assigning GM12878 super-enhancer to target gene promoter. Further constraining the SE-TP pair to have a PET count of greater than 10 and loop span greater than 100kb, there were 465 Prom-SE contacts and log10 of PET counts +1 for 0h and 6h are plotted in a boxplot (median = 26 for 0h, median = 5 for 6h).

To characterize the two categories of loops with respect to other cell lines, RNAPII ChIA-PET, RNA-seq, and chromHMM data from five cell lines H1, GM12878, K562, HepG2, and MCF7 are downloaded from the sources specified above. The RNAPII ChIA-PET loops have been filtered to contain only those with both anchors supported by peaks annotated as enhancers or promoters and with PET counts greater than or equal to 5. We next incorporate the gene expression data from these five cell lines and HCT116 0h RNA-seq data by recording TPM of each gene in each cell line, constituting 1839 genes (rows) by 6 cell lines (columns) for genes RAD21-independent P-P loops, and 361 by 6 table for RAD21- dependent E-P loops. The spearman’s correlation between a vector gene expression for all pairs of 6 cell line data is computed and are clustered with python’s seabourn clustermap function and parameters row_cluster=True, col_cluster=True, vmin=0, vmax=1, cmap=’seismic’ and a default parameter of using Euclidean distance as a metric. A similar approach is taken for 536 RAD21-dependent E-P loops and 1812 RAD21-independent P-P loops, where the total number of complexes (=‘left-ongoing’ + ‘right- ongoing’ + ‘complete looping’) are recorded by sorting the ChIA-PET single-molecule complexes at the loop anchors/peaks. The spearman’s correlation between a vector of number of complexes for all pairs of 6 cell line data is computed and clustered via hierarchical clustering with the same function and parameters as genes.

### Analysis of 16-stage Repli-seq data in HCT116 cells

The 16-stage Repli-seq data for 0h (with RAD21) and 6h (without RAD21) HCT116 cells (Emerson et al., 2022) were downloaded from the 4DN data portal. In particular, the raw bigwig files for each of P02 to P17 are downloaded (accession number provided above) and visualized using the same scale of y-axis. For each of the 361 up-regulated, 356 down-regulated, and 5391 unchanged genes, the median replication signal within the gene body is computed and plotted as a line plot for the 0h and 6h Repli-seq data. As a control, 20000 random regions of size 30 kb are also plotted.

---

**Figure S1:**
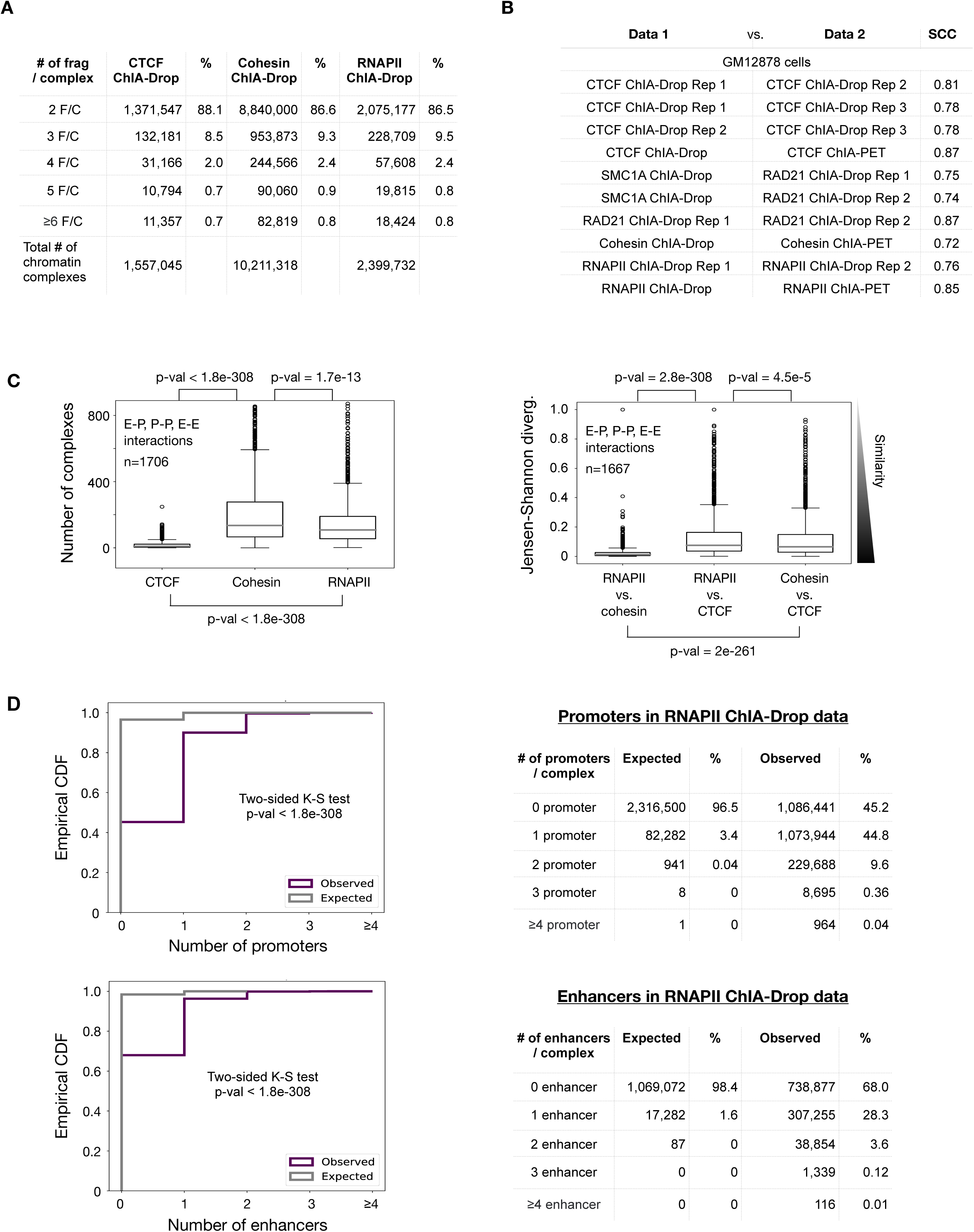
ChIA-Drop data for mapping chromatin interactions mediated by CTCF, cohesin, and RNAPII. **(A)** A table of ChIP-enriched CTCF, cohesin, and RNA Polymerase II (RNAPII) ChIA-Drop chromatin complexes by the number of fragments per complex (F/C). **(B)** Stratum-adjusted correlation coefficients (SCC) between all datasets of ChIP-enriched CTCF, RAD21, SMC1A, and RNAPII ChIA-Drop experiments and their replicates. R1, R2, and R3 denote replicates 1, 2, or 3 of a given experiment, respectively. SCC between ChIP-enriched ChIA-Drop and corresponding ChIA-PET data are also computed. **(C)** Boxplots for quantifications of transcriptional chromatin interactions. Left panel: number of chromatin complexes in CTCF, cohesin, and RNAPII ChIA-Drop data at 1,706 loop loci characterized in Figure 1D. Right panel: the Jensen-Shannon divergence of pairs of the datasets between RNAPII, cohesin, and CTCF ChIA-Drop (see **Methods**). p-values are computed from the two-sided Mann-Whitney U test. **(D)** Left panel: empirical cumulative distribution function (ECDF) of the observed (purple) and expected (grey) number of RNAPII ChIA-Drop complexes with 0, 1, 2, 3, and ≥ 4 promoters (top) and those with 0, 1, 2, 3, and ≥ 4 enhancers (bottom) are plotted. Right panel: the numbers and percentages of RNAPII ChIA-Drop complexes with promoters (top) and enhancers (bottom) that were expected and observed.

**Figure S2:**
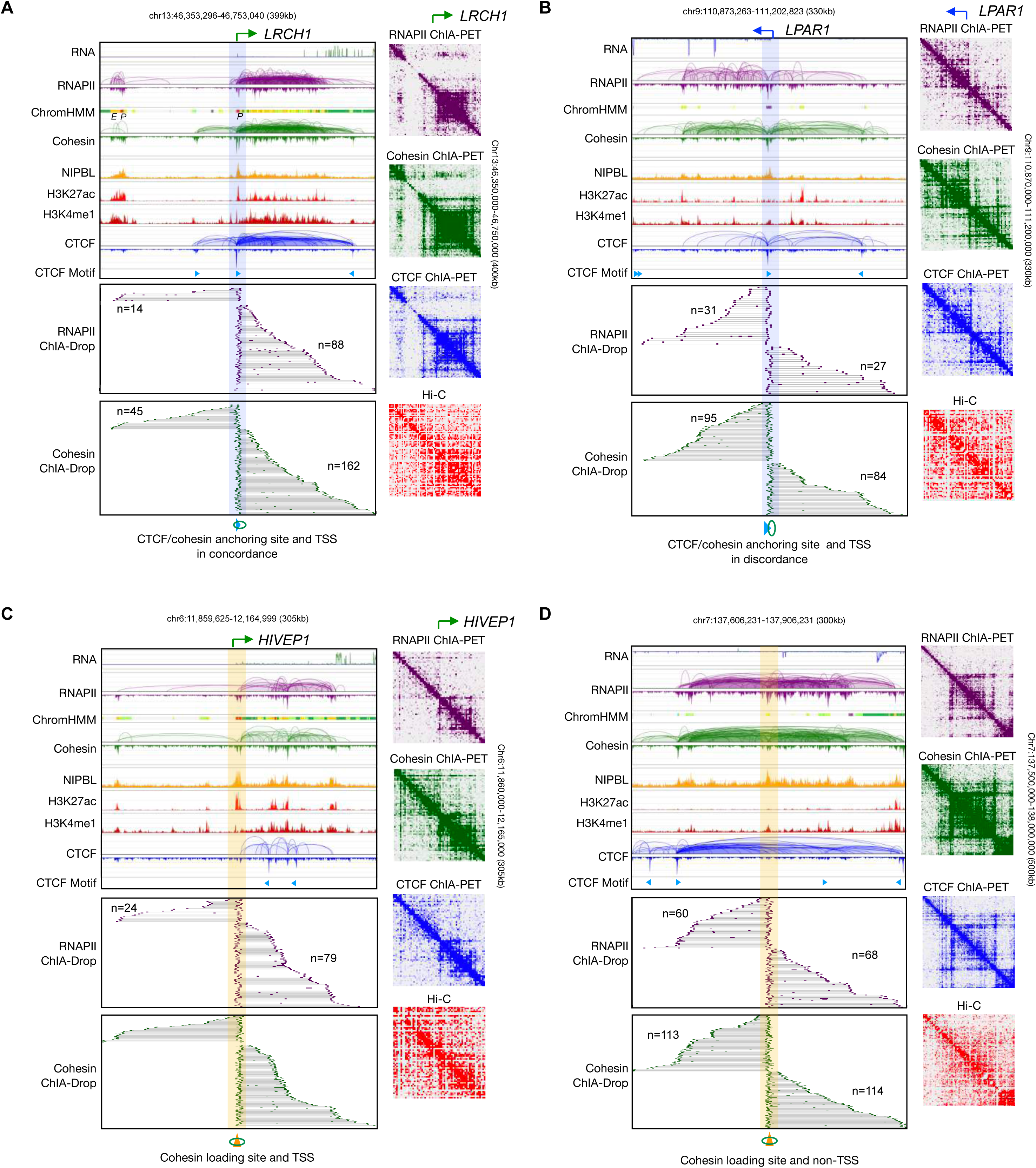
Transcriptional loops mediated by RNAPII and cohesin through concerted efforts. **(A)** RNAPII binding site coinciding with the TSS of *LRCH1* at CTCF/cohesin anchoring site (highlighted in blue) in concordance with CTCF binding motif. Data tracks are RNA-seq, RNAPII ChIA-PET loops/peaks along with chromHMM states, cohesin ChIA-PET loops/peaks, NIPBL ChIP-seq, H3K27ac ChIP-seq, H3K4me1 ChIP-seq, CTCF ChIA-PET loops/peaks, and CTCF binding motifs, followed by the sorted views of RNAPII and cohesin ChIA-Drop complexes centered at the TSS. The 2D contact maps of RNAPII ChIA-PET, cohesin ChIA-PET, CTCF ChIA-PET, and Hi-C are accompanied on the right. **(B)** Similar to panel **A**, but at the TSS of *LPAR1* in discordance with CTCF/cohesin anchoring site. **(C)** Similar to panel **A**, but presenting RNAPII binding site and TSS of *HIVEP1* at NIPBL binding/cohesin loading site (highlighted in yellow). **(D)** Similar to panel **A**, but at the NIPBL binding/cohesin loading site not overlapping with TSS of any gene.

**Figure S3:**
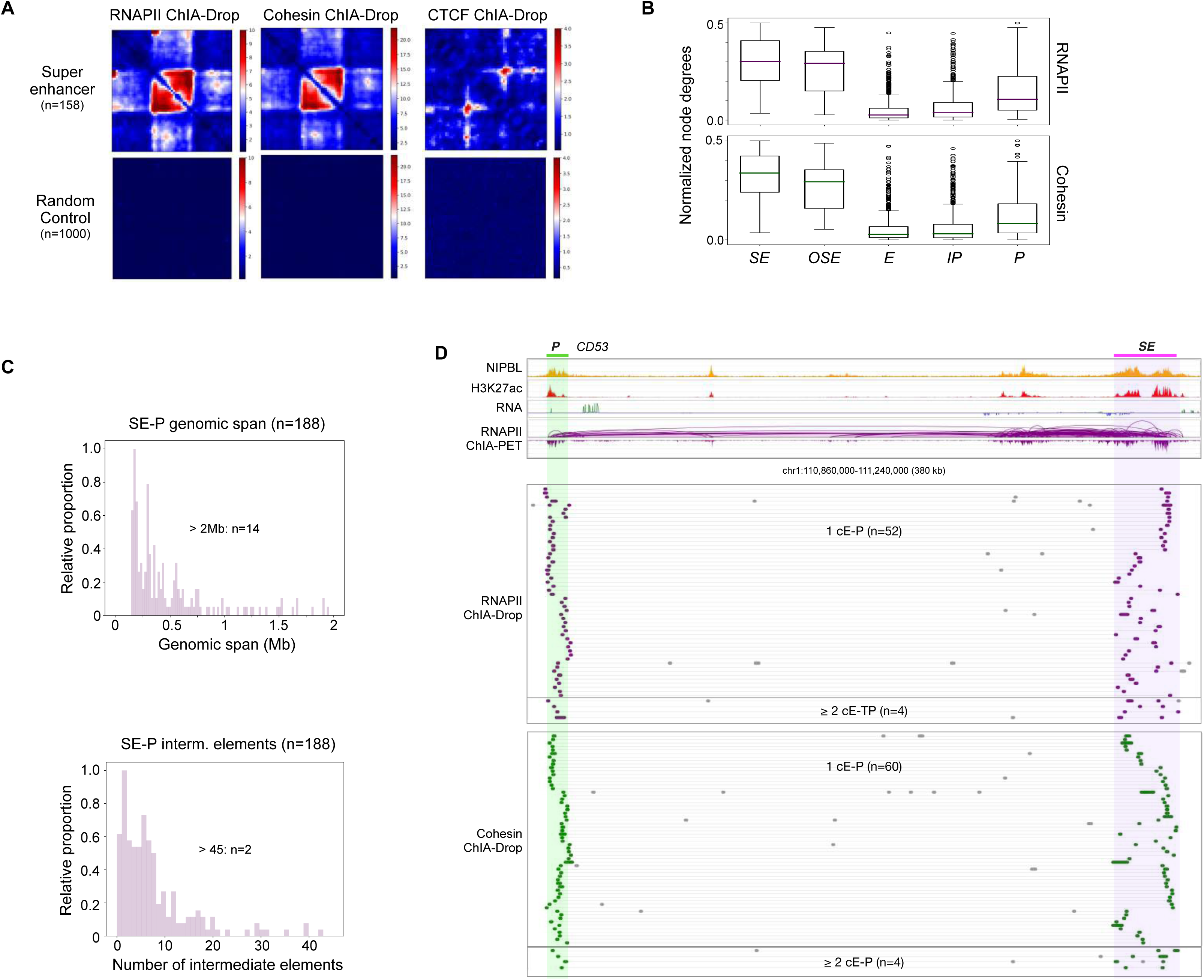
Multiplex transcriptional chromatin interactions involving super-enhancers. **(A)** Aggregation of 2D pairwise contacts of RNAPII, cohesin, and CTCF ChIA-Drop data at super-enhancer regions and at random regions as negative controls. **(B)** Boxplots of normalized node degrees of SEs, other SE along the path (*OSE*), enhancers (*E*), intermediary promoters (*IP*), and target gene promoters (*P*) are plotted for the 188 SE-P pairs in RNAPII and cohesin ChIA-Drop data. **(C)** A histogram of genomic span of 188 SE-P structures, of which 14 are larger than 2 Mb (top). The number of intermediate elements between SE and P are also plotted (bottom). **(D)** An example of SE-P interactions at the *CD53* gene locus. Top tracks are ChIP-seq of NIPBL and H3K27ac, RNA-seq, and RNAPII ChIA-PET. Below are the fragment views of RNAPII (purple) and cohesin (green) ChIA-Drop complexes showing single-molecule resolution of SE-*CD53* interactions involving one constituent enhancer (1 cE-P) and multiple constituent enhancer (³ 2 cE-P), where n denotes the number of chromatin complexes.

**Figure S4:**
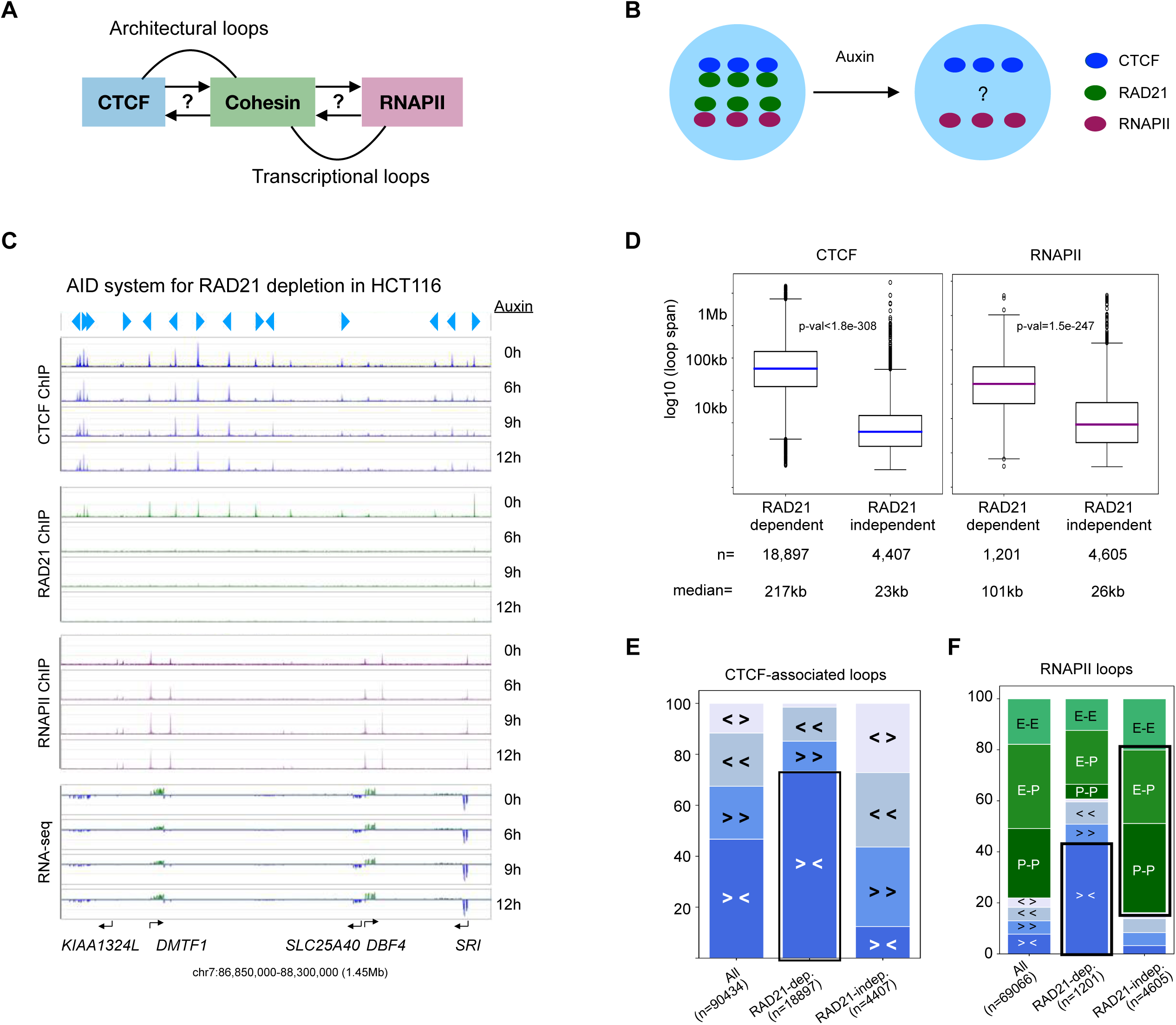
Effects of RAD21 depletion on long- and short-range chromatin interactions mediated by CTCF and RNAPII. (**A**) A diagram questioning the causal roles of cohesin in forming architectural loops with CTCF and transcriptional loops with RNAPII. **(B)** A schematic of the Auxin Inducible Degron (AID) tagged cell line HCT116-RAD21-mAC for auxin (IAA)-inducible degradation of RAD21, a subunit of cohesin. **(C)** An example browser view of ChIP- seq data of CTCF, RAD21, and RNAPII and RNA-seq data in HCT116 cells tagged with Auxin-inducible Degron AID (HCT116-RAD21-mAC) with auxin (IAA) treatment for 0, 6, 9, and 12 hours. Light blue arrows indicate CTCF binding motif and orientations. **(D)** Boxplots of chromatin loop span in the categories of ‘RAD21-dependent’ (i.e., reduced loop strengths) and ‘RAD21-independent’ (unchanged) loops in CTCF and RNAPII ChIA-PET data (see **Methods**), where n denotes the number of loops in each category and median loop span recorded below. p-values are from the two-sided Mann-Whitney U test. **(E)** Segmented bar charts for the proportions of CTCF-associated chromatin loops in ‘All’, ‘RAD21-dependent’, and ‘RAD21-independent’ HCT116 loops. CTCF loops with binding motifs in 4 categories: convergent (‘> <’), right tandem (‘> >’), left tandem (‘< <’), divergent (‘< >’). **(F)** Segmented bar charts for the proportions of RNAPII-associated chromatin loops in ‘All’, ‘RAD21-dependent’, and ‘RAD21- independent’ HCT116 loops. RNAPII loops are first characterized by CTCF binding motifs as convergent (‘> <’), right tandem (‘> >’), left tandem (‘< <’), divergent (‘< >’), and the rest of the CTCF-free loops are further categorized as promoter-promoter (‘P-P’), enhancer-promoter (‘E-P’), and enhancer-enhancer (‘E-E’) loops.

**Figure S5:**
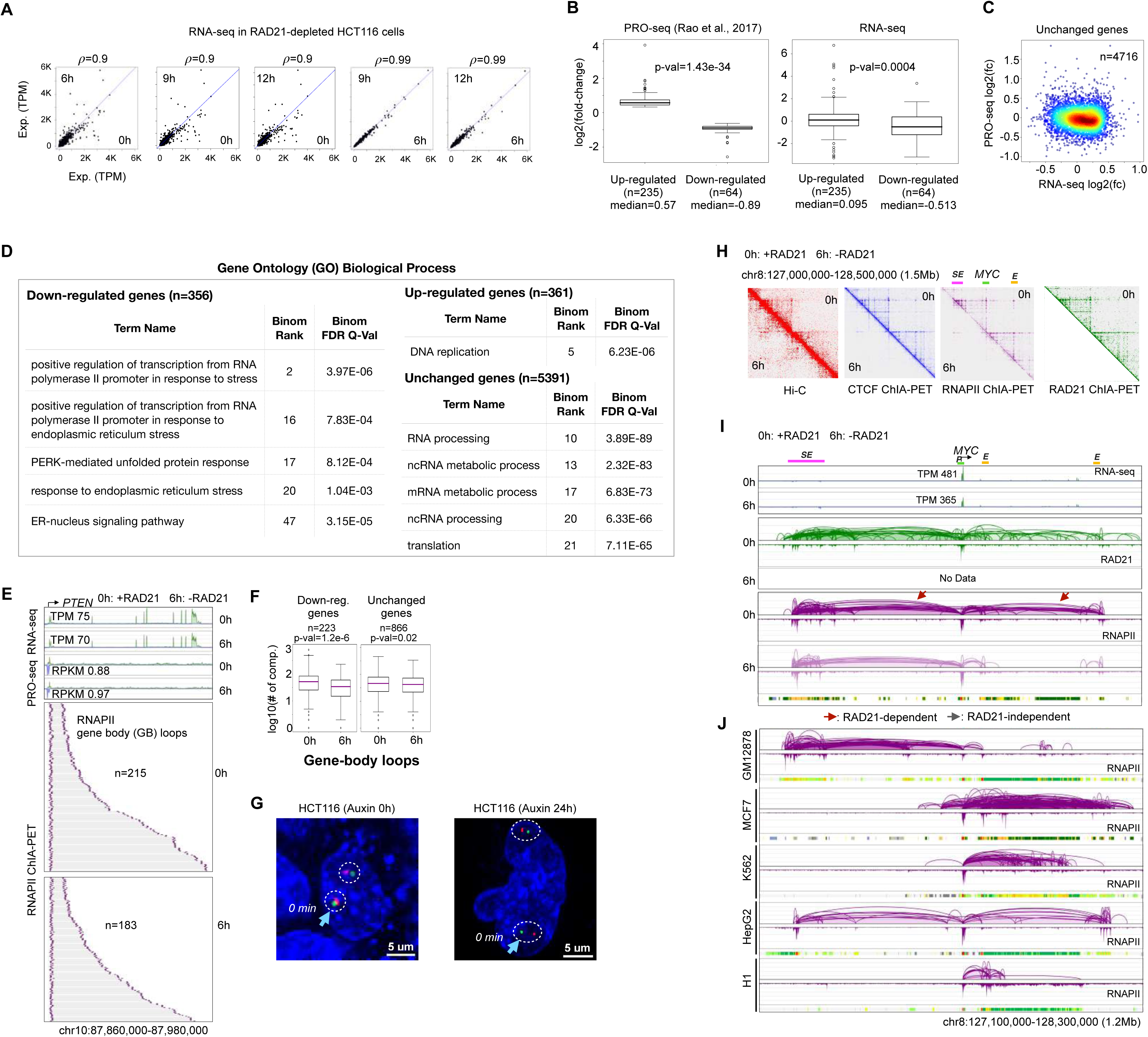
Functional roles of RAD21-dependent super-enhancer to promoter loop in transcription. **(A)** Scatter plots of TPM (transcripts per kilobase million) of genes from RNA-seq with various timepoints of auxin treatment. : Pearson’s correlation coefficient. **(B)** Boxplots of log2(fold-change) of gene expression before and after depleting RAD21 of up-regulated and down-regulated genes from PRO-seq (left) and RNA-seq (right) datasets; up-regulated and down-regulated genes were defined by PRO-seq (Rao et al., 2017). **(C)** A scatterplot of log2(fold-change) of gene expression between RNA- seq and PRO-seq data for unchanged genes, with colors denoting the density of data points. **(D)** Gene Ontology terms enriched in down-regulated, up-regulated, and unchanged genes. **(E)** Gene body loops of RNAPII ChIA-PET data before (0h; h: hours) and after (6h) depleting RAD21 are sorted from the promoter of an unchanged *PTEN* gene towards the transcription end site in the same forward orientation as the gene transcription; n denotes the number of chromatin complexes. TPM (transcript per kilobase million) from RNA-seq (this study) and RPKM (reads per kilobase million) from PRO-seq (Rao et al., 2017) are also recorded. **(F)** Boxplots of log10 of number of complexes in the gene-body loops before (0h) and after (6h) depleting RAD21, plotted separately for down-regulated genes and unchanged genes. **(G)** Representative Casilio images of *SOX9*-SE loop anchors (two pairs of probes in circles per nucleus) in control HCT116-RAD21-mAC cells (Auxin 0h) and cells with 24 hours of auxin treatment for RAD21-degradation (Auxin 24h). The pair with light blue arrow in each image is further tracked in Figure 5G. Scale bars, 5 µm. **(H)** 2D contact maps of data before (0h) and after (6h) depleting RAD21 at a 1.5 Mb region centered around *MYC* gene, super-enhancer (SE) and enhancer (E) mapped via Hi-C, CTCF ChIA-PET, RNAPII ChIA-PET, and RAD21 ChIA-PET. Only 0h data exist for RAD21 ChIA-PET. **(I)** In a large chromatin domain (1.2 Mb) harboring *MYC* gene and associated regulatory elements super-enhancer (SE), enhancers (*E*) and promoter (*P*) demarcated by ChromHMM, tracks of RNA-seq, cohesin ChIA-PET, and RNAPII ChIA-PET loops/peaks in HCT116 cell line before (0h) and after (6h) depleting RAD21 are shown. TPM: transcripts per kilobase million. Reduced loops are marked by red arrows. **(J)** In the same region as panel **I**, the RNAPII ChIA-PET loops and peaks and chromHMM states are also shown for 5 other cell lines: GM12878, MCF7, K562, HepG2, H1.

**Figure S6:**
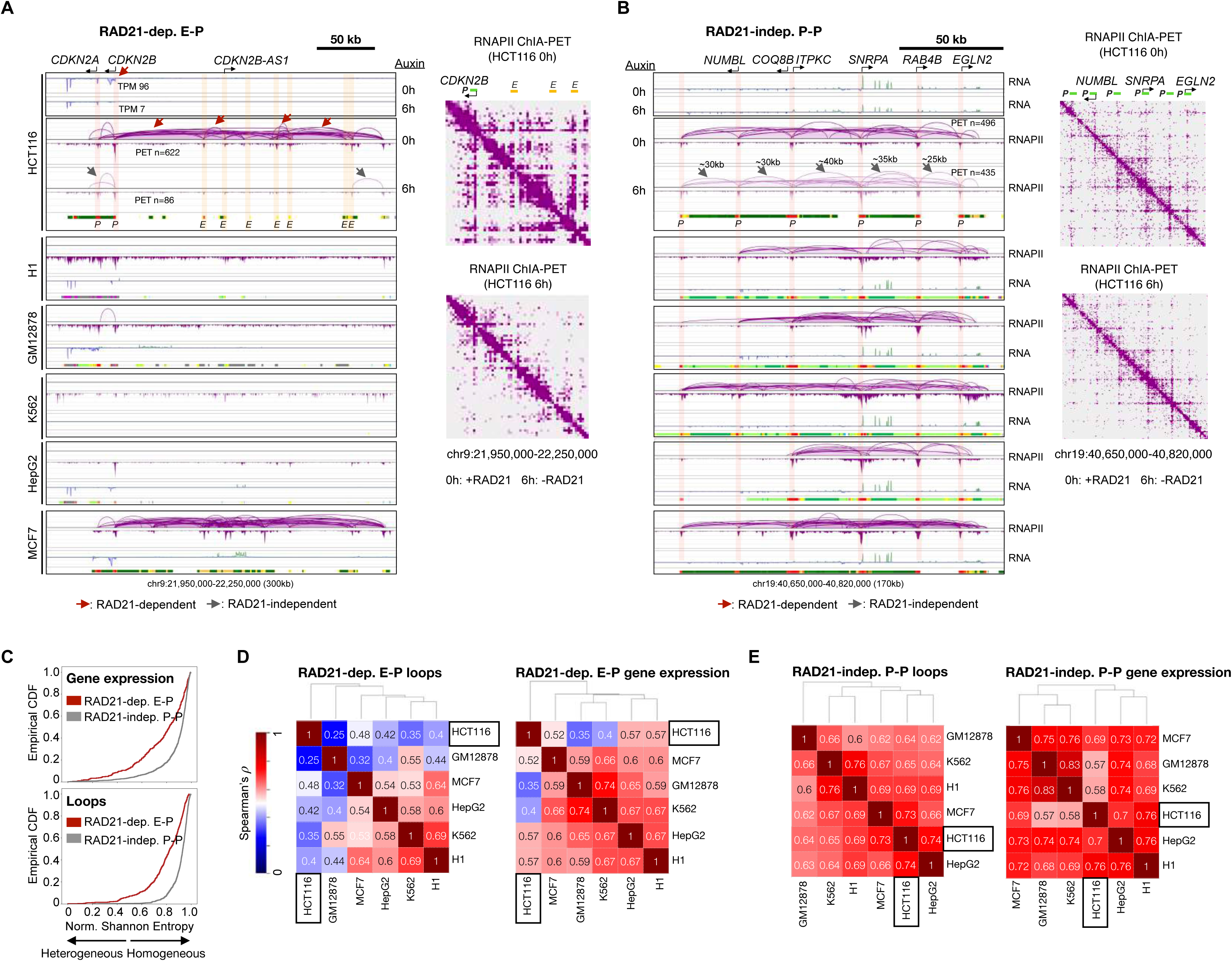
Distinct roles of RAD21-dependent E-P and RAD21-independent P-P RNAPII loops in gene regulation. **(A)** A 300 kb region including a down-regulated *CDKN2B* gene, where RNAPII ChIA-PET, RNA-seq, and ChomHMM chromatin states are shown for HCT116 and 5 other cell lines: H1, GM12878, K562, HepG2, and MCF7. 2D contact maps of RNAPII ChIA-PET in HCT116 cells before (0h) and after (6h) RAD21 depletion are also shown. **(B)** A 170 kb region encompasses RAD21-independent RNAPII loops connecting promoters (P) of active genes. Annotations are consistent with those in panel **A**. **(C)** An empirical cumulative distribution function (CDF) of the normalized Shannon entropy (see **Methods**) quantified over 6 cell lines gene expression (top panel) and chromatin interaction strengths (bottom panel) involved in RAD21-dependent enhancer-promoter (E-P) and RAD21-independent promoter-promoter (P-P) loops. **(D)** The Spearman’s correlation coefficient between the genomic profiles between all pairs of 6 cell lines, clustered via hierarchical clustering (see **Methods**). The left panel characterizes loop strengths of RAD21-dependent enhancer-promoter (E-P) loops and the right panel includes genes involved in these loops. **(E)** A similar plot as panel **H** for the RAD21-independent promoter-promoter (P-P) interactions and associated genes therein.

**Figure S7:**
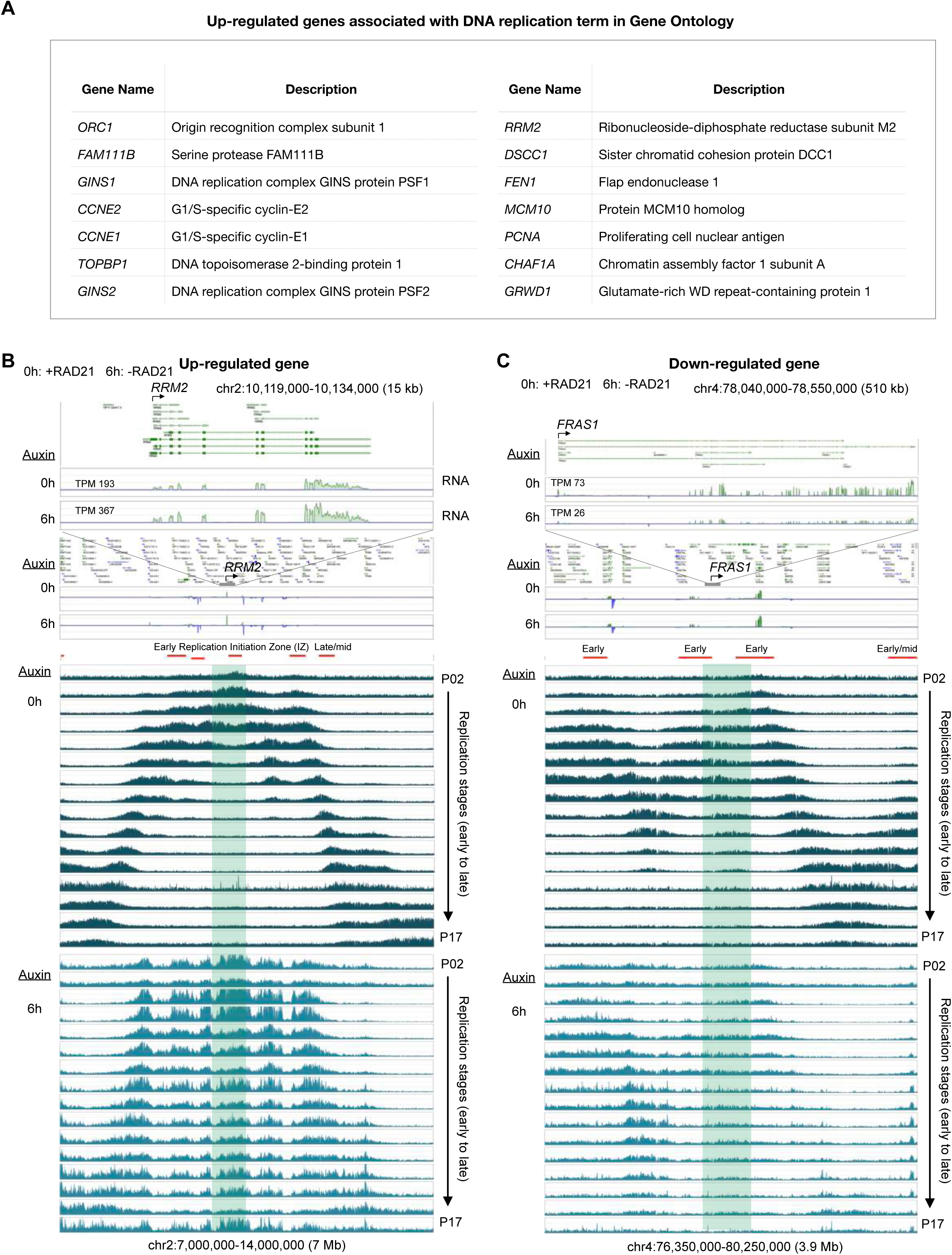
DNA replication signal patterns in differential genes, and proposed model. **(A)** A list of 14 genes identified to be associated with DNA replication in Gene Ontology of up-regulated genes, along with their descriptions of functions. **(B)** An example of an up-regulated gene *RRM2* along with RNA-seq signal (top panel) and 16-stage Repli-seq (Emerson et al., 2022) signal from early P02 to late P17 stages in HCT116 cells before (0h; middle panel) and after (6h; bottom panel) RAD21 depletion. Red bars are replication initiation zones identified using 0h data in Emerson et al. with specific labels early or late/mid replication. **(C)** Similar to panel **B**, but for a down-regulated *FRAS1* gene.

## SUPPLEMENTAL INFORMATION TITLES AND LEGENDS

**Supplemental Table S1.** Summary of mapping and imaging data. Related to **Figure 1**.

**Supplemental Video S1.** Time-lapse video of *BCL6* loop of pair N13 auxin 0h. Scale bar = 1 μm. Related to **Figure 4**.

**Supplemental Video S2.** Time-lapse video of *BCL6* loop of pair N24 RAD21 degraded auxin 24h. Scale bar = 1 μm. Related to **Figure 4**.

**Supplemental Video S3.** Time-lapse video of *SOX9* loop of auxin 0h. Scale bar = 1 μm. Related to **Figure 5**.

**Supplemental Video S4.** Time-lapse video of *SOX9* loop of RAD21 degraded auxin 24h. Scale bar = 1 μm. Related to **Figure 5**.

## REFERENCES

Abdennur, N., and Mirny, L.A. (2019). Cooler: scalable storage for Hi-C data and other genomically labeled arrays. Bioinformatics, 36, 311–316.

Allahyar, A., Vermeulen, C., Bouwman, B.A., Krijger, P.H., Verstegen, M.J., Geeven, G., van Kranenburg, M., Pieterse, M., Straver, R., Haarhuis, J.H. and Jalink, K. (2018). Enhancer hubs and loop collisions identified from single-allele topologies. Nature genetics, 50(8), 1151–1160.

Arrastia, M.V., Jachowicz, J.W., Ollikainen, N., Curtis, M.S., Lai, C., Quinodoz, S.A., Selck, D.A., Ismagilov, R.F. and Guttman, M. (2022). Single-cell measurement of higher-order 3D genome organization with scSPRITE. Nature biotechnology, 40(1), 64–73.

Bae, S., Park, J., and Kim, J.-S. (2014). Cas-OFFinder: a fast and versatile algorithm that searches for potential off-target sites of Cas9 RNA-guided endonucleases. Bioinformatics, 30, 1473–1475.

Banigan, E.J., van den Berg, A.A., Brandão, H.B., Marko, J.F., and Mirny, L.A. (2020). Chromosome organization by one-sided and two-sided loop extrusion. eLife, 9, e53558.

Banigan, E.J., Tang, W., van den Berg, A.A., Stocsits, R.R., Wutz, G., Brandão, H.B., Busslinger, G.A., Peters, J.M. and Mirny, L.A. (2023). Transcription shapes 3D chromatin organization by interacting with loop extrusion. Proceedings of the National Academy of Sciences, 120(11), e2210480120.

Beagrie, R.A., Scialdone, A., Schueler, M., Kraemer, D.C., Chotalia, M., Xie, S.Q., Barbieri, M., de Santiago, I., Lavitas, L.M., Branco, M.R. and Fraser, J., (2017). Complex multi-enhancer contacts captured by genome architecture mapping. Nature, 543(7646), 519–524.

Beagrie, R.A., Thieme, C.J., Annunziatella, C., Baugher, C., Zhang, Y., Schueler, M., Kukalev, A., Kempfer, R., Chiariello, A.M., Bianco, S. and Li, Y. (2023). Multiplex-GAM: genome-wide identification of chromatin contacts yields insights overlooked by Hi-C. Nature Methods, 1-11.

Bensaude, O. (2011). Inhibiting eukaryotic transcription. Which compound to choose? How to evaluate its activity? Which compound to choose? How to evaluate its activity?. Transcription, 2(3), 103–108.

Bintu, B., Mateo, L. J., Su, J. H., Sinnott-Armstrong, N. A., Parker, M., Kinrot, S., Yamaya, K., Boettiger, A.N. & Zhuang, X. (2018). Super-resolution chromatin tracing reveals domains and cooperative interactions in single cells. Science, 362(6413), eaau1783.

Braccioli, L. and de Wit, E. (2019). CTCF: a Swiss-army knife for genome organization and transcription regulation. Essays in biochemistry, 63(1), pp.157–165.

Busslinger, G.A., Stocsits, R.R., van der Lelij, P., Axelsson, E., Tedeschi, A., Galjart, N., and Peters, J.-M. (2017). Cohesin is positioned in mammalian genomes by transcription, CTCF and Wapl. Nature, 544, 503–507.

Chen, B., Gilbert, L.A., Cimini, B.A., Schnitzbauer, J., Zhang, W., Li, G.W., Park, J., Blackburn, E.H., Weissman, J.S., Qi, L.S. and Huang, B., 2013. Dynamic imaging of genomic loci in living human cells by an optimized CRISPR/Cas system. Cell, 155(7), pp.1479–1491.

Clow, P. A., Du, M., Jillette, N., Taghbalout, A., Zhu, J. J., & Cheng, A. W. (2022). CRISPR- mediated multiplexed live cell imaging of nonrepetitive genomic loci with one guide RNA per locus. Nature Communications, 13, 1871.

Cong, L., Ran, F.A., Cox, D., Lin, S., Barretto, R., Habib, N., Hsu, P.D., Wu, X., Jiang, W., Marraffini, L.A., et al. (2013). Multiplex Genome Engineering Using CRISPR/Cas Systems. Science, 339, 819–823.

Cuddapah, S., Jothi, R., Schones, D.E., Roh, T.Y., Cui, K. and Zhao, K. (2009). Global analysis of the insulator binding protein CTCF in chromatin barrier regions reveals demarcation of active and repressive domains. Genome research, 19(1), 24–32.

Davidson, I. F., Bauer, B., Goetz, D., Tang, W., Wutz, G., & Peters, J. M. (2019). DNA loop extrusion by human cohesin. Science, 366(6471), 1338–1345.

Davidson, I.F., Barth, R., Zaczek, M., van der Torre, J., Tang, W., Nagasaka, K., Janissen, R., Kerssemakers, J., Wutz, G., Dekker, C. and Peters, J.M., (2023). CTCF is a DNA-tension- dependent barrier to cohesin-mediated loop extrusion. Nature, 1-6.

Davis, C.A., Hitz, B.C., Sloan, C.A., Chan, E.T., Davidson, J.M., Gabdank, I., Hilton, J.A., Jain, K., Baymuradov, U.K., Narayanan, A.K., et al. (2018). The Encyclopedia of DNA elements (ENCODE): data portal update. Nucleic Acids Research, 46, D794–D801.

Dejosez, M., Dall’Agnese, A., Ramamoorthy, M., Platt, J., Yin, X., Hogan, M., Brosh, R., Weintraub, A.S., Hnisz, D., Abraham, B.J. and Young, R.A. (2023). Regulatory architecture of housekeeping genes is driven by promoter assemblies. Cell Reports, 42(5).

Dekker, J., Belmont, A.S., Guttman, M., Leshyk, V.O., Lis, J.T., Lomvardas, S., Mirny, L.A., O’Shea, C.C., Park, P.J., Ren, B., et al. (2017). The 4D nucleome project. Nature, 549, 219– 226.

Deshpande, A.S., Ulahannan, N., Pendleton, M., Dai, X., Ly, L., Behr, J.M., Schwenk, S., Liao, W., Augello, M.A., Tyer, C. and Rughani, P. (2022). Identifying synergistic high-order 3D chromatin conformations from genome-scale nanopore concatemer sequencing. Nature Biotechnology, 40(10), 1488–1499.

Dixon, J.R., Selvaraj, S., Yue, F., Kim, A., Li, Y., Shen, Y., Hu, M., Liu, J.S., and Ren, B. (2012). Topological domains in mammalian genomes identified by analysis of chromatin interactions. Nature, 485, 376–380.

Dotson, G.A., Chen, C., Lindsly, S., Cicalo, A., Dilworth, S., Ryan, C., Jeyarajan, S., Meixner, W., Stansbury, C., Pickard, J. and Beckloff, N. (2022). Deciphering multi-way interactions in the human genome. Nature Communications, 13(1), 5498.

Dowen, J. M., Fan, Z. P., Hnisz, D., Ren, G., Abraham, B. J., Zhang, L. N., … & Young, R. A. (2014). Control of cell identity genes occurs in insulated neighborhoods in mammalian chromosomes. Cell, 159(2), 374–387.

Dukler, N., Gulko, B., Huang, Y.-F., and Siepel, A. (2017). Is a super-enhancer greater than the sum of its parts? Nature Genetics, 49, 2–3.

Durand, N.C., Robinson, J.T., Shamim, M.S., Machol, I., Mesirov, J.P., Lander, E.S., and Aiden, E.L. (2016a). Juicebox Provides a Visualization System for Hi-C Contact Maps with Unlimited Zoom. Cell Systems, 3, 99–101.

Durand, N.C., Shamim, M.S., Machol, I., Rao, S.S.P., Huntley, M.H., Lander, E.S., and Aiden, E.L. (2016b). Juicer Provides a One-Click System for Analyzing Loop-Resolution Hi-C Experiments. Cell Systems, 3, 95–98.

El Khattabi, L., Zhao, H., Kalchschmidt, J., Young, N., Jung, S., Van Blerkom, P., Kieffer-Kwon, P., Kieffer-Kwon, K.-R., Park, S., Wang, X., et al. (2019). A Pliable Mediator Acts as a Functional Rather Than an Architectural Bridge between Promoters and Enhancers. Cell, 178, 1145–1158.e20.

Emerson, D.J., Zhao, P.A., Cook, A.L., Barnett, R.J., Klein, K.N., Saulebekova, D., Ge, C., Zhou, L., Simandi, Z., Minsk, M.K. and Titus, K.R. (2022). Cohesin-mediated loop anchors confine the locations of human replication origins. Nature, 606(7915), 812–819.

Ernst, J., and Kellis, M. (2012). ChromHMM: automating chromatin-state discovery and characterization. Nature Methods, 9, 215–216.

Fang, R., Yu, M., Li, G., Chee, S., Liu, T., Schmitt, A.D., and Ren, B. (2016). Mapping of long- range chromatin interactions by proximity ligation-assisted ChIP-seq. Cell Research, 26, 1345– 1348.

Filippova, G.N., Fagerlie, S., Klenova, E.M., Myers, C., Dehner, Y., Goodwin, G., Neiman, P.E., Collins, S.J. and Lobanenkov, V.V. (1996). An exceptionally conserved transcriptional repressor, CTCF, employs different combinations of zinc fingers to bind diverged promoter sequences of avian and mammalian c-myc oncogenes. Molecular and cellular biology.

Fudenberg, G., Imakaev, M., Lu, C., Goloborodko, A., Abdennur, N., & Mirny, L. A. (2016). Formation of chromosomal domains by loop extrusion. Cell reports, 15(9), 2038–2049.

Fullwood, M.J., Liu, M.H., Pan, Y.F., Liu, J., Xu, H., Mohamed, Y.B., Orlov, Y.L., Velkov, S., Ho, A., Mei, P.H., et al. (2009). An oestrogen-receptor-α-bound human chromatin interactome. Nature, 462, 58–64.

Gabriele, M., Brandão, H. B., Grosse-Holz, S., Jha, A., Dailey, G. M., Cattoglio, C., … & Hansen, A. S. (2022). Dynamics of CTCF-and cohesin-mediated chromatin looping revealed by live-cell imaging. Science, 376(6592), 496–501.

Goel, V.Y., Huseyin, M.K. and Hansen, A.S. (2023). Region Capture Micro-C reveals coalescence of enhancers and promoters into nested microcompartments. Nature Genetics, 1- 9.

Grubert, F., Srivas, R., Spacek, D. V., Kasowski, M., Ruiz-Velasco, M., Sinnott-Armstrong, N., Greenside, P., Narasimha, A., Liu, Q., Geller, B., Sanghi, A., Kulik, M., Sa, S., Rabinovitch, M., Kundaje, A., Dalton, S., Zaugg, J.B. & Snyder, M. (2020). Landscape of cohesin-mediated chromatin loops in the human genome. Nature, 583(7818), 737–743.

Haarhuis, J.H.I., van der Weide, R.H., Blomen, V.A., Yáñez-Cuna, J.O., Amendola, M., van Ruiten, M.S., Krijger, P.H.L., Teunissen, H., Medema, R.H., van Steensel, B., et al. (2017). The Cohesin Release Factor WAPL Restricts Chromatin Loop Extension. Cell, 169, 693–707.e14.

Haeussler, M., Zweig, A.S., Tyner, C., Speir, M.L., Rosenbloom, K.R., Raney, B.J., Lee, C.M., Lee, B.T., Hinrichs, A.S., Gonzalez, J.N., et al. (2019). The UCSC Genome Browser database: 2019 update. Nucleic Acids Research, 47, D853–D858.

Hagberg, A., Swart, P., and S Chult, D. (2008). Exploring network structure, dynamics, and function using networkx. Los Alamos National Laboratory, https://www.osti.gov/servlets/purl/960616.

Hand, R., (1978). Eucaryotic DNA: organization of the genome for replication. Cell, 15(2), 317–325.

Heinz, S., Texari, L., Hayes, M.G.B., Urbanowski, M., Chang, M.W., Givarkes, N., Rialdi, A., White, K.M., Albrecht, R.A., Pache, L., et al. (2018). Transcription Elongation Can Affect Genome 3D Structure. Cell, 174, 1522–1536.e22.

Hnisz, D., Abraham, B.J., Lee, T.I., Lau, A., Saint-André, V., Sigova, A.A., Hoke, H.A., and Young, R.A. (2013). Super-Enhancers in the Control of Cell Identity and Disease. Cell, 155, 934–947.

Hsieh, T.-H.S., Weiner, A., Lajoie, B., Dekker, J., Friedman, N., and Rando, O.J. (2015). Mapping Nucleosome Resolution Chromosome Folding in Yeast by Micro-C. Cell, 162, 108– 119.

Hsieh, T. H. S., Cattoglio, C., Slobodyanyuk, E., Hansen, A. S., Darzacq, X., & Tjian, R. (2022). Enhancer-promoter interactions and transcription are largely maintained upon acute loss of CTCF, cohesin, WAPL, or YY1. Nature Genetics, 54, 1919–1932.

Iborra, F.J., Pombo, A., Jackson, D.A., & Cook, P.R. (1996). Active RNA polymerases are localized within discrete transcription “factories” in human nuclei. Journal of cell science, 109(6), 1427–1436.

Jiang, Y., Huang, J., Lun, K., Li, B., Zheng, H., Li, Y., Zhou, R., Duan, W., Wang, C., Feng, Y. and Yao, H., (2020). Genome-wide analyses of chromatin interactions after the loss of Pol I, Pol II, and Pol III. Genome biology, 21(1), pp.1–28.

Kagey, M.H., Newman, J.J., Bilodeau, S., Zhan, Y., Orlando, D.A., van Berkum, N.L., Ebmeier, C.C., Goossens, J., Rahl, P.B., Levine, S.S., et al. (2010). Mediator and cohesin connect gene expression and chromatin architecture. Nature, 467, 430–435.

Kane, L., Williamson, I., Flyamer, I.M., Kumar, Y., Hill, R.E., Lettice, L.A. and Bickmore, W.A., (2022). Cohesin is required for long-range enhancer action at the Shh locus. Nature structural & molecular biology, 29(9), 891–897.

Kharchenko, P.V., Tolstorukov, M.Y., and Park, P.J. (2008). Design and analysis of ChIP-seq experiments for DNA-binding proteins. Nature Biotechnology, 26, 1351–1359.

Kieffer-Kwon, K.-R., Tang, Z., Mathe, E., Qian, J., Sung, M.-H., Li, G., Resch, W., Baek, S., Pruett, N., Grøntved, L., et al. (2013). Interactome Maps of Mouse Gene Regulatory Domains Reveal Basic Principles of Transcriptional Regulation. Cell, 155, 1507–1520.

Kim, M., Zheng, M., Tian, S.Z., Lee, B., Chuang, J.H., and Ruan, Y. (2019). MIA-Sig: multiplex chromatin interaction analysis by signal processing and statistical algorithms. Genome Biology, 20, 251.

Kim, T. H., Abdullaev, Z. K., Smith, A. D., Ching, K. A., Loukinov, D. I., Green, R. D., … & Ren, B. (2007). Analysis of the vertebrate insulator protein CTCF-binding sites in the human genome. Cell, 128(6), 1231–1245.

Kim, Y., Shi, Z., Zhang, H., Finkelstein, I.J., and Yu, H. (2019). Human cohesin compacts DNA by loop extrusion. Science, 366, 1345–1349.

Kubo, N., Ishii, H., Xiong, X., Bianco, S., Meitinger, F., Hu, R., Hocker, J.D., Conte, M., Gorkin, D., Yu, M. and Li, B. (2021). Promoter-proximal CTCF binding promotes distal enhancer- dependent gene activation. Nature structural & molecular biology, 28(2), 152–161.

Lee, B., Wang, J., Cai, L., Kim, M., Namburi, S., Tjong, H., Feng, Y., Wang, P., Tang, Z., Abbas, A., et al. (2020). ChIA-PIPE: A fully automated pipeline for comprehensive ChIA-PET data analysis and visualization. Science Advances, 6, eaay2078.

Li, L., Lyu, X., Hou, C., Takenaka, N., Nguyen, H. Q., Ong, C. T., Cubenas-Potts, C., Hu, M., Lei, E.P., Bosco, G., Qin, Z.S. & Corces, V. G. (2015). Widespread rearrangement of 3D chromatin organization underlies polycomb-mediated stress-induced silencing. Molecular cell, 58(2), 216–231.

Li, G., Ruan, X., Auerbach, R.K., Sandhu, K.S., Zheng, M., Wang, P., Poh, H.M., Goh, Y., Lim, J., Zhang, J., et al. (2012). Extensive Promoter-Centered Chromatin Interactions Provide a Topological Basis for Transcription Regulation. Cell, 148, 84–98.

Li, Y., Haarhuis, J.H.I., Sedeño Cacciatore, Á., Oldenkamp, R., van Ruiten, M.S., Willems, L., Teunissen, H., Muir, K.W., de Wit, E., Rowland, B.D., et al. (2020). The structural basis for cohesin–CTCF-anchored loops. Nature, 578, 472–476.

Lieberman-Aiden, E., van Berkum, N.L., Williams, L., Imakaev, M., Ragoczy, T., Telling, A., Amit, I., Lajoie, B.R., Sabo, P.J., Dorschner, M.O., et al. (2009). Comprehensive Mapping of Long-Range Interactions Reveals Folding Principles of the Human Genome. Science, 326, 289– 293.

Liu, N. Q., Maresca, M., van den Brand, T., Braccioli, L., Schijns, M. M., Teunissen, H., Bruneau, B.G., Nora, E.P. & de Wit, E. (2021). WAPL maintains a cohesin loading cycle to preserve cell-type-specific distal gene regulation. Nature genetics, 53(1), 100–109.

Liu, Y., Zhangding, Z., Liu, X., Gan, T., Ai, C., Wu, J., Liang, H., Chen, M., Guo, Y., Lu, R. and Jiang, Y., (2024). Fork coupling directs DNA replication elongation and termination. Science, 383(6688), 1215–1222.

Luppino, J. M., Field, A., Nguyen, S. C., Park, D. S., Shah, P. P., Abdill, R. J., Lan, Y., Yunker, R., Jain, R., Adelman, K. & Joyce, E. F. (2022). Co-depletion of NIPBL and WAPL balance cohesin activity to correct gene misexpression. PLoS genetics, 18(11), e1010528.

MacPherson, M. J., & Sadowski, P. D. (2010). The CTCF insulator protein forms an unusual DNA structure. BMC molecular biology, 11(1), 1–17.

Mannini, L., C Lamaze, F., Cucco, F., Amato, C., Quarantotti, V., Rizzo, I. M., … & Musio, A. (2015). Mutant cohesin affects RNA polymerase II regulation in Cornelia de Lange syndrome. Scientific reports, 5(1), 1–11.

Merkenschlager, M., & Nora, E. P. (2016). CTCF and cohesin in genome folding and transcriptional gene regulation. Annual review of genomics and human genetics, 17, 17–43.

Mumbach, M.R., Rubin, A.J., Flynn, R.A., Dai, C., Khavari, P.A., Greenleaf, W.J., and Chang, H.Y. (2016). HiChIP: efficient and sensitive analysis of protein-directed genome architecture. Nature Methods, 13, 919–922.

Nagano, T., Lubling, Y., Várnai, C., Dudley, C., Leung, W., Baran, Y., Mendelson Cohen, N., Wingett, S., Fraser, P. and Tanay, A. (2017). Cell-cycle dynamics of chromosomal organization at single-cell resolution. Nature, 547(7661), 61–67.

Natsume, T., Kiyomitsu, T., Saga, Y., & Kanemaki, M. T. (2016). Rapid protein depletion in human cells by auxin-inducible degron tagging with short homology donors. Cell reports, 15(1), 210–218.

Nora, E.P., Lajoie, B.R., Schulz, E.G., Giorgetti, L., Okamoto, I., Servant, N., Piolot, T., van Berkum, N.L., Meisig, J., Sedat, J. and Gribnau, J., (2012). Spatial partitioning of the regulatory landscape of the X-inactivation centre. Nature, 485(7398), 381–385.

Nora, E. P., Goloborodko, A., Valton, A. L., Gibcus, J. H., Uebersohn, A., Abdennur, N., … & Bruneau, B. G. (2017). Targeted degradation of CTCF decouples local insulation of chromosome domains from genomic compartmentalization. Cell, 169(5), 930–944.

Nuebler, J., Fudenberg, G., Imakaev, M., Abdennur, N. and Mirny, L.A., 2018. Chromatin organization by an interplay of loop extrusion and compartmental segregation. Proceedings of the National Academy of Sciences, 115(29), pp.E6697–E6706.

Papatheodorou, I., Moreno, P., Manning, J., Fuentes, A.M.P., George, N., Fexova, S., Fonseca, N.A., Füllgrabe, A., Green, M., Huang, N. and Huerta, L. (2020). Expression Atlas update: from tissues to single cells. Nucleic acids research, 48(D1), D77–D83.

Parslow, A., Cardona, A., and Bryson-Richardson, R.J. (2014). Sample Drift Correction Following 4D Confocal Time-lapse Imaging. Journal of Visualized Experiments, 86, 51086.

Paulsen, J., Liyakat Ali, T.M., Nekrasov, M., Delbarre, E., Baudement, M.-O., Kurscheid, S., Tremethick, D., and Collas, P. (2019). Long-range interactions between topologically associating domains shape the four-dimensional genome during differentiation. Nature Genetics, 51, 835–843.

Peric-Hupkes, D. and van Steensel, B. (2008). Linking cohesin to gene regulation. Cell, 132(6), 925–928.

Pherson, M., Misulovin, Z., Gause, M. and Dorsett, D., (2019). Cohesin occupancy and composition at enhancers and promoters are linked to DNA replication origin proximity in Drosophila. Genome research, 29(4), 602–612.

Phillips-Cremins, J.E., Sauria, M.E.G., Sanyal, A., Gerasimova, T.I., Lajoie, B.R., Bell, J.S.K., Ong, C.-T., Hookway, T.A., Guo, C., Sun, Y., et al. (2013). Architectural Protein Subclasses Shape 3D Organization of Genomes during Lineage Commitment. Cell, 153, 1281–1295.

Quinlan, A.R., and Hall, I.M. (2010). BEDTools: a flexible suite of utilities for comparing genomic features. Bioinformatics, 26, 841–842.

Quinodoz, S. A., Ollikainen, N., Tabak, B., Palla, A., Schmidt, J. M., Detmar, E., Lai, M.M., Shishkin, A.A., Bhat, P., Takei, Y., Trinh, V., Aznauryan E., Russell, P., Cheng, C., Jovanovic, M., Chow, A., Cai, L., McDonel, P., Garber, M. & Guttman, M. (2018). Higher-order inter- chromosomal hubs shape 3D genome organization in the nucleus. Cell, 174(3), 744–757.

Ramírez, F., Ryan, D.P., Grüning, B., Bhardwaj, V., Kilpert, F., Richter, A.S., Heyne, S., Dündar, F., and Manke, T. (2016). deepTools2: a next generation web server for deep-sequencing data analysis. Nucleic Acids Research, 44, W160–W165.

Rao, S.S.P., Huntley, M.H., Durand, N.C., Stamenova, E.K., Bochkov, I.D., Robinson, J.T., Sanborn, A.L., Machol, I., Omer, A.D., Lander, E.S., et al. (2014). A 3D Map of the Human Genome at Kilobase Resolution Reveals Principles of Chromatin Looping. Cell, 159, 1665– 1680.

Rao, S.S.P., Huang, S.-C., Glenn St Hilaire, B., Engreitz, J.M., Perez, E.M., Kieffer-Kwon, K.-R., Sanborn, A.L., Johnstone, S.E., Bascom, G.D., Bochkov, I.D., et al. (2017). Cohesin Loss Eliminates All Loop Domains. Cell, 171, 305–320.e24.

Sabari, B.R., Dall’Agnese, A., Boija, A., Klein, I.A., Coffey, E.L., Shrinivas, K., Abraham, B.J., Hannett, N.M., Zamudio, A.V., Manteiga, J.C., et al. (2018). Coactivator condensation at super- enhancers links phase separation and gene control. Science, 361, eaar3958.

Sanborn, A. L., Rao, S. S., Huang, S. C., Durand, N. C., Huntley, M. H., Jewett, A. I., … & Aiden, E. L. (2015). Chromatin extrusion explains key features of loop and domain formation in wild-type and engineered genomes. Proceedings of the National Academy of Sciences, 112(47), E6456–E6465.

Schaaf, C. A., Misulovin, Z., Gause, M., Koenig, A., Gohara, D. W., Watson, A., & Dorsett, D. (2013). Cohesin and polycomb proteins functionally interact to control transcription at silenced and active genes. PLoS genetics, 9(6), e1003560.

Schindelin, J., Arganda-Carreras, I., Frise, E., Kaynig, V., Longair, M., Pietzsch, T., Preibisch, S., Rueden, C., Saalfeld, S., Schmid, B., et al. (2012). Fiji: an open-source platform for biological-image analysis. Nature Methods, 9, 676–682.

Schuijers, J., Manteiga, J.C., Weintraub, A.S., Day, D.S., Zamudio, A.V., Hnisz, D., Lee, T.I., and Young, R.A. (2018). Transcriptional Dysregulation of MYC Reveals Common Enhancer- Docking Mechanism. Cell Reports, 23, 349–360.

Shannon, C.E. (1948). A Mathematical Theory of Communication. Bell System Technical Journal, 27, 379–423.

Tang, Z., Luo, O.J., Li, X., Zheng, M., Zhu, J.J., Szalaj, P., Trzaskoma, P., Magalska, A., Wlodarczyk, J., Ruszczycki, B., et al. (2015). CTCF-Mediated Human 3D Genome Architecture Reveals Chromatin Topology for Transcription. Cell, 163, 1611–1627.

Thiecke, M. J., Wutz, G., Muhar, M., Tang, W., Bevan, S., Malysheva, V., … & Spivakov, M. (2020). Cohesin-dependent and-independent mechanisms mediate chromosomal contacts between promoters and enhancers. Cell reports, 32(3), 107929.

Tian, S.Z., Capurso, D., Kim, M., Lee, B., Zheng, M., and Ruan, Y. (2019). ChIA-DropBox: a novel analysis and visualization pipeline for multiplex chromatin interactions. bioRxiv.

Tinevez, J.-Y., Perry, N., Schindelin, J., Hoopes, G.M., Reynolds, G.D., Laplantine, E., Bednarek, S.Y., Shorte, S.L., and Eliceiri, K.W. (2017). TrackMate: An open and extensible platform for single-particle tracking. Methods, 115, 80–90.

Ushiki, A., Zhang, Y., Xiong, C., Zhao, J., Georgakopoulos-Soares, I., Kane, L., … & Ahituv, N. (2021). Deletion of CTCF sites in the SHH locus alters enhancer–promoter interactions and leads to acheiropodia. Nature communications, 12(1), 1–12.

Vangala, P., Murphy, R., Quinodoz, S.A., Gellatly, K., McDonel, P., Guttman, M. and Garber, M., (2020). High-resolution mapping of multiway enhancer-promoter interactions regulating pathogen detection. Molecular cell, 80(2), 359–373.

Vian, L., Pękowska, A., Rao, S.S.P., Kieffer-Kwon, K.-R., Jung, S., Baranello, L., Huang, S.-C., El Khattabi, L., Dose, M., Pruett, N., et al. (2018). The Energetics and Physiological Impact of Cohesin Extrusion. Cell, 173, 1165–1178.e20.

Wang, H., Nakamura, M., Abbott, T. R., Zhao, D., Luo, K., Yu, C., Nguyen, C.M., Lo, A., Daley, T.P., La Russa, M., Liu, Y. & Qi, L. S. (2019). CRISPR-mediated live imaging of genome editing and transcription. Science, 365(6459), 1301–1305.

Wang, P., Feng, Y., Zhu, K., Chai, H., Chang, Y.T., Yang, X., Liu, X., Shen, C., Gega, E., Lee, B. and Kim, M. (2021). In Situ Chromatin Interaction Analysis Using Paired-End Tag Sequencing. Current protocols, 1(8), e174.

Weintraub, A. S., Li, C. H., Zamudio, A. V., Sigova, A. A., Hannett, N. M., Day, D. S., … & Young, R. A. (2017). YY1 is a structural regulator of enhancer-promoter loops. Cell, 171(7), 1573–1588.

Wu, H., Zhang, J., Tan, L., Xie, X.S. (2023). Extruding transcription elongation loops observed in high-resolution single-cell 3D genomes. bioRxiv, 2023-02.

Xu, B., Wang, H., Wright, S., Hyle, J., Zhang, Y., Shao, Y., … & Li, C. (2021). Acute depletion of CTCF rewires genome-wide chromatin accessibility. Genome biology, 22(1), 1–25.

Yang, T., Zhang, F., Yardımcı, G.G., Song, F., Hardison, R.C., Noble, W.S., Yue, F., and Li, Q. (2017). HiCRep: assessing the reproducibility of Hi-C data using a stratum-adjusted correlation coefficient. Genome Research, 27, 1939–1949.

Zhang, H., Emerson, D.J., Gilgenast, T.G., Titus, K.R., Lan, Y., Huang, P., Zhang, D., Wang, H., Keller, C.A., Giardine, B. and Hardison, R.C., (2019). Chromatin structure dynamics during the mitosis-to-G1 phase transition. Nature, 576(7785), 158–162.

Zhang, H.B., Kim, M., Chuang, J.H., and Ruan, Y. (2020). pyBedGraph: a python package for fast operations on 1D genomic signal tracks. Bioinformatics, 36, 3234–3235.

Zheng, M., Tian, S.Z., Capurso, D., Kim, M., Maurya, R., Lee, B., Piecuch, E., Gong, L., Zhu, J.J., Li, Z., et al. (2019). Multiplex chromatin interactions with single-molecule precision. Nature, 566, 558–562.

Zhu, Y., Denholtz, M., Lu, H., & Murre, C. (2021). Calcium signaling instructs NIPBL recruitment at active enhancers and promoters via distinct mechanisms to reconstruct genome compartmentalization. Genes & Development, 35(1-2), 65–81.

Zhu, J. J., & Cheng, A. W. (2022). JACKIE: Fast Enumeration of Genome-Wide Single-and Multicopy CRISPR Target Sites and Their Off-Target Numbers. The CRISPR Journal, 5(4), 618–628.

Zuin, J., Franke, V., van IJcken, W.F., Van Der Sloot, A., Krantz, I.D., van der Reijden, M.I., Nakato, R., Lenhard, B. and Wendt, K.S. (2014). A cohesin-independent role for NIPBL at promoters provides insights in CdLS. PLoS genetics, 10(2), e1004153.

