## Supplementary material for "Multifaceted roles of cohesin in regulating transcriptional loops": Supp_Table_S1

| Cell line | Treatment | Hours | Experiment | Factor | Source | Library ID | Replicate | Combined into | Combined into | # of reads | Orig. Complexes (F>1) | Repetitive regions filtered | Entropy filtered (F=1) | Entropy filtered (F>1) | Enrich. Test FDR 0.1 (F=1) | Enrich. Test FDR 0.1 (F>1) | Enrich. Test FDR 0.2 (F=1) | Enrich. Test FDR 0.2 (F>1) |
|---|---|---|---|---|---|---|---|---|---|---|---|---|---|---|---|---|---|---|
| GM12878 | None | 0 | ChIA-Drop | CTCF | This study | SHG0210 | Rep1-1 | GM12878-CTCF-Rep1 | GM12878-CTCF-pooled | 53,809,034 | 791,544 | 618,965 | 1,041,038 | 150,101 | NA | NA | NA | NA |
| GM12878 | None | 0 | ChIA-Drop | CTCF | This study | SHG0211 | Rep1-2 | GM12878-CTCF-Rep1 | GM12878-CTCF-pooled | 64,003,215 | 964,667 | 690,815 | 1,179,667 | 163,843 | NA | NA | NA | NA |
| GM12878 | None | 0 | ChIA-Drop | CTCF | This study | SHG0212 | Rep1-3 | GM12878-CTCF-Rep1 | GM12878-CTCF-pooled | 87,724,974 | 1,341,844 | 1,027,220 | 1,745,852 | 266,143 | NA | NA | NA | NA |
| GM12878 | None | 0 | ChIA-Drop | CTCF | This study | SHG0213 | Rep1-4 | GM12878-CTCF-Rep1 | GM12878-CTCF-pooled | 76,970,686 | 1,103,947 | 847,447 | 1,400,215 | 232,071 | NA | NA | NA | NA |
| GM12878 | None | 0 | ChIA-Drop | CTCF | This study | SHG8113 | Rep1-5 | GM12878-CTCF-Rep1 | GM12878-CTCF-pooled | 89,647,949 | 2,583,586 | 2,099,266 | 3,729,833 | 535,011 | NA | NA | NA | NA |
| GM12878 | None | 0 | ChIA-Drop | CTCF | This study | SHG8132 | Rep1-6 | GM12878-CTCF-Rep1 | GM12878-CTCF-pooled | 59,679,168 | 3,636,322 | 3,020,748 | 5,471,716 | 824,716 | NA | NA | NA | NA |
| GM12878 | None | 0 | ChIA-Drop | CTCF | This study | | Rep1 | GM12878-CTCF-Rep1 | SUM | 431,835,026 | 10,421,910 | 8,304,461 | 14,568,321 | 2,171,885 | 1,446,321 | 172,209 | 3,829,611 | 524,532 |
| GM12878 | None | 0 | ChIA-Drop | CTCF | This study | SHG0214 | Rep2-1 | GM12878-CTCF-Rep2 | GM12878-CTCF-pooled | 95,786,946 | 1,838,463 | 1,469,135 | 2,644,685 | 331,473 | NA | NA | NA | NA |
| GM12878 | None | 0 | ChIA-Drop | CTCF | This study | SHG0215 | Rep2-2 | GM12878-CTCF-Rep2 | GM12878-CTCF-pooled | 80,764,231 | 1,439,764 | 1,121,141 | 1,978,236 | 256,894 | NA | NA | NA | NA |
| GM12878 | None | 0 | ChIA-Drop | CTCF | This study | SHG0216 | Rep2-3 | GM12878-CTCF-Rep2 | GM12878-CTCF-pooled | 72,844,914 | 1,188,792 | 935,629 | 1,632,757 | 212,748 | NA | NA | NA | NA |
| GM12878 | None | 0 | ChIA-Drop | CTCF | This study | SHG0217 | Rep2-4 | GM12878-CTCF-Rep2 | GM12878-CTCF-pooled | 62,791,060 | 2,018,968 | 1,666,936 | 3,021,846 | 382,237 | NA | NA | NA | NA |
| GM12878 | None | 0 | ChIA-Drop | CTCF | This study | SHG8117 | Rep2-5 | GM12878-CTCF-Rep2 | GM12878-CTCF-pooled | 89,781,816 | 1,486,451 | 1,210,392 | 1,965,824 | 359,772 | NA | NA | NA | NA |
| GM12878 | None | 0 | ChIA-Drop | CTCF | This study | SHG8133 | Rep2-6 | GM12878-CTCF-Rep2 | GM12878-CTCF-pooled | 77,645,469 | 3,962,279 | 3,252,336 | 5,692,687 | 1,029,915 | NA | NA | NA | NA |
| GM12878 | None | 0 | ChIA-Drop | CTCF | This study | | Rep2 | GM12878-CTCF-Rep2 | SUM | 479,614,436 | 11,934,717 | 9,655,569 | 16,936,035 | 2,573,039 | 1,574,468 | 202,948 | 4,372,322 | 636,262 |
| GM12878 | None | 0 | ChIA-Drop | CTCF | This study | SHG8033 | Rep3-1 | GM12878-CTCF-Rep3 | GM12878-CTCF-pooled | 51,695,102 | 733,738 | 543,326 | 875,400 | 142,541 | NA | NA | NA | NA |
| GM12878 | None | 0 | ChIA-Drop | CTCF | This study | SHG8065 | Rep3-2 | GM12878-CTCF-Rep3 | GM12878-CTCF-pooled | 69,463,568 | 704,855 | 438,847 | 723,394 | 105,057 | NA | NA | NA | NA |
| GM12878 | None | 0 | ChIA-Drop | CTCF | This study | SHG8098 | Rep3-3 | GM12878-CTCF-Rep3 | GM12878-CTCF-pooled | 64,024,493 | 715,163 | 560,346 | 927,602 | 134,599 | NA | NA | NA | NA |
| GM12878 | None | 0 | ChIA-Drop | CTCF | This study | SHG8099 | Rep3-4 | GM12878-CTCF-Rep3 | GM12878-CTCF-pooled | 63,853,377 | 572,485 | 432,749 | 710,710 | 103,414 | NA | NA | NA | NA |
| GM12878 | None | 0 | ChIA-Drop | CTCF | This study | SHG0193 | Rep3-5 | GM12878-CTCF-Rep3 | GM12878-CTCF-pooled | 50,095,683 | 520,952 | 412,727 | 731,855 | 70,338 | NA | NA | NA | NA |
| GM12878 | None | 0 | ChIA-Drop | CTCF | This study | SHG8035 | Rep3-6 | GM12878-CTCF-Rep3 | GM12878-CTCF-pooled | 83,797,446 | 2,750,133 | 2,205,832 | 4,488,489 | 244,101 | NA | NA | NA | NA |
| GM12878 | None | 0 | ChIA-Drop | CTCF | This study | | Rep3 | GM12878-CTCF-Rep3 | SUM | 382,929,669 | 5,997,326 | 4,593,827 | 8,457,450 | 800,050 | 1,311,673 | 123,156 | 3,100,214 | 319,744 |
| GM12878 | None | 0 | ChIA-Drop | CTCF | This study | | pooled | GM12878-CTCF-pooled | SUM | 1,294,379,131 | 28,353,953 | 22,553,857 | 39,961,806 | 5,544,974 | 3,860,627 | 438,452 | 10,130,243 | 1,315,778 |
| GM12878 | None | 0 | ChIA-Drop | RAD21 | This study | SHG0180 | Rep1-1 | GM12878-RAD21-Rep1 | GM12878-cohesin-pooled | 86,737,328 | 5,114,489 | 4,396,204 | 9,438,034 | 619,616 | NA | NA | NA | NA |
| GM12878 | None | 0 | ChIA-Drop | RAD21 | This study | SHG0195 | Rep1-2 | GM12878-RAD21-Rep1 | GM12878-cohesin-pooled | 156,888,671 | 8,172,660 | 6,922,948 | 15,658,707 | 1,157,328 | NA | NA | NA | NA |
| GM12878 | None | 0 | ChIA-Drop | RAD21 | This study | SHG0198 | Rep1-3 | GM12878-RAD21-Rep1 | GM12878-cohesin-pooled | 56,143,013 | 3,451,076 | 2,971,641 | 6,186,579 | 386,731 | NA | NA | NA | NA |
| GM12878 | None | 0 | ChIA-Drop | RAD21 | This study | SHG0202 | Rep1-4 | GM12878-RAD21-Rep1 | GM12878-cohesin-pooled | 185,966,786 | 9,377,136 | 7,921,234 | 18,477,807 | 1,300,248 | NA | NA | NA | NA |
| GM12878 | None | 0 | ChIA-Drop | RAD21 | This study | SHG0205 | Rep1-5 | GM12878-RAD21-Rep1 | GM12878-cohesin-pooled | 130,723,974 | 8,069,067 | 7,027,539 | 16,154,497 | 1,035,487 | NA | NA | NA | NA |
| GM12878 | None | 0 | ChIA-Drop | RAD21 | This study | SHG0206 | Rep1-6 | GM12878-RAD21-Rep1 | GM12878-cohesin-pooled | 162,624,186 | 8,792,169 | 7,638,734 | 17,749,761 | 1,185,566 | NA | NA | NA | NA |
| GM12878 | None | 0 | ChIA-Drop | RAD21 | This study | SHG0207 | Rep1-7 | GM12878-RAD21-Rep1 | GM12878-cohesin-pooled | 62,793,728 | 4,378,656 | 3,794,757 | 7,909,865 | 587,478 | NA | NA | NA | NA |
| GM12878 | None | 0 | ChIA-Drop | RAD21 | This study | | Rep1 | GM12878-RAD21-Rep1 | SUM | 841,877,686 | 47,355,253 | 40,673,057 | 91,575,250 | 6,272,454 | 24,040,443 | 2,354,936 | 38,564,652 | 3,814,289 |
| GM12878 | None | 0 | ChIA-Drop | RAD21 | This study | SHG0181 | Rep2-1 | GM12878-RAD21-Rep2 | GM12878-cohesin-pooled | 84,530,899 | 5,855,167 | 5,066,326 | 10,806,732 | 857,395 | NA | NA | NA | NA |
| GM12878 | None | 0 | ChIA-Drop | RAD21 | This study | SHG0196 | Rep2-2 | GM12878-RAD21-Rep2 | GM12878-cohesin-pooled | 156,415,452 | 9,227,264 | 7,966,428 | 17,914,120 | 1,615,428 | NA | NA | NA | NA |
| GM12878 | None | 0 | ChIA-Drop | RAD21 | This study | SHG0199 | Rep2-3 | GM12878-RAD21-Rep2 | GM12878-cohesin-pooled | 102,445,657 | 5,881,573 | 4,992,295 | 11,148,999 | 901,052 | NA | NA | NA | NA |
| GM12878 | None | 0 | ChIA-Drop | RAD21 | This study | SHG0203 | Rep2-4 | GM12878-RAD21-Rep2 | GM12878-cohesin-pooled | 191,895,737 | 11,257,752 | 9,780,145 | 23,216,948 | 2,011,434 | NA | NA | NA | NA |
| GM12878 | None | 0 | ChIA-Drop | RAD21 | This study | SHG0208 | Rep2-5 | GM12878-RAD21-Rep2 | GM12878-cohesin-pooled | 68,451,827 | 4,531,214 | 3,915,771 | 8,216,029 | 601,217 | NA | NA | NA | NA |
| GM12878 | None | 0 | ChIA-Drop | RAD21 | This study | SHG0209 | Rep2-6 | GM12878-RAD21-Rep2 | GM12878-cohesin-pooled | 79,128,174 | 4,808,693 | 4,138,099 | 8,709,449 | 652,205 | NA | NA | NA | NA |
| GM12878 | None | 0 | ChIA-Drop | RAD21 | This study | | Rep2 | GM12878-RAD21-Rep2 | SUM | 682,867,746 | 41,561,663 | 35,859,064 | 80,012,277 | 6,638,731 | 19,156,824 | 2,049,377 | 30,243,718 | 3,641,468 |
| GM12878 | None | 0 | ChIA-Drop | RNAPII | This study | SHG8034 | Rep1-1 | GM12878-RNAPII-Rep1 | GM12878-RNAPII-pooled; | 79,192,522 | 3,149,655 | 2,545,237 | 5,250,602 | 280,345 | NA | NA | NA | NA |
| GM12878 | None | 0 | ChIA-Drop | RNAPII | This study | SHG8100 | Rep1-2 | GM12878-RNAPII-Rep1 | GM12878-RNAPII-pooled; | 65,226,738 | 2,035,305 | 1,681,315 | 3,390,414 | 175,460 | NA | NA | NA | NA |
| GM12878 | None | 0 | ChIA-Drop | RNAPII | This study | SHG8101 | Rep1-3 | GM12878-RNAPII-Rep1 | GM12878-RNAPII-pooled; | 67,282,542 | 1,913,204 | 1,583,783 | 3,175,672 | 165,845 | NA | NA | NA | NA |
| GM12878 | None | 0 | ChIA-Drop | RNAPII | This study | SHG8096 | Rep1-4 | GM12878-RNAPII-Rep1 | GM12878-RNAPII-pooled; | 68,385,904 | 1,475,063 | 1,180,104 | 1,939,470 | 337,391 | NA | NA | NA | NA |
| GM12878 | None | 0 | ChIA-Drop | RNAPII | This study | SHG8097 | Rep1-5 | GM12878-RNAPII-Rep1 | GM12878-RNAPII-pooled; | 64,871,999 | 1,345,206 | 1,052,235 | 1,736,384 | 290,936 | NA | NA | NA | NA |
| GM12878 | None | 0 | ChIA-Drop | RNAPII | This study | | Rep1 | GM12878-RNAPII-Rep1 | SUM | 344,959,705 | 9,918,433 | 8,042,674 | 15,492,542 | 1,249,977 | 498,661 | 49,328 | 1,625,961 | 176,301 |
| GM12878 | None | 0 | ChIA-Drop | RNAPII | This study | SHG8112 | Rep2-1 | GM12878-RNAPII-Rep2 | GM12878-RNAPII-pooled; | 67,273,144 | 837,532 | 631,102 | 784,063 | 300,071 | NA | NA | NA | NA |
| GM12878 | None | 0 | ChIA-Drop | RNAPII | This study | SHG8116 | Rep2-2 | GM12878-RNAPII-Rep2 | GM12878-RNAPII-pooled; | 63,420,334 | 2,040,875 | 1,649,997 | 2,681,333 | 538,143 | NA | NA | NA | NA |
| GM12878 | None | 0 | ChIA-Drop | RNAPII | This study | SHG8130 | Rep2-3 | GM12878-RNAPII-Rep2 | GM12878-RNAPII-pooled; | 73,720,406 | 1,515,204 | 1,115,839 | 1,493,472 | 505,536 | NA | NA | NA | NA |
| GM12878 | None | 0 | ChIA-Drop | RNAPII | This study | SHG8131 | Rep2-4 | GM12878-RNAPII-Rep2 | GM12878-RNAPII-pooled; | 50,726,668 | 3,737,580 | 3,107,110 | 5,080,082 | 1,208,344 | NA | NA | NA | NA |
| GM12878 | None | 0 | ChIA-Drop | RNAPII | This study | SHG0218 | Rep2-5 | GM12878-RNAPII-Rep2 | GM12878-RNAPII-pooled; | 57,701,716 | 1,348,602 | 1,079,930 | 1,973,798 | 201,193 | NA | NA | NA | NA |
| GM12878 | None | 0 | ChIA-Drop | RNAPII | This study | SHG0219 | Rep2-6 | GM12878-RNAPII-Rep2 | GM12878-RNAPII-pooled; | 57,662,440 | 1,402,528 | 1,157,338 | 2,129,293 | 209,548 | NA | NA | NA | NA |
| GM12878 | None | 0 | ChIA-Drop | RNAPII | This study | | Rep2 | GM12878-RNAPII-Rep2 | SUM | 370,504,708 | 10,882,321 | 8,741,316 | 14,142,041 | 2,962,835 | 1,766,059 | 324,424 | 4,009,518 | 802,527 |
| GM12878 | None | 0 | ChIA-Drop | RNAPII | This study | SHG0234 | Rep3-1 | GM12878-RNAPII-Rep3 | GM12878-RNAPII-pooledv | 77,820,387 | 1,783,676 | 1,516,515 | 2,835,101 | 266,791 | NA | NA | NA | NA |
| GM12878 | None | 0 | ChIA-Drop | RNAPII | This study | SHG0235 | Rep3-2 | GM12878-RNAPII-Rep3 | GM12878-RNAPII-pooledv | 87,702,678 | 6,465,918 | 5,715,047 | 12,226,757 | 1,065,348 | NA | NA | NA | NA |
| GM12878 | None | 0 | ChIA-Drop | RNAPII | This study | SHG0238 | Rep3-3 | GM12878-RNAPII-Rep3 | GM12878-RNAPII-pooledv | 78,539,454 | 617,441 | 438,423 | 750,148 | 89,533 | NA | NA | NA | NA |
| GM12878 | None | 0 | ChIA-Drop | RNAPII | This study | SHG0239 | Rep3-4 | GM12878-RNAPII-Rep3 | GM12878-RNAPII-pooledv | 94,222,395 | 2,919,656 | 2,447,198 | 4,779,894 | 425,597 | NA | NA | NA | NA |
| GM12878 | None | 0 | ChIA-Drop | RNAPII | This study | | Rep3 | GM12878-RNAPII-Rep3 | SUM | 338,284,914 | 11,786,691 | 10,117,183 | 20,591,900 | 1,847,269 | 7,280,519 | 911,541 | 11,783,316 | 1,296,009 |
| GM12878 | None | 0 | ChIA-Drop | RNAPII | This study | | pooledv2 | GM12878-RNAPII-pooledv2 | SUM | 1,053,749,327 | 32,587,445 | 26,901,173 | 50,226,483 | 6,060,081 | 9,798,884 | 1,263,151 | 15,607,687 | 2,399,732 |
| GM12878 | None | 0 | ChIA-Drop | SMC1A | This study | SHG0182 | Rep1-1 | GM12878-SMC1A-pooled | GM12878-cohesin-pooled | 88,752,922 | 6,607,743 | 5,793,086 | 12,891,264 | 866,372 | NA | NA | NA | NA |
| GM12878 | None | 0 | ChIA-Drop | SMC1A | This study | SHG0197 | Rep1-2 | GM12878-SMC1A-pooled | GM12878-cohesin-pooled | 149,720,829 | 9,006,846 | 7,879,794 | 18,316,671 | 1,300,389 | NA | NA | NA | NA |
| GM12878 | None | 0 | ChIA-Drop | SMC1A | This study | SHG0200 | Rep1-3 | GM12878-SMC1A-pooled | GM12878-cohesin-pooled | 58,803,883 | 4,233,583 | 3,642,191 | 7,991,181 | 502,120 | NA | NA | NA | NA |
| GM12878 | None | 0 | ChIA-Drop | SMC1A | This study | SHG0204 | Rep1-4 | GM12878-SMC1A-pooled | GM12878-cohesin-pooled | 177,247,438 | 12,688,027 | 11,316,882 | 28,633,048 | 2,177,644 | NA | NA | NA | NA |
| GM12878 | None | 0 | ChIA-Drop | SMC1A | This study | SHG0186 | Rep1-5 | GM12878-SMC1A-pooled | GM12878-cohesin-pooled | 147,848,455 | 7,014,380 | 5,878,466 | 13,114,151 | 962,827 | NA | NA | NA | NA |
| GM12878 | None | 0 | ChIA-Drop | SMC1A | This study | SHG0201 | Rep1-6 | GM12878-SMC1A-pooled | GM12878-cohesin-pooled | 57,751,339 | 3,872,026 | 3,276,503 | 6,829,458 | 496,042 | NA | NA | NA | NA |
| GM12878 | None | 0 | ChIA-Drop | cohesin | This study | | pooled | GM12878-cohesin-pooled | SUM | 2,204,870,298 | 132,339,521 | 114,319,043 | 259,363,300 | 19,216,579 | 54,701,855 | 5,705,174 | 93,673,795 | 10,211,318 |
| GM12878 | None | 0 | ChIA-PET | CTCF | This study | LHG0052H | Rep1 | N/A | N/A | 358,752,218 | | | | | | | | |
| GM12878 | None | 0 | ChIA-PET | SMC1A | This study | LHG0051H | Rep1 | N/A | N/A | 356,152,208 | | | | | | | | |
| GM12878 | None | 0 | ChIA-PET | RAD21 | This study | LHG0104V | Rep1 | N/A | N/A | 459,140,632 | | | | | | | | |
| GM12878 | None | 0 | ChIA-PET | Cohesin | This study | LHG0051H_0104V | pooled | N/A | N/A | 815,292,840 | | | | | | | | |
| GM12878 | None | 0 | ChIA-PET | RNAPII | This study | LHG0035N | Rep1 | N/A | N/A | 132,169,141 | | | | | | | | |
| GM12878 | None | 0 | ChIA-PET | RNAPII | This study | LHG0035V | Rep1 | N/A | N/A | 253,204,012 | | | | | | | | |
| GM12878 | None | 0 | ChIA-PET | RNAPII | This study | LHG0045V | Rep2 | N/A | N/A | 494,494,353 | | | | | | | | |
| GM12878 | None | 0 | ChIA-PET | RNAPII | This study | LHG0035N_0035V_0045V | pooled | N/A | N/A | 879,867,506 | | | | | | | | |
| GM12878 | None | 0 | SPRITE | None | 4D Nucleome | 4DNESI1U7ZW9 | pooled | N/A | N/A | 1,610,209,838 | | | | | | | | |
| GM12878 | None | 0 | Hi-C | None | 4D Nucleome | 4DNFI7J8BQ4P | pooled | N/A | N/A | 826,414,879 | | | | | | | | |
| GM19239 | None | 0 | Hi-C | None | 4D Nucleome | 4DNESVKLYDOH | pooled | N/A | N/A | 750,185,985 | | | | | | | | |
| HCT116 | None | 0 | ChIA-PET | CTCF | This study | LHH0118V | Rep1 | N/A | N/A | 293,527,935 | | | | | | | | |
| HCT116 | None | 0 | ChIA-PET | CTCF | This study | LHH0127V | Rep2 | N/A | N/A | 306,988,032 | | | | | | | | |
| HCT116 | None | 0 | ChIA-PET | CTCF | This study | LHH0118V_0127V | pooled | N/A | N/A | 600,515,967 | | | | | | | | |
| HCT116 | None | 0 | ChIA-PET | RNAPII | This study | LHH0098V | Rep1 | N/A | N/A | 517,207,217 | | | | | | | | |
| HCT116 | None | 0 | ChIA-PET | RNAPII | This study | LHH0128V | Rep2 | N/A | N/A | 316,087,217 | | | | | | | | |
| HCT116 | None | 0 | ChIA-PET | RNAPII | This study | LHH0098V_0128V | pooled | N/A | N/A | 833,294,434 | | | | | | | | |
| HCT116 | None | 0 | ChIA-PET | RAD21 | This study | LHH0139 | Rep1 | N/A | N/A | 48,523,665 | | | | | | | | |
| HCT116 | None | 0 | ChIA-PET | RAD21 | This study | LHH0139V | Rep1 | N/A | N/A | 90,562,053 | | | | | | | | |
| HCT116 | None | 0 | ChIA-PET | RAD21 | This study | LHH0140 | Rep2 | N/A | N/A | 42,973,485 | | | | | | | | |
| HCT116 | None | 0 | ChIA-PET | RAD21 | This study | LHH0140V | Rep2 | N/A | N/A | 110,921,989 | | | | | | | | |
| HCT116 | None | 0 | ChIA-PET | RAD21 | This study | LHH0139_0139V_0140_01 | pooled | N/A | N/A | 292,981,192 | | | | | | | | |
| HCT116 | RAD-mAC-auxin | 0 | Hi-C | None | This study | LHH0146V | Rep1 | N/A | N/A | 360,397,636 | | | | | | | | |
| HCT116 | RAD-mAC-auxin | 6 | Hi-C | None | This study | LHH0147V | Rep1 | N/A | N/A | 372,697,334 | | | | | | | | |
| HCT116 | RAD-mAC-auxin | 0 | Hi-C | None | 4D Nucleome | 4DNFIP71EWXC | Rep1 | N/A | N/A | 3,475,834,764 | | | | | | | | |
| HCT116 | RAD-mAC-auxin | 6 | Hi-C | None | 4D Nucleome | 4DNFILIM6FDL | Rep1 | N/A | N/A | 3,365,153,349 | | | | | | | | |
| HCT116 | RAD-mAC-auxin | 0 | ChIA-PET | CTCF | This study | LHH0157 | Rep1 | N/A | N/A | 366,508,457 | | | | | | | | |
| HCT116 | RAD-mAC-auxin | 0 | ChIA-PET | CTCF | This study | LHH0172 | Rep2 | N/A | N/A | 569,205,831 | | | | | | | | |
| HCT116 | RAD-mAC-auxin | 0 | ChIA-PET | CTCF | This study | LHH0157_0172 | pooled | N/A | N/A | 935,714,288 | | | | | | | | |
| HCT116 | RAD-mAC-auxin | 6 | ChIA-PET | CTCF | This study | LHH0158 | Rep1 | N/A | N/A | 401,699,562 | | | | | | | | |
| HCT116 | RAD-mAC-auxin | 6 | ChIA-PET | CTCF | This study | LHH0173 | Rep2 | N/A | N/A | 460,564,094 | | | | | | | | |
| HCT116 | RAD-mAC-auxin | 6 | ChIA-PET | CTCF | This study | LHH0158_0173 | pooled | N/A | N/A | 862,263,656 | | | | | | | | |
| HCT116 | RAD-mAC-auxin | 0 | ChIA-PET | RNAPII | This study | LHH0170 | Rep1 | N/A | N/A | 557,061,917 | | | | | | | | |
| HCT116 | RAD-mAC-auxin | 0 | ChIA-PET | RNAPII | This study | LHH0159 | Rep2 | N/A | N/A | 343,180,214 | | | | | | | | |
| HCT116 | RAD-mAC-auxin | 0 | ChIA-PET | RNAPII | This study | LHH0159V | Rep2 | N/A | N/A | 483,901,014 | | | | | | | | |
| HCT116 | RAD-mAC-auxin | 0 | ChIA-PET | RNAPII | This study | LHH0170_0159_0159V | pooled | N/A | N/A | 1,384,143,145 | | | | | | | | |
| HCT116 | RAD-mAC-auxin | 6 | ChIA-PET | RNAPII | This study | LHH0171 | Rep1 | N/A | N/A | 565,251,115 | | | | | | | | |
| HCT116 | RAD-mAC-auxin | 6 | ChIA-PET | RNAPII | This study | LHH0160 | Rep2 | N/A | N/A | 363,275,787 | | | | | | | | |
| HCT116 | RAD-mAC-auxin | 6 | ChIA-PET | RNAPII | This study | LHH0160V | Rep2 | N/A | N/A | 533,711,372 | | | | | | | | |
| HCT116 | RAD-mAC-auxin | 6 | ChIA-PET | RNAPII | This study | LHH0171_0160_0160V | pooled | N/A | N/A | 1,462,238,274 | | | | | | | | |
| H1 | None | 0 | ChIA-PET | RNAPII | ENCODE | LHES0007H; ENCSR782EKZ | Rep1 | N/A | N/A | 302,061,532 | | | | | | | | |
| K562 | None | 0 | ChIA-PET | RNAPII | ENCODE | LHK0009H_0011V_0012V; | pooled | N/A | N/A | | | | | | | | | |
| HepG2 | None | 0 | ChIA-PET | RNAPII | ENCODE | LHH0123_0123V_0124_01 | pooled | N/A | N/A | | | | | | | | | |
| MCF7 | None | 0 | ChIA-PET | RNAPII | ENCODE | LHM0011_0011H_0014_00 | pooled | N/A | N/A | | | | | | | | | |

| Cell line | Treatment | Hours | Experiment | Factor | Source | Library ID | # of reads |
|---|---|---|---|---|---|---|---|
| GM12878 | None | 0 | ChIP-seq | NIPBL | This study | CHG0030 | 61,848,343 |
| GM12878 | None | 0 | ChIP-seq | CTCF | This study | CHG0031 | 77,693,935 |
| GM12878 | None | 0 | ChIP-seq | WAPL | This study | CHG0032 | 47,004,834 |
| GM12878 | None | 0 | ChIP-seq | Input | This study | CHG0033 | 57,104,734 |
| GM12878 | None | 0 | ChIP-seq | RAD21 | This study | CHG0034 | 56,474,262 |
| GM12878 | None | 0 | ChIP-seq | H3K27ac | ENCODE consortium | ENCFF340JIF | 36,793,809 |
| GM12878 | None | 0 | ChIP-seq | H3K4me1 | ENCODE consortium | ENCFF831ZHL | 63,144,095 |
| GM12878 | None | 0 | chromHMM | N/A | Broad inst. UCSC | N/A | N/A |
| GM12878 | None | 0 | Super-enhancer | N/A | Hsniz et al., 2013 | N/A | N/A |
| GM12878 | None | 0 | CTCF motif | N/A | Tang et al., 2015 | N/A | N/A |
| HCT116 | RAD-mAC-auxin | 0 | ChIP-seq | RAD21 | This study | CHH0056 | 78,476,992 |
| HCT116 | RAD-mAC-auxin | 6 | ChIP-seq | RAD21 | This study | CHH0057 | 90,082,366 |
| HCT116 | RAD-mAC-auxin | 9 | ChIP-seq | RAD21 | This study | CHH0058 | 76,173,734 |
| HCT116 | RAD-mAC-auxin | 12 | ChIP-seq | RAD21 | This study | CHH0059 | 99,661,912 |
| HCT116 | RAD-mAC-auxin | 0 | ChIP-seq | CTCF | This study | CHH0060 | 86,801,317 |
| HCT116 | RAD-mAC-auxin | 6 | ChIP-seq | CTCF | This study | CHH0061 | 93,779,737 |
| HCT116 | RAD-mAC-auxin | 9 | ChIP-seq | CTCF | This study | CHH0062 | 68,463,927 |
| HCT116 | RAD-mAC-auxin | 12 | ChIP-seq | CTCF | This study | CHH0063 | 96,902,097 |
| HCT116 | RAD-mAC-auxin | 0 | ChIP-seq | RNAPII | This study | CHH0064 | 90,338,263 |
| HCT116 | RAD-mAC-auxin | 6 | ChIP-seq | RNAPII | This study | CHH0065 | 76,276,878 |
| HCT116 | RAD-mAC-auxin | 9 | ChIP-seq | RNAPII | This study | CHH0066 | 82,799,502 |
| HCT116 | RAD-mAC-auxin | 12 | ChIP-seq | RNAPII | This study | CHH0067 | 95,021,331 |
| HCT116 | RAD-mAC-auxin | 0 | Repli-seq | N/A | 4D Nucleome | 4DNESNGZM5FG | N/A |
| HCT116 | RAD-mAC-auxin | 6 | Repli-seq | N/A | 4D Nucleome | 4DNES92AU9JR | N/A |
| HCT116 | None | 0 | chromHMM | N/A | ENCODE consortium | ENCFF513PJK | N/A |
| HCT116 | None | 0 | Super-enhancer | N/A | Hsniz et al., 2013 | N/A | N/A |
| MCF7 | None | 0 | chromHMM | N/A | ENCODE consortium | ENCFF506GEX | N/A |
| K562 | None | 0 | chromHMM | N/A | Broad inst. UCSC | N/A | N/A |
| HepG2 | None | 0 | chromHMM | N/A | Broad inst. UCSC | N/A | N/A |
| H1 | None | 0 | chromHMM | N/A | Broad inst. UCSC | N/A | N/A |

| Cell line | Treatment | Hours | Experiment | Source | Library ID | # of reads |
|---|---|---|---|---|---|---|
| GM12878 | None | 0 | RNA-seq | ENCODE consortium | ENCLB555AQG | 117,876,320 |
| HCT116 | RAD-mAC-auxin | 0 | RNA-seq | This study | RHH0001 | 41,471,532 |
| HCT116 | RAD-mAC-auxin | 0 | RNA-seq | This study | RHH0005 | 69,861,080 |
| HCT116 | RAD-mAC-auxin | 6 | RNA-seq | This study | RHH0002 | 54,085,738 |
| HCT116 | RAD-mAC-auxin | 6 | RNA-seq | This study | RHH0006 | 74,474,501 |
| HCT116 | RAD-mAC-auxin | 9 | RNA-seq | This study | RHH0003 | 56,290,599 |
| HCT116 | RAD-mAC-auxin | 9 | RNA-seq | This study | RHH0007 | 57,358,596 |
| HCT116 | RAD-mAC-auxin | 12 | RNA-seq | This study | RHH0004 | 52,650,577 |
| HCT116 | RAD-mAC-auxin | 12 | RNA-seq | This study | RHH0008 | 66,396,860 |
| H1 | None | 0 | RNA-seq | ENCODE consortium | ENCLB555AMA | 125,395,196 |
| K562 | None | 0 | RNA-seq | ENCODE consortium | ENCLB555AKN | 113,588,758 |
| HepG2 | None | 0 | RNA-seq | ENCODE consortium | ENCLB555AQD | 124,172,649 |
| MCF7 | None | 0 | RNA-seq | ENCODE consortium | ENCLB555AQN | 128,178,110 |

| Cell line | Treatment | Hours | Experiment | Source | Probe loci | # of nuclei | # of pairs | # of distances | Supp. Videos |
|---|---|---|---|---|---|---|---|---|---|
| HCT116/RAD21-mAC/dCas9 | None | 0 | Casilio | This study | *BCL6* | 21 | 27 | 1107 | S1 |
| HCT116/RAD21-mAC/dCas9 | Auxin | 24 | Casilio | This study | *BCL6* | 26 | 38 | 1484 | S2 |
| HCT116/RAD21-mAC/dCas9 | None | 0 | Casilio | This study | *SOX9* | 17 | 31 | 1166 | S3 |
| HCT116/RAD21-mAC/dCas9 | Auxin | 24 | Casilio | This study | *SOX9* | 15 | 23 | 943 | S4 |

| | | | | | | | | | | | |
|---|---|---|---|---|---|---|---|---|---|---|---|
| Cell line | GM12878 | GM12878 | GM12878 | GM12878 | GM12878 | GM12878 | GM12878 | GM12878 | GM12878 | GM12878 | GM12878 |
| IP-factor | CTCF | CTCF | CTCF | CTCF | CTCF | CTCF | CTCF | CTCF | CTCF | CTCF | CTCF |
| lib | SHG0193 | SHG0210 | SHG0211 | SHG0212 | SHG0213 | SHG0214 | SHG0215 | SHG0216 | SHG0217 | SHG8033 | SHG8035 |
| Ref. Genome | hg38 | hg38 | hg38 | hg38 | hg38 | hg38 | hg38 | hg38 | hg38 | hg38 | hg38 |
| pcr_duplication | 0.02846304 | 0.02232472 | 0.02911794 | 0.03511322 | 0.02911747 | 0.03527989 | 0.03000722 | 0.02844855 | 0.02422908 | 0.02405731 | 0.0474847 |
| Total_PET_(Paired_end_reads) | 50,095,683 | 53,809,034 | 64,003,215 | 87,724,974 | 76,970,686 | 95,786,946 | 80,764,231 | 72,844,914 | 62,791,060 | 51,695,102 | 83,797,446 |
| PET_with_BC_(BarCode) | 39,872,030 | 42,176,374 | 50,948,959 | 69,070,748 | 60,371,463 | 78,497,190 | 65,747,475 | 57,541,619 | 55,393,822 | 36,833,007 | 71,371,861 |
| Uniq_Mappable_reads (R1) | 37,204,747 | 39,818,903 | 48,053,958 | 64,921,051 | 56,875,230 | 73,763,730 | 61,661,275 | 53,795,101 | 52,562,678 | 35,146,861 | 69,353,579 |
| Reads_mapq>=30 | 27,457,413 | 29,156,604 | 34,244,792 | 47,531,107 | 41,776,872 | 55,753,439 | 45,905,695 | 39,568,073 | 41,989,305 | 24,343,565 | 53,326,491 |
| Reads_q>=30_&_len>=50bp | 25,969,870 | 27,773,006 | 32,681,246 | 45,168,473 | 39,719,788 | 53,002,068 | 43,632,326 | 37,493,110 | 40,221,986 | 23,192,800 | 51,497,910 |
| Reads_q>=30_& _len>=50bp_&_3'ext500 | 25,547,959 | 27,378,204 | 32,165,303 | 44,540,686 | 39,178,246 | 52,265,577 | 43,001,284 | 36,975,642 | 39,888,849 | 22,970,144 | 50,900,386 |
| Reads_q>=30_&_len>=50bp_& _3'ext500_&_MAJOR_CHROM | 25,547,959 | 27,378,204 | 32,165,303 | 44,540,686 | 39,178,246 | 52,265,577 | 43,001,284 | 36,975,642 | 39,888,849 | 22,970,144 | 50,900,386 |
| BC_count_of_PET-w/-BC_(BarCode) | 1,772,481 | 1,762,747 | 1,865,890 | 2,016,988 | 1,917,566 | 2,089,977 | 1,959,526 | 1,937,727 | 1,829,715 | 2,102,213 | 2,386,536 |
| BC_of_Uniq_Mappable_reads_(R1) | 1,747,622 | 1,741,282 | 1,839,903 | 1,990,117 | 1,892,314 | 2,060,361 | 1,930,980 | 1,910,217 | 1,807,211 | 2,083,999 | 2,369,710 |
| BC_of_Reads_q>=30 & len>=50bp & 3'ext500 & MAJOR_CHROM | 1,596,356 | 1,594,196 | 1,679,722 | 1,817,450 | 1,728,402 | 1,903,802 | 1,778,739 | 1,759,094 | 1,688,512 | 1,940,561 | 2,243,710 |
| GEM_TOTAL | 1,596,356 | 1,594,196 | 1,679,722 | 1,817,450 | 1,728,402 | 1,903,802 | 1,778,739 | 1,759,094 | 1,688,512 | 1,940,561 | 2,243,710 |
| GEM_singleFrag | 444,317 | 440,807 | 519,437 | 527,685 | 491,659 | 544,079 | 516,730 | 502,183 | 405,159 | 620,589 | 564,290 |
| GEM_multiFrags | 1,152,039 | 1,153,389 | 1,160,285 | 1,289,765 | 1,236,743 | 1,359,723 | 1,262,009 | 1,256,911 | 1,283,353 | 1,319,972 | 1,679,420 |
| GEM_of_intra_chrom | 18,514 | 14,662 | 13,683 | 12,180 | 13,282 | 9,194 | 9,651 | 11,347 | 5,367 | 25,332 | 10,573 |
| GEM_of_inter_chrom | 1,133,525 | 1,138,727 | 1,146,602 | 1,277,585 | 1,223,461 | 1,350,529 | 1,252,358 | 1,245,564 | 1,277,986 | 1,294,640 | 1,668,847 |
| GEM_of_inter_chrom_divby_ chrom_has_multiFrags | 502,438 | 776,882 | 950,984 | 1,329,664 | 1,090,665 | 1,829,269 | 1,430,113 | 1,177,445 | 2,013,601 | 708,406 | 2,739,560 |
| GEM_of_inter_chrom_divby_ chrom_has_singleFrag | 3,811,330 | 4,209,325 | 4,486,721 | 5,343,336 | 4,833,333 | 6,257,545 | 5,482,278 | 5,136,950 | 6,347,235 | 4,321,196 | 8,633,129 |
| GEM_of_TOTAL_SA_ [SingleFrag+IntraFrags] | 4,776,599 | 5,441,676 | 5,970,825 | 7,212,865 | 6,428,939 | 8,640,087 | 7,438,772 | 6,827,925 | 8,771,362 | 5,675,523 | 11,947,552 |
| GEM_with_Frags=1 | 4,255,647 | 4,650,132 | 5,006,158 | 5,871,021 | 5,324,992 | 6,801,624 | 5,999,008 | 5,639,133 | 6,752,394 | 4,941,785 | 9,197,419 |
| GEM_with_Frags=2 | 451,522 | 646,944 | 766,429 | 1,033,107 | 863,940 | 1,385,716 | 1,107,676 | 938,347 | 1,506,691 | 598,865 | 2,090,724 |
| GEM_with_Frags=3 | 55,895 | 109,882 | 145,283 | 219,748 | 172,215 | 322,728 | 239,076 | 184,496 | 363,299 | 97,114 | 488,682 |
| GEM_with_Frags=4 | 9,078 | 23,185 | 34,360 | 56,084 | 42,740 | 84,639 | 60,090 | 43,415 | 96,960 | 22,565 | 119,319 |
| GEM_with_Frags=5 | 2,222 | 6,630 | 10,230 | 18,052 | 13,413 | 26,072 | 18,461 | 12,864 | 30,163 | 7,850 | 32,645 |
| GEM_with_Frags=6 | 851 | 2,286 | 3,805 | 6,770 | 5,138 | 9,502 | 6,721 | 4,680 | 10,949 | 3,494 | 10,364 |
| GEM_with_Frags=7 | 472 | 1,077 | 1,737 | 3,164 | 2,482 | 4,110 | 3,010 | 2,034 | 4,748 | 1,581 | 3,784 |
| GEM_with_Frags=8 | 259 | 539 | 939 | 1,682 | 1,388 | 2,059 | 1,615 | 1,038 | 2,395 | 849 | 1,747 |
| GEM_with_Frags=9 | 193 | 333 | 534 | 1,032 | 766 | 1,162 | 908 | 618 | 1,309 | 541 | 985 |
| GEM_with_Frags=10 | 128 | 182 | 390 | 639 | 517 | 698 | 589 | 387 | 810 | 284 | 509 |
| GEM_with_Frags=11 | 67 | 126 | 265 | 415 | 346 | 470 | 405 | 216 | 486 | 184 | 376 |
| GEM_with_Frags=12 | 57 | 108 | 157 | 265 | 243 | 300 | 308 | 194 | 353 | 107 | 229 |
| GEM_with_Frags=13 | 52 | 70 | 128 | 196 | 187 | 240 | 187 | 119 | 212 | 96 | 177 |
| GEM_with_Frags=14 | 39 | 47 | 91 | 168 | 129 | 155 | 162 | 88 | 157 | 46 | 120 |
| GEM_with_Frags=15~19 | 79 | 90 | 225 | 358 | 291 | 412 | 364 | 197 | 344 | 123 | 243 |
| GEM_with_Frags=20~29 | 25 | 42 | 88 | 150 | 137 | 164 | 163 | 84 | 88 | 33 | 164 |
| GEM_with_Frags=30~39 | 11 | 2 | 5 | 14 | 12 | 28 | 28 | 13 | 2 | 5 | 45 |
| GEM_with_Frags=40~49 | 1 | 1 | 0 | 0 | 2 | 5 | 1 | 2 | 2 | 1 | 12 |
| GEM_with_Frags=50~99 | 1 | 0 | 1 | 0 | 1 | 3 | 0 | 0 | 0 | 0 | 8 |
| GEM_with_Frags=100~199 | 0 | 0 | 0 | 0 | 0 | 0 | 0 | 0 | 0 | 0 | 0 |
| GEM_with_Frags=200~499 | 0 | 0 | 0 | 0 | 0 | 0 | 0 | 0 | 0 | 0 | 0 |
| GEM_with_Frags=500~999 | 0 | 0 | 0 | 0 | 0 | 0 | 0 | 0 | 0 | 0 | 0 |
| GEM_with_Frags>=1000 | 0 | 0 | 0 | 0 | 0 | 0 | 0 | 0 | 0 | 0 | 0 |
| GEM_with_Frags>=2 | 520,952 | 791,544 | 964,667 | 1,341,844 | 1,103,947 | 1,838,463 | 1,439,764 | 1,188,792 | 2,018,968 | 733,738 | 2,750,133 |

| GM12878 | GM12878 | GM12878 | GM12878 | GM12878 | GM12878 | GM12878 | GM12878 | GM12878 | GM12878 | GM12878 | GM12878 | GM12878 | GM12878 |
|---|---|---|---|---|---|---|---|---|---|---|---|---|---|
| CTCF | CTCF | CTCF | CTCF | CTCF | CTCF | CTCF | RAD21 | RAD21 | RAD21 | RAD21 | RAD21 | RAD21 | RAD21 |
| SHG8065 | SHG8098 | SHG8099 | SHG8113 | SHG8117 | SHG8132 | SHG8133 | SHG0180 | SHG0181 | SHG0195 | SHG0196 | SHG0198 | SHG0199 | SHG0202 |
| hg38 | hg38 | hg38 | hg38 | hg38 | hg38 | hg38 | hg38 | hg38 | hg38 | hg38 | hg38 | hg38 | hg38 |
| 0.02039344 | 0.01942816 | 0.01845405 | 0.02347501 | 0.02144112 | 0.0490809 | 0.06331885 | 0.04587731 | 0.0423211 | 0.07546764 | 0.06999088 | 0.02459719 | 0.04832803 | 0.06393175 |
| 69,463,568 | 64,024,493 | 63,853,377 | 89,647,949 | 89,781,816 | 59,679,168 | 77,645,469 | 86,737,328 | 84,530,899 | 156,888,671 | 156,415,452 | 56,143,013 | 102,445,657 | 185,966,786 |
| 42,913,924 | 41,494,304 | 38,705,371 | 68,003,364 | 59,499,825 | 45,518,623 | 56,416,833 | 81,245,068 | 79,961,732 | 145,105,015 | 146,085,990 | 53,219,658 | 96,778,952 | 174,291,034 |
| 39,628,989 | 39,246,156 | 36,291,400 | 65,265,888 | 56,746,979 | 44,097,839 | 54,504,978 | 77,257,629 | 76,226,127 | 137,567,545 | 139,079,600 | 51,039,139 | 92,936,861 | 166,767,800 |
| 23,129,640 | 26,565,428 | 23,160,539 | 49,161,136 | 40,069,067 | 34,851,418 | 42,275,084 | 61,759,016 | 61,074,010 | 107,831,916 | 110,441,419 | 41,192,989 | 72,508,767 | 136,516,711 |
| 21,664,710 | 25,101,266 | 21,702,443 | 47,035,308 | 37,995,117 | 34,018,551 | 41,136,175 | 59,549,310 | 58,978,852 | 103,697,647 | 106,442,199 | 39,859,302 | 70,336,480 | 131,809,441 |
| 21,415,272 | 24,880,328 | 21,492,394 | 46,471,646 | 37,529,466 | 33,664,036 | 40,720,911 | 59,256,727 | 58,715,139 | 103,054,513 | 105,909,846 | 39,694,561 | 69,945,102 | 131,148,769 |
| 21,415,272 | 24,880,328 | 21,492,394 | 46,471,646 | 37,529,466 | 33,664,036 | 40,720,911 | 59,256,727 | 58,715,139 | 103,054,513 | 105,909,846 | 39,694,561 | 69,945,102 | 131,148,769 |
| 1,870,026 | 1,908,267 | 1,816,956 | 2,153,502 | 2,180,138 | 1,900,443 | 1,907,343 | 2,038,467 | 2,037,963 | 2,724,084 | 2,683,085 | 1,890,494 | 2,210,317 | 2,921,118 |
| 1,846,019 | 1,892,738 | 1,799,767 | 2,137,020 | 2,165,412 | 1,894,014 | 1,899,338 | 2,017,770 | 2,017,626 | 2,701,969 | 2,661,243 | 1,875,073 | 2,183,052 | 2,889,612 |
| 1,650,217 | 1,753,372 | 1,629,261 | 2,007,376 | 2,023,825 | 1,838,423 | 1,832,873 | 1,902,013 | 1,905,534 | 2,562,403 | 2,521,212 | 1,781,341 | 2,009,871 | 2,715,025 |
| 1,650,217 | 1,753,372 | 1,629,261 | 2,007,376 | 2,023,825 | 1,838,423 | 1,832,873 | 1,902,013 | 1,905,534 | 2,562,403 | 2,521,212 | 1,781,341 | 2,009,871 | 2,715,025 |
| 484,442 | 468,648 | 441,258 | 411,650 | 453,143 | 200,641 | 204,899 | 357,600 | 352,028 | 698,983 | 680,411 | 330,641 | 722,801 | 829,390 |
| 1,165,775 | 1,284,724 | 1,188,003 | 1,595,726 | 1,570,682 | 1,637,782 | 1,627,974 | 1,544,413 | 1,553,506 | 1,863,420 | 1,840,801 | 1,450,700 | 1,287,070 | 1,885,635 |
| 22,281 | 20,868 | 23,235 | 7,420 | 16,759 | 4,367 | 3,559 | 3,548 | 3,186 | 15,075 | 13,452 | 2,993 | 9,979 | 15,848 |
| 1,143,494 | 1,263,856 | 1,164,768 | 1,588,306 | 1,553,923 | 1,633,415 | 1,624,415 | 1,540,865 | 1,550,320 | 1,848,345 | 1,827,349 | 1,447,707 | 1,277,091 | 1,869,787 |
| 682,574 | 694,295 | 549,250 | 2,576,166 | 1,469,692 | 3,631,955 | 3,958,720 | 5,110,941 | 5,851,981 | 8,157,585 | 9,213,812 | 3,448,083 | 5,871,594 | 9,361,288 |
| 3,814,982 | 4,327,109 | 3,787,752 | 7,730,734 | 6,187,010 | 8,673,167 | 8,711,157 | 9,732,819 | 10,028,458 | 11,013,824 | 11,094,276 | 8,526,621 | 7,757,077 | 11,157,778 |
| 5,004,279 | 5,510,920 | 4,801,495 | 10,725,970 | 8,126,604 | 12,510,130 | 12,878,335 | 15,204,908 | 16,235,653 | 19,885,467 | 21,001,951 | 12,308,338 | 14,361,451 | 21,364,304 |
| 4,299,424 | 4,795,757 | 4,229,010 | 8,142,384 | 6,640,153 | 8,873,808 | 8,916,056 | 10,090,419 | 10,380,486 | 11,712,807 | 11,774,687 | 8,857,262 | 8,479,878 | 11,987,168 |
| 539,816 | 596,225 | 480,188 | 1,877,831 | 1,148,503 | 2,488,683 | 2,624,917 | 3,439,261 | 3,797,704 | 4,777,764 | 5,170,163 | 2,501,133 | 3,444,020 | 5,245,974 |
| 100,830 | 90,974 | 69,233 | 480,013 | 239,804 | 738,695 | 827,338 | 1,120,698 | 1,325,357 | 1,988,666 | 2,281,151 | 681,673 | 1,441,837 | 2,338,256 |
| 30,937 | 18,531 | 14,548 | 139,186 | 61,630 | 239,827 | 286,280 | 362,041 | 459,430 | 811,731 | 981,888 | 187,276 | 582,009 | 1,005,485 |
| 13,881 | 5,267 | 4,484 | 47,287 | 19,841 | 88,648 | 112,004 | 121,582 | 164,105 | 333,657 | 425,887 | 54,273 | 235,059 | 431,837 |
| 7,539 | 2,041 | 1,904 | 18,838 | 7,863 | 37,086 | 48,652 | 42,413 | 62,006 | 140,874 | 189,997 | 16,858 | 97,222 | 190,166 |
| 4,124 | 941 | 831 | 8,660 | 3,579 | 17,296 | 24,007 | 15,851 | 24,355 | 61,690 | 87,118 | 5,472 | 41,603 | 85,052 |
| 2,467 | 440 | 500 | 4,493 | 2,005 | 9,132 | 12,963 | 6,343 | 10,490 | 28,567 | 41,800 | 2,083 | 18,879 | 39,759 |
| 1,713 | 288 | 276 | 2,588 | 1,101 | 5,388 | 7,730 | 2,631 | 4,844 | 13,266 | 20,665 | 863 | 8,945 | 18,890 |
| 1,118 | 140 | 149 | 1,556 | 650 | 3,365 | 4,907 | 1,218 | 2,351 | 6,562 | 10,742 | 460 | 4,441 | 9,355 |
| 729 | 101 | 96 | 928 | 404 | 2,273 | 3,302 | 703 | 1,297 | 3,351 | 5,921 | 245 | 2,415 | 4,659 |
| 482 | 69 | 70 | 637 | 253 | 1,505 | 2,339 | 419 | 800 | 1,776 | 3,335 | 163 | 1,306 | 2,570 |
| 321 | 34 | 50 | 434 | 185 | 1,032 | 1,653 | 247 | 490 | 1,184 | 2,110 | 112 | 829 | 1,418 |
| 244 | 19 | 23 | 300 | 158 | 794 | 1,228 | 188 | 362 | 693 | 1,411 | 60 | 601 | 822 |
| 480 | 63 | 84 | 606 | 318 | 1,805 | 3,067 | 484 | 887 | 1,503 | 2,770 | 242 | 1,295 | 1,704 |
| 144 | 27 | 39 | 184 | 130 | 666 | 1,407 | 260 | 465 | 886 | 1,548 | 125 | 775 | 816 |
| 17 | 3 | 5 | 38 | 21 | 100 | 294 | 83 | 117 | 280 | 417 | 27 | 201 | 208 |
| 8 | 0 | 3 | 5 | 5 | 18 | 86 | 30 | 60 | 105 | 148 | 6 | 68 | 87 |
| 5 | 0 | 2 | 2 | 1 | 9 | 86 | 36 | 47 | 101 | 170 | 5 | 64 | 70 |
| 0 | 0 | 0 | 0 | 0 | 0 | 15 | 1 | 0 | 4 | 22 | 0 | 3 | 8 |
| 0 | 0 | 0 | 0 | 0 | 0 | 4 | 0 | 0 | 0 | 1 | 0 | 1 | 0 |
| 0 | 0 | 0 | 0 | 0 | 0 | 0 | 0 | 0 | 0 | 0 | 0 | 0 | 0 |
| 0 | 0 | 0 | 0 | 0 | 0 | 0 | 0 | 0 | 0 | 0 | 0 | 0 | 0 |
| 704,855 | 715,163 | 572,485 | 2,583,586 | 1,486,451 | 3,636,322 | 3,962,279 | 5,114,489 | 5,855,167 | 8,172,660 | 9,227,264 | 3,451,076 | 5,881,573 | 9,377,136 |

| | | | | | | | | | | | | | |
|---|---|---|---|---|---|---|---|---|---|---|---|---|---|
| GM12878 | GM12878 | GM12878 | GM12878 | GM12878 | GM12878 | GM12878 | GM12878 | GM12878 | GM12878 | GM12878 | GM12878 | GM12878 | GM12878 |
| RAD21 | RAD21 | RAD21 | RAD21 | RAD21 | RAD21 | SMC1A | SMC1A | SMC1A | SMC1A | SMC1A | SMC1A | RNAPII | RNAPII |
| SHG0203 | SHG0205 | SHG0206 | SHG0207 | SHG0208 | SHG0209 | SHG0182 | SHG0186 | SHG0197 | SHG0200 | SHG0201 | SHG0204 | SHG0218 | SHG0219 |
| hg38 | hg38 | hg38 | hg38 | hg38 | hg38 | hg38 | hg38 | hg38 | hg38 | hg38 | hg38 | hg38 | hg38 |
| 0.06507599 | 0.04080005 | 0.05103529 | 0.0272089 | 0.02739122 | 0.02925437 | 0.03681377 | 0.05813272 | 0.04303481 | 0.02459852 | 0.02701237 | 0.03288814 | 0.02927647 | 0.02712616 |
| 191,895,737 | 130,723,974 | 162,624,186 | 62,793,728 | 68,451,827 | 79,128,174 | 88,752,922 | 147,848,455 | 149,720,829 | 58,803,883 | 57,751,339 | 177,247,438 | 57,701,716 | 57,662,440 |
| 181,580,861 | 123,784,145 | 153,072,139 | 59,211,581 | 64,680,345 | 74,251,010 | 86,265,518 | 139,345,536 | 144,916,276 | 56,903,448 | 54,925,050 | 173,467,548 | 47,703,440 | 47,582,400 |
| 174,244,222 | 118,072,681 | 145,811,190 | 56,587,584 | 61,714,453 | 70,710,360 | 83,689,009 | 133,177,422 | 140,949,358 | 55,412,659 | 52,788,350 | 169,298,401 | 44,780,648 | 44,727,208 |
| 142,866,662 | 97,255,562 | 119,664,193 | 46,028,720 | 49,078,393 | 55,920,819 | 70,352,994 | 103,749,435 | 118,986,672 | 46,604,769 | 42,174,020 | 147,103,752 | 31,953,228 | 32,058,056 |
| 138,316,876 | 93,862,760 | 115,339,163 | 44,521,320 | 47,427,796 | 53,963,819 | 68,703,503 | 100,205,515 | 116,115,293 | 45,681,378 | 40,943,087 | 144,187,673 | 30,478,266 | 30,510,648 |
| 137,637,355 | 93,454,012 | 114,829,001 | 44,331,077 | 47,222,452 | 53,726,175 | 67,918,010 | 98,814,463 | 114,538,527 | 44,911,203 | 40,483,176 | 142,558,840 | 30,299,353 | 30,347,279 |
| 137,637,355 | 93,454,012 | 114,829,001 | 44,331,077 | 47,222,452 | 53,726,175 | 67,918,010 | 98,814,463 | 114,538,527 | 44,911,203 | 40,483,176 | 142,558,840 | 30,299,353 | 30,347,279 |
| 2,800,485 | 2,618,896 | 2,798,679 | 2,204,312 | 2,095,451 | 2,178,202 | 1,914,369 | 2,532,457 | 2,298,015 | 1,825,199 | 2,076,666 | 2,446,515 | 2,029,433 | 2,016,268 |
| 2,769,385 | 2,586,966 | 2,764,032 | 2,180,309 | 2,075,332 | 2,156,250 | 1,897,061 | 2,507,478 | 2,278,233 | 1,803,937 | 2,053,586 | 2,419,741 | 1,999,990 | 1,990,892 |
| 2,593,084 | 2,424,384 | 2,586,355 | 2,049,577 | 1,954,728 | 2,024,006 | 1,799,976 | 2,368,735 | 2,147,125 | 1,681,152 | 1,914,466 | 2,283,339 | 1,780,041 | 1,793,040 |
| 2,593,084 | 2,424,384 | 2,586,355 | 2,049,577 | 1,954,728 | 2,024,006 | 1,799,976 | 2,368,735 | 2,147,125 | 1,681,152 | 1,914,466 | 2,283,339 | 1,780,041 | 1,793,040 |
| 801,145 | 722,533 | 802,303 | 511,059 | 442,773 | 489,157 | 286,266 | 681,109 | 527,349 | 461,177 | 489,631 | 651,260 | 423,117 | 386,400 |
| 1,791,939 | 1,701,851 | 1,784,052 | 1,538,518 | 1,511,955 | 1,534,849 | 1,513,710 | 1,687,626 | 1,619,776 | 1,219,975 | 1,424,835 | 1,632,079 | 1,356,924 | 1,406,640 |
| 13,357 | 10,142 | 13,514 | 4,943 | 4,025 | 4,763 | 2,303 | 12,606 | 6,264 | 4,466 | 4,987 | 7,318 | 9,443 | 8,927 |
| 1,778,582 | 1,691,709 | 1,770,538 | 1,533,575 | 1,507,930 | 1,530,086 | 1,511,407 | 1,675,020 | 1,613,512 | 1,215,509 | 1,419,848 | 1,624,761 | 1,347,481 | 1,397,713 |
| 11,244,395 | 8,058,925 | 8,778,655 | 4,373,713 | 4,527,189 | 4,803,930 | 6,605,440 | 7,001,774 | 9,000,582 | 4,229,117 | 3,867,039 | 12,680,709 | 1,339,159 | 1,393,601 |
| 10,719,823 | 10,725,236 | 10,900,905 | 9,312,672 | 9,306,439 | 9,463,101 | 9,998,098 | 10,146,056 | 10,501,513 | 7,495,830 | 8,484,451 | 10,047,768 | 5,874,233 | 6,132,695 |
| 22,778,720 | 19,516,836 | 20,495,377 | 14,202,387 | 14,280,426 | 14,760,951 | 16,892,107 | 17,841,545 | 20,035,708 | 12,190,590 | 12,846,108 | 23,387,055 | 7,645,952 | 7,921,623 |
| 11,520,968 | 11,447,769 | 11,703,208 | 9,823,731 | 9,749,212 | 9,952,258 | 10,284,364 | 10,827,165 | 11,028,862 | 7,957,007 | 8,974,082 | 10,699,028 | 6,297,350 | 6,519,095 |
| 5,750,646 | 4,784,628 | 5,044,682 | 3,031,457 | 3,108,659 | 3,253,199 | 4,131,383 | 4,219,834 | 5,086,835 | 2,771,706 | 2,689,145 | 6,011,352 | 1,082,922 | 1,133,230 |
| 2,867,529 | 1,965,856 | 2,176,272 | 921,848 | 965,732 | 1,040,253 | 1,552,406 | 1,671,613 | 2,243,498 | 959,255 | 805,678 | 3,266,537 | 203,713 | 210,465 |
| 1,365,339 | 780,487 | 904,026 | 281,993 | 301,119 | 334,272 | 569,593 | 651,896 | 951,453 | 325,462 | 243,671 | 1,677,510 | 43,385 | 43,139 |
| 645,462 | 311,856 | 375,594 | 90,150 | 97,829 | 112,114 | 212,440 | 259,580 | 402,540 | 111,611 | 78,484 | 842,856 | 10,920 | 10,375 |
| 309,354 | 127,301 | 159,676 | 31,110 | 33,625 | 39,664 | 82,078 | 108,115 | 174,149 | 39,484 | 27,927 | 424,815 | 3,477 | 2,979 |
| 151,595 | 53,681 | 69,720 | 11,428 | 12,661 | 15,136 | 32,493 | 47,852 | 77,277 | 14,561 | 11,182 | 217,430 | 1,560 | 1,064 |
| 75,924 | 23,233 | 31,503 | 4,724 | 5,308 | 6,318 | 13,358 | 22,503 | 35,604 | 5,738 | 5,238 | 113,577 | 864 | 526 |
| 39,534 | 10,665 | 14,590 | 2,172 | 2,317 | 2,949 | 5,914 | 11,339 | 16,578 | 2,460 | 2,843 | 59,815 | 485 | 238 |
| 21,228 | 4,983 | 6,946 | 1,171 | 1,232 | 1,479 | 2,888 | 6,179 | 7,996 | 1,140 | 1,741 | 32,040 | 328 | 150 |
| 11,468 | 2,449 | 3,518 | 674 | 741 | 871 | 1,513 | 3,683 | 3,979 | 596 | 1,280 | 17,544 | 218 | 111 |
| 6,457 | 1,371 | 1,855 | 422 | 443 | 524 | 827 | 2,359 | 2,198 | 342 | 832 | 9,580 | 155 | 65 |
| 3,863 | 718 | 1,011 | 278 | 334 | 364 | 529 | 1,598 | 1,211 | 222 | 561 | 5,454 | 139 | 45 |
| 2,372 | 434 | 612 | 217 | 200 | 284 | 389 | 1,227 | 754 | 151 | 520 | 3,048 | 103 | 40 |
| 4,324 | 823 | 1,191 | 562 | 573 | 741 | 949 | 3,225 | 1,475 | 435 | 1,429 | 4,496 | 245 | 73 |
| 1,770 | 405 | 669 | 291 | 329 | 352 | 629 | 2,066 | 797 | 268 | 946 | 1,237 | 78 | 25 |
| 496 | 106 | 183 | 82 | 71 | 96 | 189 | 658 | 257 | 74 | 307 | 368 | 9 | 2 |
| 179 | 32 | 57 | 46 | 24 | 45 | 78 | 273 | 111 | 38 | 124 | 163 | 1 | 0 |
| 200 | 38 | 57 | 29 | 14 | 31 | 80 | 338 | 120 | 39 | 107 | 187 | 0 | 1 |
| 12 | 1 | 7 | 2 | 1 | 1 | 7 | 37 | 14 | 1 | 11 | 17 | 0 | 0 |
| 0 | 0 | 0 | 0 | 2 | 0 | 0 | 5 | 0 | 0 | 0 | 1 | 0 | 0 |
| 0 | 0 | 0 | 0 | 0 | 0 | 0 | 0 | 0 | 0 | 0 | 0 | 0 | 0 |
| 0 | 0 | 0 | 0 | 0 | 0 | 0 | 0 | 0 | 0 | 0 | 0 | 0 | 0 |
| 11,257,752 | 8,069,067 | 8,792,169 | 4,378,656 | 4,531,214 | 4,808,693 | 6,607,743 | 7,014,380 | 9,006,846 | 4,233,583 | 3,872,026 | 12,688,027 | 1,348,602 | 1,402,528 |

| | | | | | | | | | | | | |
|---|---|---|---|---|---|---|---|---|---|---|---|---|
| GM12878 | GM12878 | GM12878 | GM12878 | GM12878 | GM12878 | GM12878 | GM12878 | GM12878 | GM12878 | GM12878 | GM12878 | GM12878 |
| RNAPII | RNAPII | RNAPII | RNAPII | RNAPII | RNAPII | RNAPII | RNAPII | RNAPII | RNAPII | RNAPII | RNAPII | RNAPII |
| SHG0234 | SHG0235 | SHG0238 | SHG0239 | SHG8034 | SHG8096 | SHG8097 | SHG8100 | SHG8101 | SHG8112 | SHG8116 | SHG8130 | SHG8131 |
| hg38 | hg38 | hg38 | hg38 | hg38 | hg38 | hg38 | hg38 | hg38 | hg38 | hg38 | hg38 | hg38 |
| 0.21233489 | 0.09380037 | 0.03209397 | 0.03319454 | 0.04271081 | 0.01937799 | 0.01926378 | 0.02247902 | 0.02052824 | 0.02017425 | 0.01793076 | 0.08709192 | 0.02821182 |
| 77,820,387 | 87,702,678 | 78,539,454 | 94,222,395 | 79,192,522 | 68,385,904 | 64,871,999 | 65,226,738 | 67,282,542 | 67,273,144 | 63,420,334 | 73,720,406 | 50,726,668 |
| 50,406,922 | 73,653,661 | 56,537,054 | 82,232,378 | 68,262,493 | 50,175,704 | 47,443,932 | 52,747,901 | 53,873,955 | 43,251,628 | 48,269,836 | 43,468,577 | 42,842,355 |
| 44,300,995 | 69,032,368 | 50,387,088 | 76,485,957 | 66,347,013 | 47,711,991 | 45,001,883 | 50,980,222 | 52,140,432 | 40,793,699 | 46,152,768 | 41,309,867 | 41,623,219 |
| 26,780,319 | 51,781,798 | 33,117,989 | 56,874,367 | 49,364,758 | 33,548,953 | 31,233,688 | 38,038,027 | 38,882,339 | 27,541,504 | 33,217,001 | 28,287,609 | 33,408,065 |
| 25,194,475 | 50,269,960 | 30,999,765 | 54,237,090 | 47,724,151 | 31,974,403 | 29,745,078 | 36,582,179 | 37,397,417 | 26,029,743 | 31,846,649 | 27,064,622 | 32,719,472 |
| 25,004,909 | 49,825,903 | 30,813,074 | 53,910,606 | 46,973,678 | 31,703,645 | 29,462,112 | 36,108,371 | 36,929,116 | 25,851,751 | 31,564,400 | 26,888,040 | 32,442,144 |
| 25,004,909 | 49,825,903 | 30,813,074 | 53,910,606 | 46,973,678 | 31,703,645 | 29,462,112 | 36,108,371 | 36,929,116 | 25,851,751 | 31,564,400 | 26,888,040 | 32,442,144 |
| 1,703,798 | 1,775,014 | 1,848,488 | 1,939,444 | 2,452,263 | 2,062,823 | 2,034,704 | 2,081,179 | 2,097,364 | 2,041,353 | 2,149,215 | 1,859,947 | 1,891,863 |
| 1,680,412 | 1,756,762 | 1,813,751 | 1,918,177 | 2,437,392 | 2,047,941 | 2,018,528 | 2,067,835 | 2,083,171 | 2,024,895 | 2,135,846 | 1,851,848 | 1,885,027 |
| 1,580,754 | 1,686,054 | 1,675,995 | 1,835,682 | 2,314,739 | 1,926,369 | 1,895,109 | 1,955,121 | 1,967,698 | 1,869,092 | 2,024,244 | 1,745,892 | 1,830,041 |
| 1,580,754 | 1,686,054 | 1,675,995 | 1,835,682 | 2,314,739 | 1,926,369 | 1,895,109 | 1,955,121 | 1,967,698 | 1,869,092 | 2,024,244 | 1,745,892 | 1,830,041 |
| 166,446 | 194,016 | 493,995 | 302,266 | 567,018 | 440,587 | 449,117 | 423,954 | 422,838 | 582,133 | 458,197 | 273,006 | 214,615 |
| 1,414,308 | 1,492,038 | 1,182,000 | 1,533,416 | 1,747,721 | 1,485,782 | 1,445,992 | 1,531,167 | 1,544,860 | 1,286,959 | 1,566,047 | 1,472,886 | 1,615,426 |
| 5,445 | 1,287 | 19,317 | 4,442 | 12,300 | 14,140 | 14,834 | 7,348 | 8,202 | 34,006 | 13,431 | 22,271 | 7,246 |
| 1,408,863 | 1,490,751 | 1,162,683 | 1,528,974 | 1,735,421 | 1,471,642 | 1,431,158 | 1,523,819 | 1,536,658 | 1,252,953 | 1,552,616 | 1,450,615 | 1,608,180 |
| 1,778,231 | 6,464,631 | 598,124 | 2,915,214 | 3,137,355 | 1,460,923 | 1,330,372 | 2,027,957 | 1,905,002 | 803,526 | 2,027,444 | 1,492,933 | 3,730,334 |
| 6,740,246 | 9,988,324 | 3,982,750 | 8,286,229 | 9,100,378 | 5,979,925 | 5,699,881 | 7,482,459 | 7,384,740 | 3,937,064 | 6,719,699 | 5,333,231 | 8,037,165 |
| 8,690,368 | 16,648,258 | 5,094,186 | 11,508,151 | 12,817,051 | 7,895,575 | 7,494,204 | 9,941,718 | 9,720,782 | 5,356,729 | 9,218,771 | 7,121,441 | 11,989,360 |
| 6,906,692 | 10,182,340 | 4,476,745 | 8,588,495 | 9,667,396 | 6,420,512 | 6,148,998 | 7,906,413 | 7,807,578 | 4,519,197 | 7,177,896 | 5,606,237 | 8,251,780 |
| 1,406,525 | 4,064,813 | 518,712 | 2,162,174 | 2,345,004 | 1,142,105 | 1,047,046 | 1,608,528 | 1,528,077 | 620,802 | 1,486,381 | 1,052,892 | 2,380,711 |
| 288,125 | 1,503,526 | 75,534 | 547,781 | 585,238 | 241,416 | 215,447 | 329,991 | 301,054 | 136,630 | 372,625 | 271,127 | 788,108 |
| 63,807 | 548,586 | 15,081 | 145,910 | 151,866 | 61,145 | 54,127 | 70,879 | 62,083 | 43,183 | 110,550 | 93,479 | 296,334 |
| 16,507 | 207,350 | 4,317 | 41,665 | 42,965 | 18,566 | 16,747 | 17,023 | 14,649 | 17,369 | 38,885 | 40,615 | 125,817 |
| 4,968 | 81,518 | 1,707 | 13,256 | 13,617 | 6,525 | 6,243 | 4,835 | 4,148 | 8,087 | 15,977 | 20,555 | 60,073 |
| 1,872 | 33,076 | 789 | 4,677 | 5,189 | 2,591 | 2,636 | 1,751 | 1,436 | 4,249 | 7,230 | 11,660 | 31,777 |
| 788 | 13,925 | 423 | 1,935 | 2,292 | 1,171 | 1,221 | 781 | 657 | 2,432 | 3,633 | 7,306 | 18,283 |
| 442 | 6,291 | 278 | 884 | 1,179 | 643 | 657 | 414 | 374 | 1,523 | 1,970 | 4,664 | 11,104 |
| 226 | 2,975 | 185 | 476 | 668 | 312 | 371 | 282 | 223 | 990 | 1,191 | 3,192 | 7,195 |
| 123 | 1,575 | 106 | 251 | 434 | 194 | 231 | 162 | 132 | 606 | 741 | 2,287 | 4,770 |
| 93 | 839 | 91 | 180 | 277 | 123 | 139 | 134 | 76 | 460 | 478 | 1,641 | 3,374 |
| 56 | 473 | 57 | 109 | 196 | 73 | 82 | 98 | 73 | 311 | 301 | 1,206 | 2,361 |
| 32 | 275 | 33 | 77 | 151 | 48 | 69 | 52 | 39 | 192 | 204 | 957 | 1,676 |
| 81 | 511 | 84 | 178 | 299 | 91 | 118 | 204 | 100 | 504 | 448 | 2,301 | 3,989 |
| 27 | 137 | 42 | 84 | 208 | 47 | 59 | 115 | 60 | 165 | 207 | 1,093 | 1,631 |
| 2 | 33 | 2 | 15 | 45 | 8 | 10 | 36 | 12 | 23 | 35 | 178 | 259 |
| 2 | 5 | 0 | 2 | 17 | 5 | 2 | 14 | 9 | 4 | 12 | 38 | 77 |
| 0 | 10 | 0 | 2 | 10 | 0 | 1 | 6 | 2 | 2 | 7 | 13 | 39 |
| 0 | 0 | 0 | 0 | 0 | 0 | 0 | 0 | 0 | 0 | 0 | 0 | 2 |
| 0 | 0 | 0 | 0 | 0 | 0 | 0 | 0 | 0 | 0 | 0 | 0 | 0 |
| 0 | 0 | 0 | 0 | 0 | 0 | 0 | 0 | 0 | 0 | 0 | 0 | 0 |
| 0 | 0 | 0 | 0 | 0 | 0 | 0 | 0 | 0 | 0 | 0 | 0 | 0 |
| 1,783,676 | 6,465,918 | 617,441 | 2,919,656 | 3,149,655 | 1,475,063 | 1,345,206 | 2,035,305 | 1,913,204 | 837,532 | 2,040,875 | 1,515,204 | 3,737,580 |

| Library_ID | LHG0052H | LHG0051H | LHG0104V | LHG0051H_0104V | LHG0035N_0035V | LHH0118V | LHH0127V |
|---|---|---|---|---|---|---|---|
| Library_type | hiseq | hiseq | novaseq | pooled | pooled | novaseq | novaseq |
| Reference_genome | hg38 | hg38 | hg38 | hg38 | hg38 | hg38 | hg38 |
| Cell_type | GM12878 | GM12878 | GM12878 | GM12878 | GM12878 | HCT116 | HCT116 |
| Factor | **CTCF** | **SMC1A** | **RAD21** | **Cohesin** | **RNAPII** | **CTCF** | **CTCF** |
| Total_read_pairs | 358,752,218 | 356152208 | 459,140,632 | 815,292,840 | 879,867,506 | 293,527,935 | 306,988,032 |
| Read_pairs_with_linker | 328,927,665 | 327803209 | 322,463,454 | 650,266,663 | 805,998,749 | 209,379,373 | 278,017,188 |
| Fraction_read_pairs_with_linl | 0.92 | 0.92 | 0.7 | 0.8 | 0.92 | 0.71 | 0.91 |
| One_tag | 91,116,895 | 92611052 | 121,729,821 | 214,340,873 | 305,223,593 | 67,711,510 | 90,202,553 |
| PET | 228,074,463 | 225677969 | 184,468,375 | 410,146,344 | 468,121,770 | 137,694,970 | 178,342,047 |
| Uniquely_mapped_PET | 179,600,076 | 179509164 | 147,434,776 | 326,943,940 | 359,389,378 | 101,683,183 | 139,475,184 |
| Non-redundant_PET | 106,110,023 | 112913401 | 67,917,076 | 180,798,216 | 58,636,573 | 14,691,093 | 58,699,205 |
| Redundancy | 0.41 | 0.37 | 0.54 | 0.45 | 0.84 | 0.86 | 0.58 |
| Non-redundant_tag | 362,869,381 | 362795441 | 379860551 | 736,311,882 | 269,106,487 | 86007544 | 214963074 |
| Peak | 28,992 | 19659 | 39,082 | 29,119 | 50,232 | 72,052 | 355,784 |
| Self-ligation_PET | 36,374,046 | 40192551 | 40,718,911 | 80,880,097 | 30,016,547 | 7,646,260 | 44,574,688 |
| Inter-ligation_PET | 69,735,977 | 72720850 | 27,198,165 | 99,918,119 | 28,620,026 | 7,044,833 | 14,124,517 |
| Intra-chr_PET | 53,879,279 | 57355817 | 24,463,289 | 81,818,234 | 22,888,857 | 5,371,336 | 12,558,160 |
| Inter-chr_PET | 15,856,698 | 15365033 | 2,734,876 | 18,099,885 | 5,731,169 | 1,673,497 | 1,566,357 |
| ratio_of_intra/inter_PET | 3.4 | 3.73 | 8.94 | 4.52 | 3.99 | 3.21 | 8.02 |
| Singleton | 54,239,711 | 57277746 | 20,413,204 | 72,885,936 | 19,742,331 | 5,489,032 | 11,124,088 |
| Intra-chr_singleton | 39,945,853 | 43361447 | 17,896,022 | 56,461,791 | 15,187,368 | 4,060,994 | 9,635,525 |
| Inter-chr_singleton | 14,293,858 | 13916299 | 2,517,182 | 16,424,145 | 4,554,963 | 1,428,038 | 1,488,563 |
| PET_cluster | 5,231,012 | 5487852 | 2,055,333 | 7,913,116 | 2,840,108 | 590,202 | 920,945 |
| ratio_of_intra/inter_cluster | 6.25 | 7.02 | 19.32 | 9.02 | 5.15 | 4.62 | 24.8 |
| Intra-chr_PET_cluster | 4,509,611 | 4803284 | 1,954,185 | 7,123,625 | 2,378,076 | 485,091 | 885,250 |
| pets_number_2 | 3,328,184 | 3639546 | 1,401,063 | 5,104,152 | 1,581,827 | 358,928 | 608,866 |
| pets_number_3 | 653,815 | 669744 | 282,716 | 1,057,262 | 395,541 | 68,675 | 133,453 |
| pets_number_4 | 215,205 | 209719 | 101,668 | 367,194 | 159,917 | 24,637 | 51,355 |
| pets_number_5 | 95,704 | 90839 | 47,834 | 171,135 | 81,272 | 11,965 | 26,092 |
| pets_number_6 | 51,669 | 48183 | 26,767 | 96,803 | 47,435 | 6,490 | 15,256 |
| pets_number_7 | 32,366 | 29546 | 17,124 | 60,673 | 30,227 | 3,797 | 10,002 |
| pets_number_8 | 21,928 | 20090 | 11,760 | 41,862 | 19,361 | 2,524 | 6,993 |
| pets_number_9 | 15,747 | 14331 | 8,686 | 30,342 | 12,970 | 1,655 | 5,190 |
| pets_number_10 | 12,021 | 10657 | 6,672 | 23,162 | 8,993 | 1,275 | 3,921 |
| pets_number>10 | 82,972 | 70629 | 49,895 | 171,040 | 40,533 | 5,145 | 24,122 |
| Inter-chr_PET_cluster | 721,401 | 684568 | 101,148 | 789,491 | 462,032 | 105,111 | 35,695 |
| pets_number_2 | 631,456 | 618023 | 89,814 | 710,643 | 338,328 | 86,076 | 31,230 |
| pets_number_3 | 70,736 | 56398 | 8,625 | 65,653 | 68,613 | 11,466 | 3,275 |
| pets_number_4 | 13,491 | 8181 | 1,956 | 10,365 | 25,405 | 3,897 | 788 |
| pets_number_5 | 3,492 | 1498 | 528 | 2,083 | 12,313 | 1,662 | 254 |
| pets_number_6 | 1,165 | 316 | 153 | 502 | 6,815 | 852 | 97 |
| pets_number_7 | 447 | 92 | 39 | 140 | 4,210 | 464 | 19 |
| pets_number_8 | 221 | 30 | 11 | 52 | 2,549 | 272 | 14 |
| pets_number_9 | 169 | 7 | 6 | 12 | 1,541 | 146 | 5 |
| pets_number_10 | 81 | 8 | 3 | 8 | 970 | 115 | 2 |
| pets_number>10 | 143 | 15 | 13 | 33 | 1,288 | 161 | 11 |

| | | | | | | | |
|---|---|---|---|---|---|---|---|
| intra_pets_number>2 | 1,181,427 | 1,163,738 | 553,122 | 2,019,473 | 796,249 | 126,163 | 276,384 |

| LHH0118V_0 | LHH0098V | LHH0128V | LHH0098V_0 | LHH0139 | LHH0139V | LHH0140 | LHH0140V | LHH0139_01 | LHH0146V | LHH0147V |
|---|---|---|---|---|---|---|---|---|---|---|
| pooled | novaseq | novaseq | pooled | qcseq | hiseq | qcseq | hiseq | pooled | novaseq | novaseq |
| hg38 | hg38 | hg38 | hg38 | hg38 | hg38 | hg38 | hg38 | hg38 | hg38 | hg38 |
| HCT116-wt | HCT116 | HCT116 | HCT116-wt | HCT116-RAD2 | HCT116-wt | HCT116-RAD2 | HCT116-wt | HCT116-wt | HCT116-mAC | HCT116-mAC |
| **CTCF** | **RNAPII** | **RNAPII** | **RNAPII** | **RAD21** | **RAD21** | **RAD21** | **RAD21** | **RAD21** | **none-likehic** | **none-likehic** |
| 600,515,967 | 517207217 | 316,087,217 | 833,294,434 | 48,523,665 | 90,562,053 | 42,973,485 | 110,921,989 | 292,981,192 | 360,397,636 | 372,697,334 |
| 487,396,561 | 451545618 | 272,698,811 | 724,244,429 | 40,331,721 | 75,028,257 | 36,548,964 | 93,886,130 | 245,795,072 | 332,763,506 | 328,476,305 |
| 0.81 | 0.87 | 0.86 | 0.87 | 0.83 | 0.83 | 0.85 | 0.85 | 0.84 | 0.92 | 0.88 |
| 157,914,063 | 173652631 | 78,399,895 | 252,052,526 | 11,591,968 | 21,789,368 | 10,344,168 | 26,846,543 | 70,572,047 | 102,735,699 | 88,151,514 |
| 316,037,017 | 270237912 | 184,446,325 | 454,684,237 | 27,411,989 | 51,330,793 | 25,034,264 | 64,739,897 | 168,516,943 | 221,734,530 | 231,282,572 |
| 241,158,367 | 177441741 | 148,915,219 | 326,356,960 | 23,001,256 | 43,118,396 | 20,973,285 | 54,221,758 | 141,314,695 | 175,237,861 | 185,042,954 |
| 73,213,214 | 38443824 | 44,042,591 | 82,471,851 | 6,705,673 | 8,345,186 | 7,585,990 | 10,991,359 | 21,572,799 | 106,924,059 | 110,480,076 |
| 0.7 | 0.78 | 0.7 | 0.75 | 0.71 | 0.81 | 0.64 | 0.8 | 0.85 | 0.39 | 0.4 |
| 297905125 | 189630121 | 164602051 | 351858151 | 25860590 | 33247217 | 27716144 | 42054236 | 85527138 | 355115179 | 359906998 |
| 117,603 | 598505 | 390,734 | 117,768 | 51,228 | 59,697 | 49,305 | 64,637 | 87,011 | 177,907 | 141,079 |
| 52,092,777 | 20140372 | 34,897,451 | 55,023,272 | 5,327,600 | 6,615,282 | 6,056,074 | 8,734,119 | 17,099,822 | 70,678,761 | 68,671,890 |
| 21,120,437 | 18303452 | 9,145,140 | 27,448,579 | 1,378,073 | 1,729,904 | 1,529,916 | 2,257,240 | 4,472,977 | 36,245,298 | 41,808,186 |
| 17,889,612 | 14357432 | 7,733,172 | 22,090,597 | 1,248,156 | 1,565,122 | 1,415,930 | 2,080,900 | 4,080,367 | 29,812,795 | 32,436,743 |
| 3,230,825 | 3946020 | 1,411,968 | 5,357,982 | 129,917 | 164,782 | 113,986 | 176,340 | 392,610 | 6,432,503 | 9,371,443 |
| 5.54 | 3.64 | 5.48 | 4.12 | 9.61 | 9.5 | 12.42 | 11.8 | 10.39 | 4.63 | 3.46 |
| 15,771,334 | 15085865 | 7,766,436 | 22,053,300 | 1,167,944 | 1,351,801 | 1,345,956 | 1,792,701 | 3,282,048 | 31,965,122 | 37,613,583 |
| 12,864,812 | 11630363 | 6,454,446 | 17,286,529 | 1,048,582 | 1,210,361 | 1,239,282 | 1,640,644 | 2,953,267 | 25,735,750 | 28,607,251 |
| 2,906,522 | 3455502 | 1,311,990 | 4,766,771 | 119,362 | 141,440 | 106,674 | 152,057 | 328,781 | 6,229,372 | 9,006,332 |
| 1,617,543 | 1335842 | 506,966 | 2,024,843 | 78,635 | 141,066 | 70,779 | 176,097 | 413,910 | 1,699,287 | 1,721,093 |
| 10.46 | 5.11 | 10.42 | 6.69 | 15.86 | 13.1 | 20.55 | 15.97 | 14.65 | 16.46 | 8.82 |
| 1,476,339 | 1117059 | 462,584 | 1,761,433 | 73,972 | 131,060 | 67,495 | 165,719 | 387,454 | 1,601,936 | 1,545,821 |
| 999,042 | 889107 | 349,334 | 1,331,315 | 55,739 | 97,185 | 52,311 | 124,057 | 274,784 | 1,250,091 | 1,236,323 |
| 227,025 | 142431 | 63,563 | 245,926 | 10,140 | 18,636 | 8,945 | 23,394 | 58,207 | 215,147 | 194,839 |
| 89,437 | 43807 | 21,989 | 83,219 | 3,661 | 6,672 | 2,967 | 8,172 | 22,254 | 67,625 | 58,155 |
| 45,656 | 18485 | 9,938 | 37,137 | 1,618 | 3,122 | 1,251 | 3,920 | 10,995 | 28,242 | 23,620 |
| 27,210 | 8805 | 5,225 | 19,304 | 840 | 1,773 | 628 | 2,098 | 6,301 | 14,348 | 11,554 |
| 17,424 | 4724 | 3,016 | 11,074 | 525 | 1,061 | 389 | 1,280 | 3,848 | 8,127 | 6,518 |
| 12,246 | 2752 | 1,965 | 6,987 | 308 | 625 | 250 | 767 | 2,475 | 4,835 | 3,888 |
| 9,012 | 1668 | 1,382 | 4,793 | 234 | 417 | 145 | 479 | 1,778 | 3,279 | 2,621 |
| 6,896 | 1097 | 969 | 3,424 | 158 | 302 | 111 | 328 | 1,236 | 2,275 | 1,826 |
| 42,391 | 4183 | 5,203 | 18,254 | 749 | 1,267 | 498 | 1,224 | 5,576 | 7,967 | 6,477 |
| 141,204 | 218783 | 44,382 | 263,410 | 4,663 | 10,006 | 3,284 | 10,378 | 26,456 | 97,351 | 175,272 |
| 117,616 | 185576 | 37,140 | 222,896 | 3,842 | 8,046 | 2,765 | 8,334 | 20,637 | 90,766 | 163,464 |
| 14,795 | 22164 | 4,910 | 27,108 | 550 | 1,211 | 375 | 1,295 | 3,407 | 5,511 | 9,940 |
| 4,696 | 6386 | 1,464 | 7,866 | 180 | 402 | 96 | 396 | 1,179 | 810 | 1,438 |
| 1,923 | 2557 | 493 | 3,050 | 60 | 198 | 31 | 171 | 579 | 167 | 294 |
| 949 | 1107 | 207 | 1,321 | 22 | 85 | 9 | 93 | 314 | 51 | 74 |
| 492 | 525 | 96 | 621 | 6 | 36 | 5 | 36 | 148 | 15 | 26 |
| 288 | 238 | 29 | 272 | 2 | 15 | 1 | 29 | 79 | 5 | 10 |
| 152 | 112 | 18 | 130 | 0 | 3 | 0 | 12 | 51 | 6 | 4 |
| 120 | 61 | 13 | 73 | 0 | 4 | 1 | 6 | 26 | 4 | 3 |
| 173 | 57 | 12 | 73 | 1 | 6 | 1 | 6 | 36 | 16 | 19 |

| | | | | | | | | | | |
|---|---|---|---|---|---|---|---|---|---|---|
| 477,297 | 227,952 | 113,250 | 430,118 | 18,233 | 33,875 | 15,184 | 41,662 | 112,670 | 351,845 | 309,498 |

| | | | | | | | | | | |
|---|---|---|---|---|---|---|---|---|---|---|
| LHH0157 | LHH0172 | LHH0157_01 | LHH0158 | LHH0173 | LHH0158_01 | LHH0170 | LHH0159 | LHH0159V | LHH0170_0159 | LHH0171 |
| novaseq | novaseq | pooled | novaseq | novaseq | pooled | novaseq | novaseq | novaseq | pooled | novaseq |
| hg38 | hg38 | hg38 | hg38 | hg38 | hg38 | hg38 | hg38 | hg38 | hg38 | hg38 |
| HCT116-RAD2 | HCT116-RAD2 | HCT116-RAD2 | HCT116-RAD2 | HCT116-RAD2 | HCT116-RAD2 | HCT116-RAD2 | HCT116-RAD2 | HCT116-RAD2 | HCT116-RAD21 | HCT116-RAD2 |
| **CTCF** | **CTCF** | **CTCF** | **CTCF** | **CTCF** | **CTCF** | **RNAPII** | **RNAPII** | **RNAPII** | **RNAPII** | **RNAPII** |
| 366,508,457 | 569,205,831 | 935,714,288 | 401,699,562 | 460,564,094 | 862,263,656 | 557,061,917 | 343,180,214 | 483,901,014 | 1,384,143,145 | 565,251,115 |
| 339,696,732 | 527,351,506 | 867,048,238 | 371,666,476 | 425,780,166 | 797,446,642 | 517,998,624 | 320,331,706 | 453,410,923 | 1,291,741,253 | 523,472,548 |
| 0.93 | 0.93 | 0.93 | 0.93 | 0.92 | 0.92 | 0.93 | 0.93 | 0.94 | 0.93 | 0.93 |
| 107,052,736 | 175,480,282 | 282,533,018 | 115,962,944 | 142,223,489 | 258,186,433 | 197,218,539 | 107,125,259 | 152,138,436 | 456,482,234 | 193,704,777 |
| 212,107,094 | 342,780,972 | 554,888,066 | 232,472,432 | 275,740,089 | 508,212,521 | 316,451,293 | 192,719,614 | 269,464,891 | 778,635,798 | 323,895,885 |
| 171,149,739 | 272,626,140 | 443,775,879 | 187,436,945 | 220,538,751 | 407,975,696 | 247,521,404 | 152,353,109 | 213,103,487 | 612,978,000 | 255,007,981 |
| 43,952,826 | 78,684,532 | 122,544,001 | 46,417,943 | 51,541,251 | 97,870,444 | 34,436,532 | 36,654,383 | 39,516,824 | 80,322,705 | 51,158,533 |
| 0.74 | 0.71 | 0.72 | 0.75 | 0.77 | 0.76 | 0.86 | 0.76 | 0.81 | 0.87 | 0.8 |
| 162587502 | 292882828 | 449256959 | 172911962 | 203452802 | 371635392 | 161925339 | 146107485 | 161556435 | 350483376 | 220694073 |
| 110,176 | 127,600 | 131,371 | 106,552 | 122,255 | 127,824 | 109,499 | 108,411 | 112,496 | 129,576 | 122,089 |
| 26,184,726 | 41,874,843 | 67,966,716 | 30,300,253 | 29,878,151 | 60,089,841 | 21,048,717 | 23,672,339 | 25,521,178 | 50,582,246 | 33,260,249 |
| 17,768,100 | 36,809,689 | 54,577,285 | 16,117,690 | 21,663,100 | 37,780,603 | 13,387,815 | 12,982,044 | 13,995,646 | 29,740,459 | 17,898,284 |
| 14,623,852 | 26,969,581 | 41,593,019 | 11,722,636 | 14,413,758 | 26,136,274 | 9,377,270 | 10,254,888 | 11,040,356 | 22,232,867 | 11,803,079 |
| 3,144,248 | 9,840,108 | 12,984,266 | 4,395,054 | 7,249,342 | 11,644,329 | 4,010,545 | 2,727,156 | 2,955,290 | 7,507,592 | 6,095,205 |
| 4.65 | 2.74 | 3.2 | 2.67 | 1.99 | 2.24 | 2.34 | 3.76 | 3.74 | 2.96 | 1.94 |
| 13,028,489 | 29,912,273 | 40,805,430 | 12,732,023 | 18,289,936 | 30,050,380 | 10,487,081 | 10,006,862 | 10,745,264 | 21,057,467 | 15,085,696 |
| 10,266,628 | 20,861,230 | 28,997,270 | 8,876,478 | 11,802,614 | 19,712,597 | 7,101,928 | 7,670,360 | 8,213,675 | 14,921,577 | 9,694,807 |
| 2,761,861 | 9,051,043 | 11,808,160 | 3,855,545 | 6,487,322 | 10,337,783 | 3,385,153 | 2,336,502 | 2,531,589 | 6,135,890 | 5,390,889 |
| 1,526,800 | 2,348,011 | 4,063,337 | 1,214,251 | 1,328,864 | 2,659,834 | 1,054,709 | 1,079,685 | 1,130,837 | 2,845,881 | 1,104,285 |
| 7.93 | 5.74 | 6.8 | 4.08 | 2.97 | 3.62 | 3.17 | 5.3 | 5.37 | 4.14 | 2.67 |
| 1,355,830 | 1,999,617 | 3,542,226 | 975,006 | 993,955 | 2,083,725 | 801,892 | 908,331 | 953,336 | 2,291,727 | 803,627 |
| 939,893 | 1,419,871 | 2,393,958 | 701,185 | 748,694 | 1,485,805 | 571,362 | 674,246 | 678,869 | 1,574,241 | 607,243 |
| 205,317 | 296,536 | 548,079 | 140,485 | 137,258 | 303,202 | 117,299 | 129,304 | 144,179 | 355,364 | 110,076 |
| 79,815 | 110,861 | 216,026 | 52,970 | 48,897 | 114,356 | 46,394 | 45,721 | 55,700 | 141,451 | 39,463 |
| 39,519 | 53,730 | 108,662 | 25,907 | 22,476 | 55,774 | 23,271 | 21,328 | 26,657 | 71,848 | 18,680 |
| 22,387 | 30,405 | 63,309 | 14,353 | 11,974 | 31,008 | 13,472 | 11,246 | 14,680 | 41,900 | 9,815 |
| 14,351 | 19,054 | 40,717 | 9,012 | 6,860 | 19,295 | 8,361 | 6,538 | 8,746 | 26,420 | 5,441 |
| 9,649 | 12,916 | 27,975 | 6,055 | 4,258 | 13,071 | 5,556 | 3,993 | 5,237 | 17,439 | 3,289 |
| 6,754 | 9,102 | 20,376 | 4,220 | 2,874 | 9,262 | 3,703 | 2,672 | 3,510 | 11,951 | 2,036 |
| 5,274 | 6,780 | 15,349 | 3,169 | 1,938 | 6,997 | 2,614 | 1,848 | 2,409 | 8,495 | 1,444 |
| 32,871 | 40,362 | 107,775 | 17,650 | 8,726 | 44,955 | 9,860 | 11,435 | 13,349 | 42,618 | 6,140 |
| 170,970 | 348,394 | 521,111 | 239,245 | 334,909 | 576,109 | 252,817 | 171,354 | 177,501 | 554,154 | 300,658 |
| 144,115 | 292,172 | 437,460 | 199,792 | 277,398 | 478,560 | 194,951 | 141,177 | 138,782 | 424,615 | 241,669 |
| 18,596 | 36,346 | 55,287 | 26,454 | 37,378 | 64,177 | 31,704 | 19,950 | 23,105 | 71,833 | 35,975 |
| 5,104 | 11,409 | 16,646 | 7,952 | 11,918 | 20,014 | 11,978 | 5,994 | 8,442 | 26,757 | 12,341 |
| 1,924 | 4,504 | 6,485 | 2,990 | 4,626 | 7,656 | 5,856 | 2,421 | 3,636 | 13,133 | 5,427 |
| 745 | 2,057 | 2,817 | 1,244 | 2,036 | 3,309 | 3,214 | 1,023 | 1,748 | 7,130 | 2,687 |
| 278 | 1,012 | 1,296 | 461 | 891 | 1,368 | 1,963 | 443 | 902 | 4,180 | 1,305 |
| 116 | 448 | 571 | 217 | 377 | 600 | 1,202 | 200 | 454 | 2,511 | 629 |
| 53 | 217 | 273 | 89 | 161 | 251 | 727 | 72 | 218 | 1,593 | 321 |
| 23 | 115 | 139 | 28 | 65 | 94 | 481 | 45 | 113 | 977 | 151 |
| 16 | 114 | 137 | 18 | 59 | 80 | 741 | 29 | 101 | 1,425 | 153 |

| | | | | | | | | | | |
|---|---|---|---|---|---|---|---|---|---|---|
| 415,937 | 579,746 | 1,148,268 | 273,821 | 245,261 | 597,920 | 230,530 | 234,085 | 274,467 | 717,486 | 196,384 |

| | | | | | | |
|---|---|---|---|---|---|---|
| LHH0160 | LHH0160V | LHH0171_0160_ | LHES0007H | LHK0009H_0 | LHH0123_01 | LHM0011_0011H_0014_0014H |
| novaseq | novaseq | pooled | hiseq | pooled | pooled | pooled |
| hg38 | hg38 | hg38 | hg38 | hg38 | hg38 | hg38 |
| HCT116-RAD | HCT116-RAD | HCT116-RAD21- | H1 | K562 | HepG2 | MCF7 |
| **RNAPII** | **RNAPII** | **RNAPII** | **RNAPII** | **RNAPII** | **RNAPII** | **RNAPII** |
| 363,275,787 | 533,711,372 | 1,462,238,274 | 302061532 | 914,524,429 | 743,739,702 | 716,354,301 |
| 342,116,735 | 503,748,354 | 1,369,337,637 | 269758389 | 739,766,316 | 622,275,298 | 643,504,683 |
| 0.94 | 0.94 | 0.94 | 0.89 | 0.81 | 0.84 | 0.9 |
| 125,284,102 | 184,849,728 | 503,838,607 | 57451567 | 201,703,349 | 200,525,038 | 178,125,576 |
| 194,175,452 | 282,276,316 | 800,347,653 | 210129518 | 519,892,314 | 374,836,539 | 444,009,576 |
| 150,912,198 | 219,341,332 | 625,261,511 | 171470262 | 391,929,409 | 295,625,450 | 348,697,395 |
| 38,995,470 | 42,335,354 | 99,886,632 | 74426727 | 156,295,365 | 65,026,411 | 154,826,157 |
| 0.74 | 0.81 | 0.84 | 0.57 | 0.6 | 0.78 | 0.56 |
| 162250996 | 181560853 | 429169409 | 234348256 | 582615618 | 276774478 | 529025826 |
| 104,830 | 107,036 | 128,211 | 28333 | 21,757 | 48,741 | 20,673 |
| 28,505,332 | 30,964,279 | 68,778,048 | 42233786 | 84,392,579 | 48,372,785 | 87,161,316 |
| 10,490,138 | 11,371,075 | 31,108,584 | 32192941 | 71,902,786 | 16,653,626 | 67,664,841 |
| 7,335,665 | 7,938,284 | 20,986,880 | 26596494 | 44,670,542 | 13,101,973 | 51,270,878 |
| 3,154,473 | 3,432,791 | 10,121,704 | 5596447 | 27,232,244 | 3,551,653 | 16,393,963 |
| 2.33 | 2.31 | 2.07 | 4.75 | 1.64 | 3.69 | 3.13 |
| 8,330,886 | 8,944,532 | 23,954,192 | 25130789 | 53,688,182 | 10,525,832 | 41,130,149 |
| 5,600,175 | 5,992,254 | 15,370,358 | 20114319 | 31,207,039 | 7,425,655 | 29,048,919 |
| 2,730,711 | 2,952,278 | 8,583,834 | 5016470 | 22,481,143 | 3,100,177 | 12,081,230 |
| 788,470 | 848,627 | 2,468,116 | 2542710 | 6,657,361 | 1,363,984 | 8,765,510 |
| 3.22 | 3.21 | 2.87 | 8.39 | 2.15 | 6.48 | 3.77 |
| 601,727 | 646,960 | 1,830,733 | 2271974 | 4,546,668 | 1,181,743 | 6,928,170 |
| 447,022 | 460,321 | 1,296,916 | 1740365 | 3,259,054 | 715,177 | 4,747,398 |
| 83,730 | 95,704 | 272,754 | 316246 | 749,303 | 190,897 | 1,251,428 |
| 29,782 | 37,706 | 105,231 | 98702 | 248,839 | 85,424 | 434,829 |
| 13,881 | 18,228 | 52,271 | 41074 | 106,124 | 47,751 | 184,866 |
| 7,387 | 9,904 | 29,352 | 20724 | 53,767 | 29,959 | 92,042 |
| 4,446 | 5,976 | 18,010 | 11989 | 31,609 | 20,048 | 52,045 |
| 2,868 | 3,820 | 11,678 | 7962 | 19,486 | 14,150 | 32,134 |
| 1,971 | 2,580 | 7,834 | 5618 | 13,416 | 10,507 | 21,810 |
| 1,473 | 1,848 | 5,566 | 4046 | 9,735 | 8,000 | 15,658 |
| 9,167 | 10,873 | 31,121 | 25248 | 55,335 | 59,830 | 95,960 |
| 186,743 | 201,667 | 637,383 | 36947 | 2,110,693 | 182,241 | 1,837,340 |
| 154,715 | 157,999 | 498,278 | 270736 | 1,725,833 | 139,794 | 1,429,651 |
| 21,426 | 26,041 | 80,906 | 240706 | 292,527 | 22,645 | 291,013 |
| 6,288 | 9,494 | 28,921 | 23648 | 65,042 | 9,261 | 76,835 |
| 2,514 | 4,259 | 13,457 | 4828 | 17,418 | 4,635 | 24,538 |
| 1,059 | 2,044 | 7,092 | 1200 | 5,539 | 2,540 | 8,711 |
| 416 | 969 | 3,906 | 260 | 2,111 | 1,414 | 3,375 |
| 181 | 422 | 2,094 | 71 | 940 | 826 | 1,443 |
| 70 | 215 | 1,190 | 15 | 459 | 479 | 598 |
| 32 | 110 | 678 | 5 | 216 | 282 | 319 |
| 42 | 114 | 861 | 1 | 608 | 365 | 857 |
| 154,705 | 186,639 | 533,817 | 531,609 | 1,287,614 | 466,566 | 2,180,772 |
